# Quantitative metaproteomics of patient fecal microbiota identifies host and microbial proteins associated with ulcerative colitis

**DOI:** 10.1101/2020.11.11.378836

**Authors:** Peter S. Thuy-Boun, Ana Y. Wang, Ana Crissien-Martinez, Janice H. Xu, Sandip Chatterjee, Gregory S. Stupp, Andrew I. Su, Walter J. Coyle, Dennis W. Wolan

## Abstract

Mass spectrometry-based metaproteomics technologies enable the direct observation of proteins within complex multi-organism environments. A major hurdle in mapping metaproteomic fragmentation spectra to their corresponding peptides is the need for large peptide databases encompassing all anticipated species contained within a biological sample. As we cannot predict the taxonomic composition of microbiomes *a priori*, we developed the ComPIL database which contains a comprehensive collection of 4.8 billion unique peptides from public sequencing repositories to enable our proteomics analyses. We analyzed fecal samples from ulcerative colitis (UC) patients using a tandem mass spectrometry (LC-MS/MS) workflow coupled to ComPIL in search of aberrant UC-associated proteins. We found 176 host and microbial protein groups differentially enriched between the healthy (control) or UC volunteer groups. Notably, gene ontology (GO) enrichment analysis revealed that serine-type endopeptidases are overrepresented in UC compared to healthy volunteers. Additionally, we demonstrate the feasibility of serine hydrolase chemical enrichment from fecal samples using a biotinylated fluorophosphate (FP) probe. Our findings illustrate that probe-susceptible hydrolases from hosts and microbes are likely active in the distal gut. Finally, we applied *de novo* peptide sequencing methods to our metaproteomics data to estimate the size of the “dark peptidome,” the complement of peptides unidentified using ComPIL. We posit that our metaproteomics methods are generally applicable to future microbiota analyses and that our list of FP probe-enriched hydrolases may represent an important functionality to understanding the etiology of UC.

## Introduction

Inflammatory bowel disease (IBD) is a chronic medical condition characterized by relapsing inflammation of the gastrointestinal (GI) tract. This disease is broadly divisible into two categories based on where inflammation occurs. In ulcerative colitis (UC), inflammation is restricted to the large intestine, while in Crohn’s disease (CD), inflammation can occur anywhere along the GI tract (1, 2). In addition to reduced life expectancy, IBD patients can suffer dramatic quality-of-life reductions and are at increased risk to develop gastrointestinal tract malignancy (3). The incidence and prevalence of IBD in developed countries has steadily increased in the last few decades, making this disease a public health concern with a potentially heavy cost burden due to a requirement for long-term management (4). Targeted cures for UC and CD are highly desirable, but the search for such treatments is hampered by our incomplete understanding of disease development. Genome wide association studies (GWAS) have identified over 200 genetic loci associated with UC and CD, but the polygenic nature of these conditions explains only a minor portion of disease incidence (5–7). Concordance rates of about 30% for CD and 15% for UC among monozygotic twins suggests a significant non-genetic contribution to disease development (8). Because our gut microbes are in perpetual contact with our GI tracts, they comprise important but ill-defined environmental variables that many studies have implicated in IBD development. IBD triggers are unknown, but its progression is hypothesized to be amplified by inappropriate host-microbe interactions that lead to dysbiosis and, eventually, observable gross pathology (9–13).

Efforts to identify potential microbial drivers of IBD are ongoing but stymied by the immense taxonomic complexity of the gut microbiota. The gut harbors hundreds of distinct species per individual that can change over time and with perturbations to host lifestyle or xenobiotic exposure (14–16). Campaigns to characterize and monitor the gut microbiota frequently utilize metagenomic sequencing technologies, which can provide information about microbial community structure, genetic potential, and transcriptional activities (17,18). As the collective size of shotgun metagenomic sequence space has expanded from these efforts, so too have the opportunities for liquid chromatography tandem mass spectrometry (LC-MS/MS)-based metaproteomics (19), which rely on protein reference databases constructed from translated genome sequences (20). Host protein profiling is a straight-forward process given the relative completeness of host (*e.g.*, human, mouse, *etc*.) genome assemblies, but microbiota protein profiling is more difficult due to the high strain diversity and presence of unculturable microbes in the gut (21). Reference database curation has proven to be an important consideration in metaproteomics but becomes computationally burdensome as community diversity expands (22). To address this problem, we developed the Comprehensive Protein Identification Library (ComPIL), a large and scalable proteomics database generally intended for metaproteomics studies (23). In its current iteration (ComPIL 2.0), it houses >4.8 billion unique, tryptic peptides derived from >113 million bacterial, archeal, viral, and eukaryotic parent protein sequences assembled from public sequencing repositories (24). With periodic incorporation of new sequences from shotgun metagenomics repositories, we envision that ComPIL will help enable interlaboratory consonance in the global interpretation and communication of bottom-up metaproteomics results. In addition to enabling the direct, large-scale observation of proteins in a complex mixture, LC-MS/MS-based metaproteomics techniques obviate a requirement for intact cells, facilitate the observation of post-translational protein modifications, and enable functional interrogation of new or incompletely annotated proteins through such cognate techniques as activity- and affinity-based protein profiling (ABPP) (25,26).

Relative to metagenomics, LC-MS/MS-based metaproteomics are less commonly applied and more rarely employed in IBD studies. In fact, the first large-scale endeavor to identify proteins from a microbial biofilm community was only disclosed by Banfield, *et al.* in 2005 (27–29). In 2009, Jansson, *et al*. leveraged high-resolution LC-MS/MS to demonstrate the viability of large-scale metaproteomics in fecal samples collected from a twin pair (30). The aforementioned study demonstrated for the first time that bottom-up LC-MS/MS-based proteomics technology is suitable for such a complex environment and that it could generate a model of the gut microbiota that is orthogonal to that produced by metagenomics. Since then, several groups have deployed bottom-up metaproteomics to investigate the etiology of IBD in humans by examining patient fecal extracts, intestinal biopsy tissue, and/or blood samples (31–40). Additionally, several groups including our own have paired traditional proteomic profiling with ABPP techniques in microbiota samples to detect and annotate key protein functionalities often undetectable without pre-enrichment (41–44).

The insights garnered from previous metaproteomics studies are valuable for forming a consensus about the constellation of IBD-related environmental factors. We aim to contribute to this nascent pool by presenting a combined 16S amplicon sequencing and metaproteomics analysis of fecal samples from healthy volunteers and ulcerative colitis patients to identify novel proteins associated with health or disease. Using a pipeline that incorporates a novel, strong-acid sample preparation procedure (45), label-free high resolution LC-MS/MS, and the ComPIL database coupled to the ProLuCID/SEQUEST search engine (20, 46, 47), we identify 176 protein groups and several gene ontology (GO) terms enriched in either cohort (48). We show that proteomics can provide a more complete picture of gut microbiota taxonomy that includes host, microbial, and even dietary proteins. Using ABPP, we demonstrate that not only are microbiota proteins enzymatically active after collection, but additional proteomic depth can be achieved using ABPP enrichment strategies. Finally, using *de novo* peptide sequencing tools, we provide a means for estimating the size of database-elusive peptide space in our LC-MS/MS data, enabling a rough estimation of metaproteome completeness. This measure can help shape future decision-making processes regarding the need for additional shotgun metagenomic work to support a given metaproteomics study.

## Results

### Demographics of healthy control and ulcerative colitis volunteers

Healthy (n = 8, M/F ratio = 5:3, mean age = 44 years) and UC (n = 10, M/F ratio = 8:2, mean age = 46 years) patient volunteers consisted of a mixture of males and females between the ages of 21-76 years of age (global mean age = 45 years, global median age = 39 years) at the time of enrollment. UC patients presented with mild to severe symptoms and a range of Mayo scores (0-9) during enrollment. Individuals with BMI values > 60, as well as any recent antibiotics usage (< 3 months prior to sample collection), severe diarrheal illnesses, or *Clostridium difficile* infections were excluded.

### Metaproteomics yields a more comprehensive taxonomic diversity than 16S rDNA sequencing

At the kingdom level, LC-MS/MS-based proteomics identified peptides mapping to bacteria (38.3%), eukaryotes (43.5%) (including host), archaea (< 0.1%), and viruses as well as a significant proportion of unassigned peptide intensities (18.2%) (**Fig. S1**). In contrast, a majority of 16S reads (> 99%) were attributable to bacteria with a much smaller proportion attributable to archaea (< 0.1%) and < 0.1% remaining unassigned (**Fig. S1**).

Large discrepancies in taxonomic resolution begin to emerge at the phylum level. By LC-MS/MS proteomics, we identified 46 phyla, including Ascomycota, Basidiomycota, Spirochaetes, Chordata, and Streptophya in addition to the 8 identified by 16S rDNA sequencing alone (Euryarchaeota, Actinobacteria, Bacteroidetes, Cyanobacteria, Firmicutes, Fusobacteria, Proteobacteria and Verrucomicrobia) (**Fig. 1**). Firmicutes account for only 22.7% of the sample composition according to the LC-MS/MS; however, this phylum dominates the composition of patient microbiota according to 16S amplicon sequencing and accounts for 86.2% of all reads. Interestingly, the abundance of Bacteroidetes was relatively minimal by 16S rDNA sequencing (except for samples H5, UC9, UC15, UC23) yielding a Bacteroidetes:Firmicutes ratio of approximately 1:100 (**Fig. 1**). In contrast, Bacteroidetes account for an average of 4.2% of the microbiota peptide content by LC-MS/MS-based proteomics yielding a Bacteroidetes:Firmicutes ratio of approximately 1:5. The majority of identified peptides identified via metaproteomics predominantly originate from the Chordata phylum and are presumably host-derived. Additionally, a significant number of peptides are derived from the Streptophyta and are attributable to a variety of dietary plants, including *Solanum tubersum* (potato), *Seasum indicum* (sesame), *Theobroma cacao* (chocolate), *Zea mays* (corn), and *Oryza sativa* (rice) among many others (**SI_A**). Where comparable, we posit that differences in DNA extraction efficiencies (*e.g.,* Gram^+^ vs Gram^−^), differences in metabolic/secretory activities, and shared tryptic peptides between microbes likely contribute to the discrepancy in taxonomic compositions between 16S sequencing and proteomics methods.

**Figure 1.**
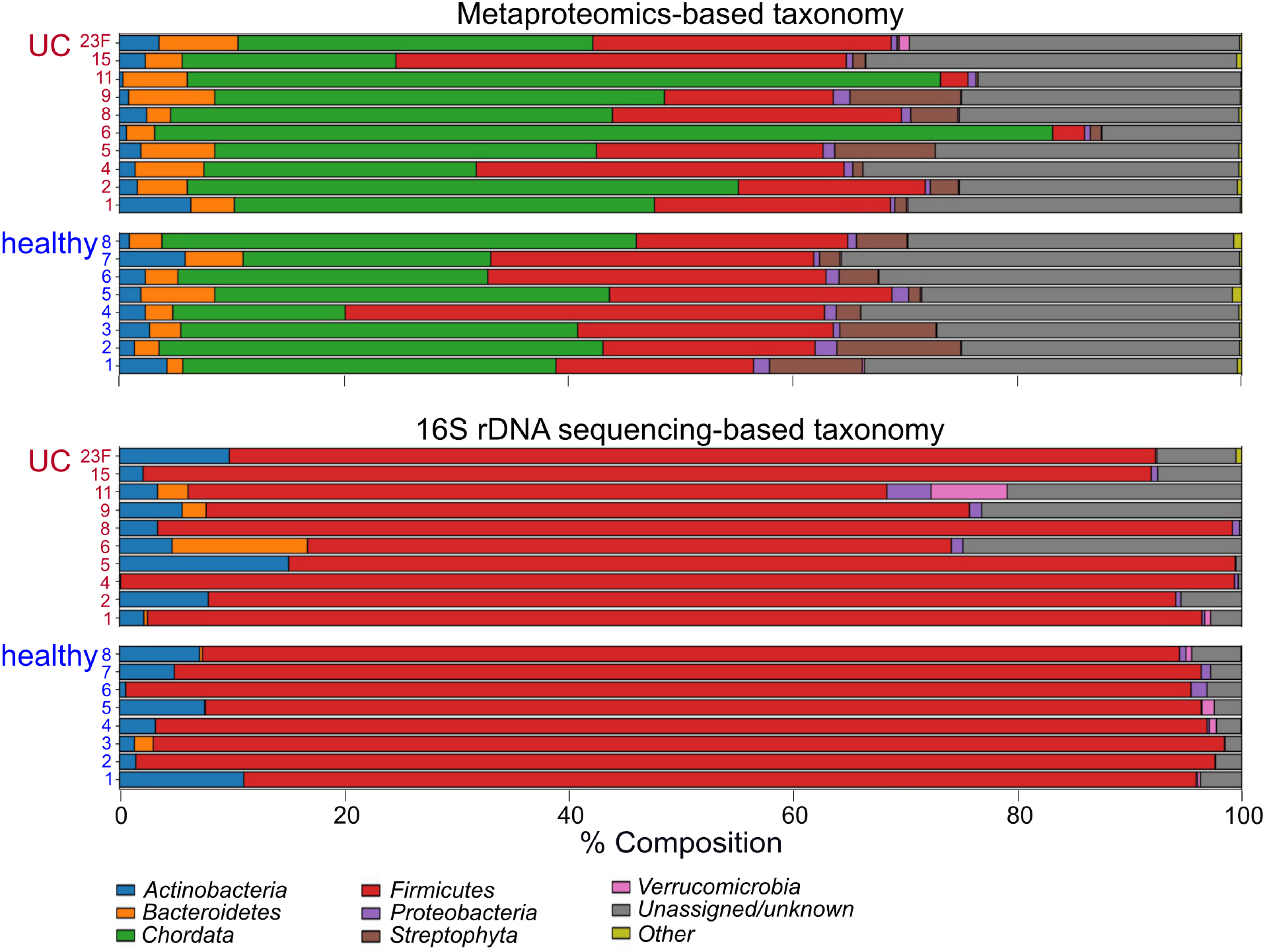
Relative abundance plots for all 18 patient samples comparing microbiome phylum-level taxonomy by 16S amplicon sequencing (lower 18 bars) against LC-MS/MS bottom-up proteomics (upper 18 bars).

The trend of increased taxonomic diversity across microbiota samples observed by LC-MS/MS proteomics over 16S amplicon sequencing is conserved at each classification tier (**Fig. S1-S7**). At the species level, we identified a total of 848 species/strains by LC-MS/MS and only 38 by amplicon sequencing. Despite the predicted increase in diversity by LC-MS/MS, an average of 85.9% of the identifiable peptides are not mappable to a particular species. This observation is due to the redundancy and conservation of microbial proteins across distinct and divergent species. Thus, taxonomic predictions based on the peptide composition in samples becomes increasingly difficult with more granular classification levels.

### 176 protein groups are significantly altered between the healthy and UC cohorts

Differential expression analysis yielded 176 protein groups (from host, microbes, and diet) significantly altered between healthy and UC patients (*p* ≤ 0.005 and *q* < 0.1), with 65 groups enriched in healthy volunteers and 111 protein groups enriched in the UC volunteers (**Fig. 2A** **and SI_B**). Principal components analysis (PCA) of the dataset revealed a modest but distinguishable separation between the proteomic composition of healthy and UC fecal samples (**Fig. 2B**) which is in agreement with the Euclidean distance matrix generated for the same dataset (**Fig. 2C**). 29 of the 111 protein groups significantly enriched in UC fecal samples were host-derived; however, no host protein groups were enriched (*q*-value < 0.1) in the healthy individuals. STRING analysis of significantly enriched UC host proteins yielded 26 edges among 25 nodes and a highly significant protein-protein interaction (PPI) enrichment *p-*value of 1.84×10^−12^ at medium confidence (0.400) suggesting a strong association between tested proteins (**Fig. 2D**, **Table SI_C**) (49). At highest confidence (0.900), 13 edges between 25 nodes were found reinforcing a significant PPI enrichment *p*-value of 2.0×10^−11^. At medium confidence, the most significant reactome pathways (fdr < 0.001) were neutrophil degranulation, innate immune system, antimicrobial peptides, immune system, and metal sequestration by antimicrobial proteins. GO biological process terms (fdr < 3.3×10^−8^; regulated exocytosis, neutrophil degranulation, secretion by cell, transport, leukocyte mediated immunity, and antimicrobial humoral response) and GO cellular component terms (fdr < 2×10^−7^; cytoplasmic vesicle lumen, secretory granule, secretory granule lumen, vesicle, cytoplasmic vesicle part, and cytoplasmic vesicle) associated with host proteins all support the assertion that host immune-related secretory events are prevalent in the gastrointestinal tracts of UC patients. Interestingly, because no host proteins were significantly enriched across healthy fecal samples, we posit that host-centric biological pathways associated with colitis occur in addition to rather than in lieu of processes associated with homeostasis.

**Figure 2.**
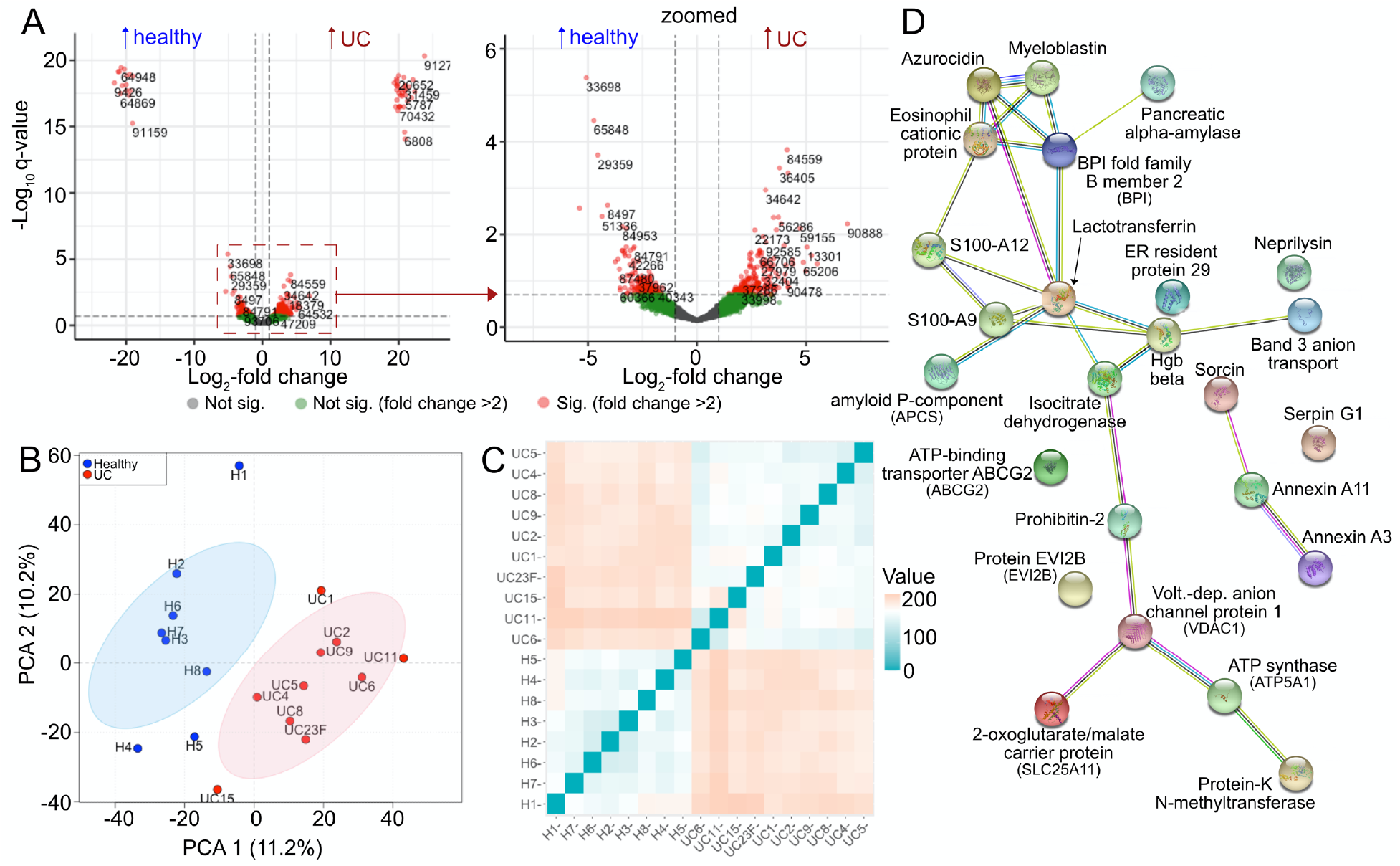
(A) Volcano plot depicting differentially detected protein groups from patient fecal samples; whole plot (left), zoomed-in plot (right), red=significant (*q* < 0.1, foldchange > 2), green=not significant (*q* > 0.1, foldchange >2), grey = N.S. (*q* > 0.1, foldchange ≤ 2). (B) Principal component analysis of all 18 patient samples across all differentially tested contrasts; red = healthy individuals, blue = UC patients. (C) Euclidean distance plot of all 18 patient samples across all differentially tested contrasts, blue = more similarity, orange/red = less similarity. (D) STRING protein network analysis of host protein groups differentially enriched in UC patients’ fecal samples at medium confidence (0.400); edges: known interactions (aqua: from curated database, magenta: experimentally determined), predicted interactions (green: gene neighborhood, red: gene fusions, blue: gene co-occurrence), others (chartreuse: textmining, black: co-expression, lavender: protein homology).

More than half of the non-host protein groups have limited to no annotations despite being significantly altered. For example, 37 of 65 and 48 of 82 non-host protein groups enriched in the healthy or UC cohorts, respectively (*p* ≤ 0.005 and *q* < 0.1), were poorly annotated in the ComPIL database (*i.e.,* no annotation, annotated as hypothetical proteins or as domain of unknown function-containing proteins). While additional BLAST homology searches and InterProScan analyses were performed on each poorly annotated protein group, we were unable to make signficant additional annotations for 11 of 37 protein groups enriched in the healthy cohort and 15 of 48 protein groups enriched in the ulcerative colitis cohort (**Table SI_C**) (50–52). Such protein groups represent interesting targets for structural and biochemical validation as further inquiry could elucidate their possible roles, in the propagation of inflammatory or anti-inflammatory processes.

A majority of significantly enriched, non-host protein groups were annotated among both healthy and UC cohorts with predicted and/or biochemically verified functions ranging from metabolism (glyceraldehyde-3-phosphate dehydrogenase and translation elongation factor Tu) to defense (type II secretion system protein). Notable non-host entries that were increased in healthy volunteers include an acid-soluble spore protein (WP_071120403.1), methylene tetrahydrofolate reductase (SRS064276.159392-T1-C), and fruit bromelain (BROM1_ANACO). The enriched small spore protein is the only entry in its protein group and originates from the recently described bacterium *Romboutsia timonensis,* whose depletion has been associated with colorectal cancer incidence (53, 54). The methylene tetrahydrofolate reductase protein group enriched in the healthy cohort contains 29 members possessing similarity scores in the range of 98.6-100%. By BLAST analysis, these reductases likely originate from the *Lachnospiraceae* family of bacteria, and their examination could provide a glimpse into the microbial B-vitamin economy that importantly underpins host homeostasis, as humans are unable to *de novo* synthesize many essential B vitamins (55–57). Interestingly, fruit bromelain detected in 4 of 8 healthy patient fecal extracts is a pineapple-derived cysteine protease we did not expect to encounter (58, 59). This protease is commonly sold as an over-the-counter supplement or as a component of meat tenderizers, and its detection may be an artifact introduced through patients’ diets.

With respect to protein groups enriched in UC patients, notable annotated non-host entries include hyaluronon glucosaminidase (CLONEX_02131), a transglycosylase SLT domain-containing protein (HMPREF0462_0704), and a metallohydrolase (WP_081140786.1). The enriched hyaluronan glucosaminidase group contains 4 members with almost identical sequences (99.95-100% identity). These proteins likely originate from the *Lachnospiraceae* family members *Tyzzerlla* or *Coprococcus*. Hyaluronan is a high-molecular weight carbohydrate component of the human extracellular matrix that can serve as an inflammatory/injury signal for host immune receptors when degraded by host hyalurondases (60). Hyaluronon glucosaminidase activity may exacerbate host inflammatory processes and contribute to UC-related inflammation, as well as afford microbes the ability to infiltrate host barriers. The transglycosylase SLT (lytic) domain-containing protein group includes 70 members with sequence identities of 95.71-100% compared to the *Helicobacter pylori* (*H. pylori*) enzyme. These enzymes catalyze the non-hydrolytic intramolecular cyclization of *N*-acetylmuramyl residues on bacterial cell walls propagating the cell wall remodeling process (61, 62). For *H. pylori*, previous studies demonstrate the importance of transglycosylase-generated cell wall muropeptide fragments to inducing host inflammation which can in turn promote gut colonization (63, 64). Finally, the enriched metallohydrolase originates from *Pantoea latae* and is annotated as a non-peptide amide C–N bond hydrolase that may inactivate amide-containing molecules such as lactams, which are contained in an important class of antibiotics (65, 66). A complete list of protein groups found differentially expressed in this patient cohort can be found in the supplement (**SI_C**).

### Serine-type endopeptidase activity is significantly enriched in UC samples

For GO term relative abundance analysis, we used averaged peptide intensities for peptides within the same protein group to account for comparisons between proteins of different lengths. Of the 4,622 protein groups selected for relative abundance plotting we identified 575, 394, and 85 terms for the molecular function, biological process, and cellular component GO namespaces, respectively. In general, GO relative abundance breakdowns between all samples for each namespace appear similar by unweighted assembly (**Fig. 3B**, **S8-S10**). However, when weighted by corresponding ion intensities, GO term relative abundances between samples differ dramatically (**Fig. 3A**, **S8-S10**). The ‘None’ and ‘Other’ categories occupy the largest areas both by unweighted and weighted assembly for all three GO namespaces. With respect to molecular function, global relative abundances (when averaged over all samples) for the terms glutamate-cysteine ligase activity (GO: 0004357), aminopeptidase activity (GO: 0004177), and serine-type endopeptidase activity (GO: 0004252) expand 31-, 18-, and 14-fold respectively going from an unweighted to weighted assembly. Conversely, global relative abundance for the terms enoyl-[acyl-carrier-protein] reductase (NADH) activity (GO: 0004318) and mismatched DNA binding (GO: 0030983) contract >50-fold going from unweighted to weighted assembly. Similar relative abundance expansions and contractions going from unweighted to weighted assemblies were observed for the biological process and cellular component namespaces (**Fig. S9, S10**).

**Figure 3.**
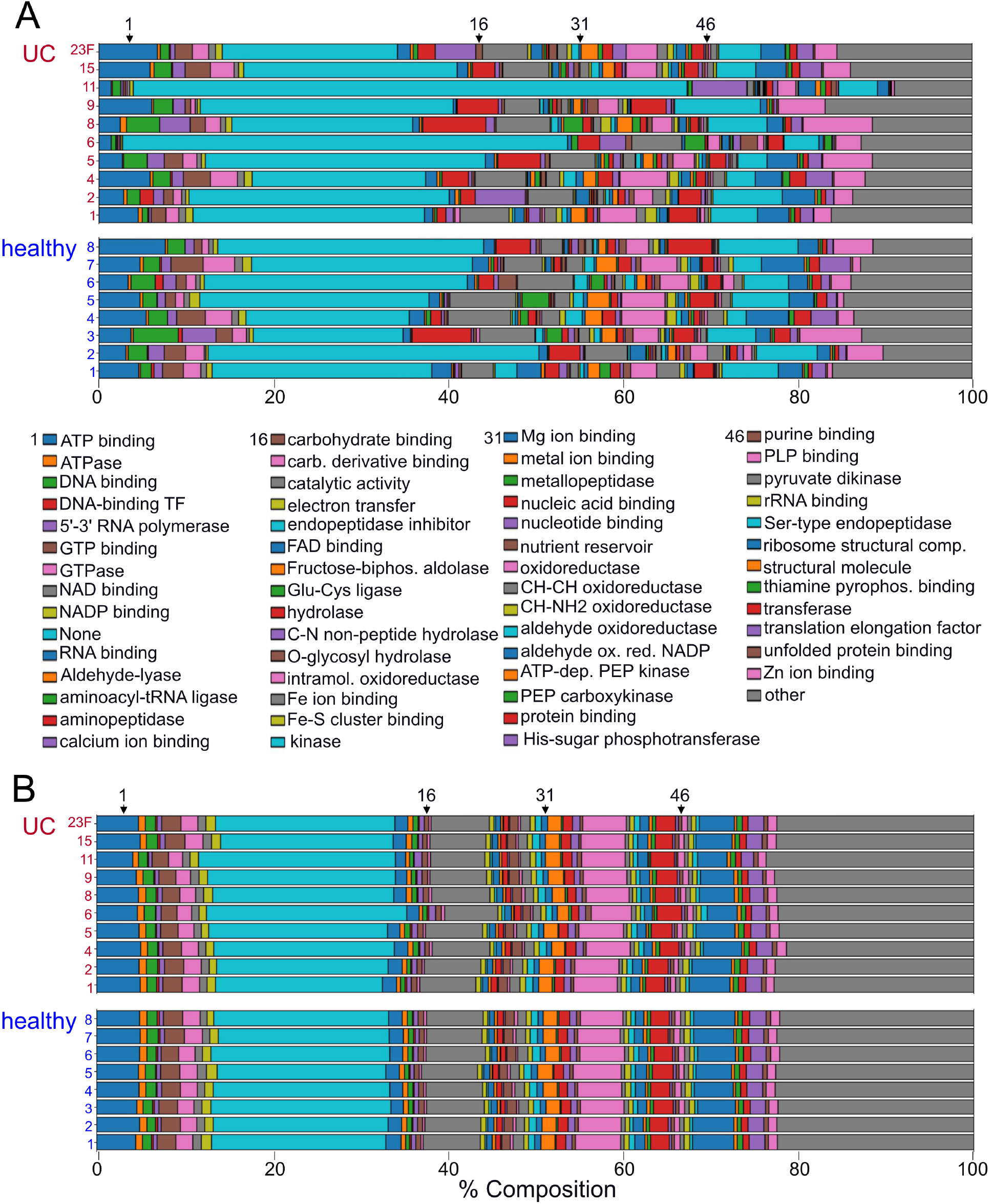
Relative abundance plots for all 18 patient samples comparing microbiota GO molecular function abundance by LC-MS/MS using either (A) weighted measures (upper 18 bars) or (B) unweighted measures (lower 18 bars).

The large 31-fold unweighted-to-weighted relative abundance expansion of glutamate-cysteine ligase activity could be attributed to one protein group with one member (WP_027345637.1). This enzyme originates from *Hamadaea tsunoensis* and catalyzes a key step in the synthesis of glutathione, a key antioxidant for the microbiota (67). The 18-fold unweighted-to-weighted relative abundance expansion observed for the aminopeptidase activity term originated from 19 protein groups with one very dominant protein group (WP_027209280.1) representing 96.2% of the GO term namespace’s relative abundance. This predicted M18 family protease originates from *Butyrivibrio hungatei* but currently has no structural or biochemical annotation. Finally, the 14-fold expansion observed for the serine-type endopeptidase activity GO term is attributable to 38 protein groups with the majority share originating from host (85.7%) and minor shares originating from pig (13.5%) and microbes (0.8%). The serine-type endopeptidase from pig is an artifact, as it originates from the sequencing grade porcine trypsin used to generate peptides for LC-MS/MS analysis. Interestingly, human chymotrypsin-like elastase 3A (CEL3A_Human) and chymotrypsin-C (CTRC_human) are more abundant than porcine trypsin, resulting in 35.8% and 14.9% of the GO term share compared to porcine trypsin. Other prominent protein groups (less abundant than porcine trypsin), include chymotrypsin-like elastase 3B (CEL3B_human), cathepsin G (CATG_human), and trypsin-1 (TRY1_human) and comprise 11.3%, 5.1%, and 3.0% of the serine-type endopeptidase activity GO term, respectively.

We performed GO enrichment analysis in the R GOstats package with GO terms corresponding to differentially expressed protein groups serving as the enriched GO term set and GO terms mapping to all non-differentially expressed protein groups as the “universe” set (68). GO terms overrepresented in either healthy or UC cohorts for each GO namespace (*p* < 0.01) are listed in the supplement (**Fig. S11-S18)**.

For the healthy patient cohort, 9 molecular function GO terms were enriched including hydrolase activity (hydrolyzing O-glycosyl compounds) (GO:0016787), cysteine-type peptidase activity (GO:0008234), rRNA binding (GO:0019843), and oxidoreductase activity (acting on iron-sulfur proteins as donors) (GO:0016730) among the most significantly enriched (*p* < 0.001, odds ratio > 10). The biological process namespace contained 9 GO terms enriched in healthy patients with polysaccharide catabolic process (GO:0000272), homeostatic process (GO:0042592), sulfur amino acid metabolic process (GO:0000096), nitrogen fixation (GO:0009399), and asexual sporulation (GO:0030436) among the most significantly enriched terms (*p* < 0.001, odds ratio > 20). Only 3 GO cellular component terms were found to be enriched in healthy patients including oxidoreductase complex (GO:1990204), vanadium-iron nitrogenase complex (GO:0016613), and endospore-forming forespore (GO:0042601) (*p* < 0.001, odds ratio > 100). Together the 3 GO namespaces strongly associate carbohydrate processing and microbial sporulation activities with healthy patients.

For the ulcerative colitis cohort, 21 terms were enriched in the molecular function GO namespace including calcium ion binding (GO:0005509), peptidase activity (acting on L-amino acid peptides) (GO:0008233), lipid binding (GO:0008289), catalytic activity (acting on a protein) (GO:0140096), serine-type endopeptidase activity (GO:0004252), serine hydrolase activity (GO:0017171), and calcium-dependent phospholipid binding (GO:0005544) (*p* < 0.001, odds ratio > 10) (**SI_D**). The biological process namespace contains 18 entries including proteolysis (GO:0006508), aromatic amino acid family metabolic processes (GO:0009072), propionate metabolic process (methylcitrate cycle) (GO:0019679), acetate metabolic process (GO:0006083), short-chain fatty acid metabolic process (GO:0046459), antibacterial humoral response (GO:0019731), antifungal humoral response (GO:0019732), regulation of cytokine production (GO:0001817), and response to fungus (GO:0009620) among the most enriched terms (*p* < 0.003, odds ratio > 10). The enriched cellular component term list was much shorter with only 5 entries, including extracellular space (GO:0005615), organelle lumen (GO:0043233), endoplasmic reticulum lumen (GO:0005788), intracellular membrane-bound organelle (GO:0043231), and extracellular region (GO:0005576) (*p* < 0.008, odds ratio > 10). Together, these enriched GO terms associate the extracellular space/secretome and antimicrobial host responses with UC patients in our study. Proteolysis and serine-type endopeptidase activity are highly enriched terms that likely occur in concert with antimicrobial host responses as many of the prominent constituents of the proteolysis and serine-type endopeptidase activity GO terms are proteins of host origin.

### ABPP confirms presence of active serine-type endopeptidases and identifies previously undetected serine-type endopeptidases

We treated 3 patient fecal samples with a biotinylated flurophosphonate-based (FP-based) serine-reactive probe to label and thus establish whether serine hydrolases were active in patient fecal samples (**Fig. 4A**) (69). Biotinylated fluorophosphonate probe (FP probe)-labeled proteins were enriched with streptavidin-agarose beads, visualized by Western blot (**Fig. 4B**), and identified by LC-MS/MS analysis using a previously described 2-step large-to-focused database search strategy (70). Analysis of the LC-MS/MS data revealed several hundred non-contaminant protein sequences. These sequences were clustered into 104 distinct protein groups (95% similarity cutoff using CD-HIT) and further reduced to 63 protein groups with highly homologous host proteins condensed together. Of note, 27 and 35 protein groups derived from host (**Fig. 4C**) and non-host (**Fig. 4D**) were identified, respectively.

**Figure 4.**
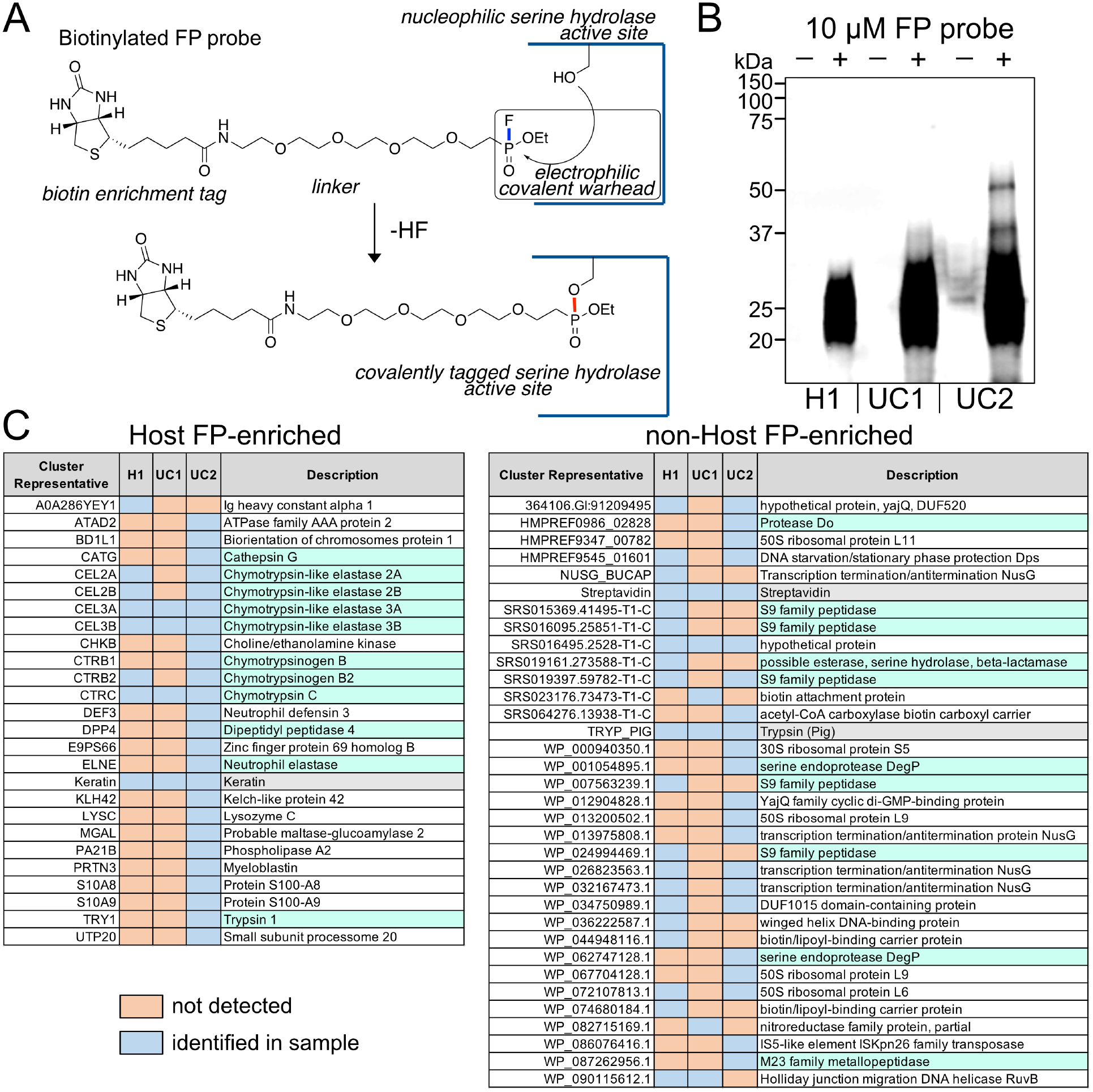
(A) FP probe structure and reaction schematic for covalent attachment to nucleophilic active-site serine in hydrolases. (B) Western blot of 3 patient fecal lysates treated and enriched with FP probe followed by streptavidin bead enrichment visualized with fluorophore-conjugated streptavidin. Host (C) and non-host (D) proteins from patient fecal samples enriched by FP probe and detected by LC-MS/MS (blue = protein detected in corresponding patient sample, orange = protein not detected in corresponding patient sample).

The majority of host-derived protein groups labeled and enriched with the FP probe were also identified within the unenriched LC-MS/MS datasets (**Fig. SI_C**). 14 of 27 FP probe-enriched host proteins are known serine hydrolases including the chymotrypsin-like elastase family (2A, 2B, 3A, 3B), cathepsin G, dipeptidyl peptidase 4, neutrophil elastase, myeloblastin, trypsin 1, and phospholipase A2 (**Fig. 4C**). Enrichment of these particular hydrolases over all other host proteins provides confidence that the FP probe is selective for nucleophilic serines in the tremendously complex fecal protein matrix. Aside from demonstrating that the hydrolases are active, these results also suggest that this fraction of proteases remain uninhibited by anti-proteolytic proteins often found in the gut (71–73).

Most non-host proteins are microbial in origin with the exception of streptavidin and porcine trypsin introduced during sample preparation (**Fig. 4D**). The most promising FP probe-susceptible proteins, include protease Do entries (DegP), S9 family peptidases, and a beta-lactamase, as determined by sequence analysis. Interestingly, of the 10 identified non-host serine hydrolase protein groups, only 1 (SRS019397.59782-T1-C) was detected without FP-probe demonstrating the utility of chemical-based enrichment strategies for the identification of novel proteins in a complex environment. Of the 167,554 MS2 spectra collected for all FP-enriched LC-MS/MS data sets, only 5,352 (3.2%) were confidently assigned by ComPIL database searches. There is a strong likelihood that other serine hydrolase-derived peptides are present in our microbiota samples but they remain unidentified due to limitations imposed by the incompleteness problem associated with metaproteomics database searching (22). Unfortunately, the techniques for database-independent, high-confidence identification of these peptides and their parent protein sequences are currently not well established.

### *De novo* peptide sequencing enables glimpses into the dark peptidome

Of the 2,829,920 MS2 fragmentation spectra we collected overall, only 503,244 (17.8%) were matched to a corresponding peptide with a 1% peptide false positive rate using a target-decoy search strategy paired with the ComPIL database. The modest number of peptide spectrum matches we observed is likely attributable to (1) a loss in filtering sensitivity that often accompanies database expansion and (2) a never-complete database that is perennially associated with metaproteomics. We posit that a non-trivial portion of unmatched MS2 spectra map to either known or unknown peptide sequences, and we aim to estimate the size of unmatched MS2 spectrum space or “the dark peptidome,” using a complimentary *de novo* peptide sequencing approach (74).

We subjected MS2 spectra from all patient fecal sample LC-MS/MS data sets to *de novo* peptide sequencing using the Novor algorithm (75). Novor attempts to deduce peptide sequence from MS2 fragmentation spectra, generating a *de novo* peptide spectrum match (PSM) and an accompanying confidence score (Novor score; higher scores indicated better predicted matches). Where available, we paired *de novo* PSMs with their corresponding database PSMs (ComPIL2-assigned) and calculated an additional Novor-ComPIL2 similarity score (*de novo*-database similarity score) based on the Needleman-Wunsch comparison algorithm (raw scores were scaled to 100, where 100 represents a perfect match) (76). We used these values to construct histograms (**Fig. 5A**, **S19, S20**) and joint plots (**Fig. 5B**, **S21, S22**) depicting possible unidentified peptide space in patient fecal microbiota samples.

**Figure 5.**
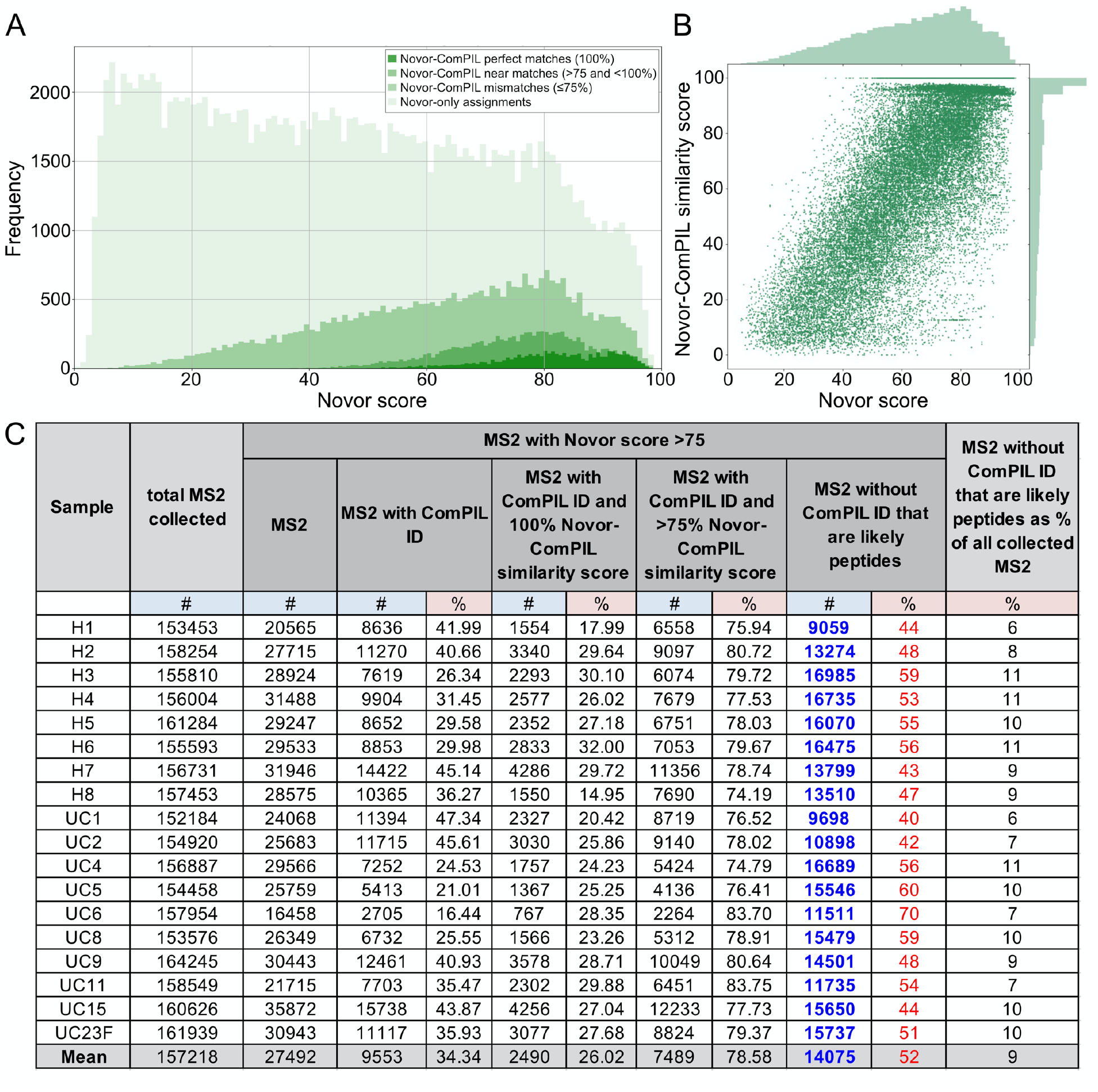
(A) Histogram of all assigned H2 patient sample MS2 spectra grouped by Novor score (0-100); darkest green area represents MS2 correctly assigned by Novor determined by comparison to ComPIL (database) result; lightest green area represents MS2 without ComPIL peptide assignment. (B) Joint plot of H2 patient sample MS2 spectra depicting correlation between Novor score (0-100) and Novor-ComPIL similarity score (0-100). (C) Number of MS2 spectra from all patient samples with Novor scores >75 that likely represent peptides but do not have ComPIL peptide matches (“dark peptidome”).

ComPIL database-assigned PSMs were non-uniformly distributed along the Novor score axis with a larger proportion of database PSMs grouped near the high-confidence *de novo* sequencing Novor scores (70-100) (**Fig. 5A**). While perfect *de novo*-database agreements were rare, a large proportion of database PSMs possessed strong similarity to *de novo* PSMs, a relationship best depicted by a Novor score versus Novor-ComPIL2 similarity score joint plot (**Fig. 5B**). Thus, it is reasonable to expect that above a conservative cut-off value (Novor score = 75), significant numbers of MS2 spectra without database assignments correspond to peptides that either are not contained in the ComPIL2 database or were rejected due to our high search-filter stringency. Based on this assessment, it is estimated that an average of 14,075 MS2 spectra with a Novor score of 75 or greater remain unidentified per sample (**Fig. 5C**). This corresponds to approximately 9% of all MS2 spectra per patient sample, which could increase global identifications by approximately 50%. BLAST analysis of several *de novo* PSMs which do not have corresponding database PSMs returned reasonable, high-similarity score matches to microbial peptides from the NCBI non-redundant database supporting this assertion. Below a Novor score of 75, there are likely many unidentified peptide-MS2; however, these peptides are also likely intermixed with many non-peptide-MS2.

## Discussion

16S amplicon sequencing has been a workhorse technique for microbiota studies over the last decade due in large part to the simplicity of material extraction and the broad availability of resources needed to generate meaningful data. In contrast, the use of LC-MS/MS-based metaproteomics profiling has been more sparse for the opposite reasons. While functional proteomic interrogation of the microbiota by LC-MS/MS was our main goal, we considered it important to contrast our proteomics-based taxonomy findings with those generated by 16S sequencing, a technique more familiar to the microbiome research community. Gut microbiota taxonomy through the lens of 16S amplicon sequencing versus LC-MS/MS-based proteomics is expectedly different, but in some unexpected ways (30). At the phylum level, we expected to see a similar configuration by both techniques with the exception that proteomics would include a small concession for host proteins. Unexpectedly, we observed both host and diet-derived (potato, rice, corn, *etc*.) proteins in great relative abundance to microbial proteins. Our analyzed samples were pre-enriched for microbes by filtration, differential centrifugation, and several washing steps, yet nearly half of the detected proteome was mapped back to host or dietary plant proteins. Another unexpected finding was a discrepancy between the relative abundance ratios of Bacteroidetes and Firmicutes. This discrepancy could be artifactual and originate from differences in DNA extraction efficiencies between microbes, but it could also be the result of biologically relevant phenomena. For example, overrepresentation of Firmicutes by 16S sequencing could stem from an abundance of Firmicutes cells that are metabolically inactive (spores) relative to Bacteroidetes cells (77). This would be in agreement with our finding that the asexual sporulation GO term is enriched in healthy patient samples. Finally, we found that at more granular taxonomic strata (**Fig. S1-S7**), the number of identifiable organisms was unexpectedly greater for proteomics than for 16S sequencing. By proteomics, we observed hundreds of distinct organisms at the species level versus several dozen by 16S. Expectedly, the proportion of uniquely mappable peptides progressively decreased at more granular taxonomic levels such that at the species level, about 75% of all peptide intensity could not be mapped to a particular species. Because peptides are proxies for both taxonomy and function, this observation hints at a functional redundancy among microbes in the gut, which could be better examined by differential expression and GO term analysis of proteomics data.

One of the leading motivations for performing differential expression analyses on microbiota samples is to identify specific biomarkers or disease-associated microbiota proteins for further examination. Toward this goal, we identified 176 protein groups significantly enriched (*q* < 0.1) in either healthy or UC volunteers, with a major share originating from microbes. Interestingly, no host proteins were identified as significantly enriched in the fecal extracts of healthy volunteers while several were found enriched in the fecal extracts of UC patients. Among the host proteins enriched in UC patients, we identified the calibrating entry, protein S100-A9 (*p* < 0.004, *q* < 0.07), a component of fecal calprotection and known IBD biomarker (78, 79). According to STRING and reactome analyses, many host proteins enriched in UC patients are also inflammation-aligned lending more credibility to the prospect that the enriched proteins we have identified are truly UC-associated. For a comprehensive list of enriched protein groups, see **SI_C**. While most enriched protein groups had some annotation, a significant portion had little to none. This finding presents an exciting opportunity for the structural and biochemical study of novel sequences. Given the enormous number of domains of unknown function (DUF) and unknown function-type proteins catalogued from microbiome metagenomic sequencing efforts, we are faced with a prioritization problem wherein the most disease-relevant sequences are obscured by less impactful ones (80). LC-MS/MS-based proteomics appears in this context to be an important tool for identifying sequences that are both expressed and biologically relevant, which will help focus our future studies. Lastly, it is important to point out that poorly annotated proteins (*i.e.,* proteins without GO assignments) factor weakly or not at all into broader functional analyses like GO enrichment simply due to the nature of enrichment testing (*i.e.,* hypergeometric). Therefore, novel sequences without known biological- or disease-relevance are important to eventually characterize.

Within the detected microbiota proteome, known functional diversity is high with several hundred molecular function GO terms represented. A flat depiction of molecular function wherein a 1-sequence-1-count paradigm is applied reveals a consistent relative abundance configuration between all samples. Interestingly, when an intensity-based paradigm is applied, such that relative abundance is weighted by peptide intensity (a proxy for protein abundance), a very different picture emerges. The relative abundance of many molecular function GO terms shift, sometimes dramatically. One of the most conspicuous terms to us was “serine-type endopeptidase activity” that expanded an average of 14-fold among all patient samples going from count-based to intensity-based representation. Additionally, we found this same term enriched in UC patient fecal samples by hypergeometric testing, warranting a closer inspection of the protein groups that contribute to this term. We found that the major contributors were host-derived serine proteases (85.7%) such as chymotrypsin-C and the chymotrypsin-like elastase family with minor contributions from porcine trypsin (13.5%) (an artifact of sample preparation), and microbial serine proteases (0.8%). The high relative abundance of serine proteases in fecal samples is not surprising given that they are important components of host digestive enzyme cocktails secreted into the gut lumen. We were however, surprised to find both host and microbial serpins, which are known active-site directed suicide inhibitors for serine/cysteine proteases, in fecal samples. This observation suggests that there might be important regulatory host-microbe cross-talk with respect to proteolytic activity that occurs in the gut. By comparing the abundance of proteases or serpins, it would still be difficult to determine which and what fraction of serine proteases remained active upon fecal sample collection. To identify active serine proteases, we treated fecal samples with an active-site directed serine-hydrolase selective chemical probe (FP probe) for labeling, enrichment, and target identification via LC-MS/MS (ABPP) (25, 26, 69). We examined 3 patient fecal samples (1 healthy, 2 UC) and qualitatively found human chymotrypsin-like elastases 3A and 3B and chymotrypsin-C enriched and, therefore active in all samples. For one UC sample (UC2), we identified additional FP probe-enriched host proteases including cathepsin G, chymotrypsin-like elastase 2A and 2B, dipeptidyl peptidase 4, neutrophil elastase, and trypsin 1, lending support to the idea that aberrantly increased protease activity is associated with IBD (72, 81). In addition to host proteases, we identified several microbial proteases from all 3 patient samples upon FP probe enrichment. Surprisingly, we found that 9 of 10 microbial proteases were not detected by LC-MS/MS at all without FP probe enrichment. This finding suggests that there are likely many microbial proteases expressed in the gut microbiota and that they are likely below the limit of detection by most current sampling and LC-MS/MS profiling strategies. We speculate that this sentiment also holds true for other low-abundance, high-impact protein functionalities, underscoring the importance of pre-enrichment strategies for future proteomics studies.

In addition to sampling limits, many microbiota peptides that are sampled by LC-MS/MS are liable to go undetected due in large part to database-completeness limitations. We attempted to estimate the number of peptide-likely fragmentation spectra per LC-MS/MS experiment using a *de novo* sequencing tool (Novor) in order to define a rough boundary around the amount of unassigned peptide space captured by the mass spectrometer but unidentified by our database workflow. Random homology searching of high-confidence peptide-like fragmentation spectra revealed high numbers of exact and near-matches to peptide sequences in the NCBI non-redundant database. Though *de novo* peptide sequencing coupled to homology searching can help capture database-elusive peptides in more defined contexts (74), we were reluctant to rely more heavily on this strategy without a stringent methodology for distinguishing between sequencing errors and truly homologous peptide sequences, especially in a context as taxonomically diverse as the gut microbiota. An additional layer of difficulty rests in determining how to treat *de novo*-only peptides that are constituents of completely unknown parent protein sequences. Additional deep shotgun genome sequencing is an obvious path forward, but for protein-coding genes that elude this approach, perhaps genomics-agnostic, middle- and top- down proteomics sequencing could be applied (82, 83). We anticipate that the expanded use of ABPP techniques in the microbiota will enrich for many protein sequences not contained in large databases like ComPIL, and robust high-throughput methods for identifying these whole novel protein sequences will be needed.

## Conclusions

We deployed high-resolution LC-MS/MS and a comprehensive database search to identify host and novel microbial proteins enriched in the fecal extracts of healthy volunteers and ulcerative colitis patients. This process led us to identify 176 discrete protein groups and several protein functions associated with ulcerative colitis, with the function “serine-type endopeptidase activity” featuring prominently. We also identified host and microbial serine protease inhibitors in concert with serine proteases. We used an activity-based chemical tagging strategy (ABPP) to enrich for serine hydrolases/proteases and showed that these enzymes are still active in the gut despite the presence of active-site directed protease inhibitors. This strategy also revealed the presence of previously undetected serine proteases demonstrating the utility of activity-based tagging for the amplification of low-abundance proteins. Finally, we paired our database proteomics strategy with *de novo* peptide sequencing to estimate the size of high-confidence peptide space in our samples that remains unidentified despite the use of a large comprehensive database.

## Materials and Methods

### Patient sample collection

Human subject research was approved by the Institutional Review Board (IRB) at Scripps Research and Scripps Health; protocol #: IRB-14-6352. Volunteers self-collected their own stool samples using administered standardized in-home sample collection kits and were instructed to immediately freeze specimens at −20°C. Samples were stored in provided consumer-grade −20 °C mini-freezers immediately after collection, transported by courier services on dry ice, and stored in laboratory-grade freezers at −20 °C until microbial extraction. Collected stool samples were highly heterogeneous in color, texture, and viscosity both before and after microbial extraction.

### Microbe extraction

Stool samples were thawed to room temperature, diluted in PBS (pH 7.4), vortexed thoroughly to yield slurries, and then centrifuged at 100 xg for 1 min. The flocculent upper layer was extracted, filtered over 70 μm nylon mesh cell strainers to remove large, recalcitrant masses, and then centrifuged at 8000 xg for 5 min to pellet. Pellets were rinsed twice with PBS, then resuspended in PBS to a density of 100 mg wet microbial pellet per 500 μL of suspension.

### DNA extraction, 16S amplicon sequencing, and data processing

Microbial DNA was extracted from thawed fecal microbe aliquots using a fecal/soil extraction kit (Zymo Research, Irvine, CA, USA). 50-100 ng of DNA per patient sample were submitted to the Scripps Research genomics core for next-generation sequencing which was performed using the MiSeq platform (Illumina Inc, San Diego, CA, USA). For taxonomy based on 16S rDNA amplicon sequencing, we targeted the bacterial 16S V4 region using a 300bp paired-end approach aiming for 100,000 reads per sample. Reads were taxonomically mapped using QIIME2 (and associated plug-ins) and classifiers created from the SILVA 132 database (84, 85). For access to raw data, see Zenodo repository (doi: 10.5281/zenodo.4265371). For detailed methods, see supplementary information.

### Protein extraction, protein digestion, proteomics data collection

Protein was extracted according to a previously described protocol (45). Extracted protein was resuspended in H_2_O and concentration was measured by BCA assay (ThermoFisher, Waltham, MA, USA). Extracted microbiome protein (100 μg) was reduced, alkylated, trypsinized, and desalted (ZipTip C18, MilliporeSigma, Burlington, MA, USA) prior to LC-MS/MS analysis. Desalted peptides (1 μg) were separated using a 4 hour C18 gradient by nano-flow liquid chromatography coupled to an Orbitrap Fusion Tribrid (ThermoFisher, San Jose, CA) operating in data dependent mode. Both MS1 and MS2 spectra were recorded in the Orbitrap at 120K and 30K resolution, respectively. For detailed methods, see supplementary information.

### Proteomics data analysis

We collected a total of 2,829,920 MS2 spectra between all 18 patient samples. These spectra were searched against the ComPIL 2.0 database (contains 4.8 billion unique tryptic peptides from >225 million forward and reverse protein sequences) (23, 24) using the ProLuCID/SEQUEST search engine (20, 46, 47). 503,244 (17.8%) MS2 spectra were mapped to 54,378 distinct peptides at a 1% peptide false discovery rate (2 peptide per protein minimum) using a target-decoy strategy (86). 576,625 protein sequences were identified and clustered into 95,000 protein groups at a 95% sequence similarity cut-off using CD-HIT (87, 88).Quantification at the MS1 level was performed using FlashLFQ with a match-between-runs strategy enabled (10 ppm precursor tolerance, 15 min window) (89). Peptide MS1 area-under-the-curve intensities were mapped to protein groups. Intensity belonging to peptides that map to >1 protein group were discarded. After removing protein groups with too many missing values, 4,622 protein groups remained for differential expression analysis which was performed using Limma as part of the DEP package in the R statistical environment (90, 91). Protein groups were annotated with GO terms using InterProScan; these annotations were used for GO enrichment analysis in the GOstats package and for GO relative abundance analysis (50, 51, 68). Peptides were mapped to their respective taxa of origin using Unipept (92–94). Peptide intensities were then mapped to taxa to construct relative abundance tables and plots. Finally, *de novo* peptide sequencing was performed in Novor (76) and database-*de novo* peptide comparisons were performed using the Scikit-bio Python library. For more detailed methods, see supplementary information. For access to LC-MS/MS data, see PRIDE repository (PXD022433). For *de novo* datasets, protein fasta files, and protein group files (CD-HIT clusters) see Zenodo repository (doi: 10.5281/zenodo.4265371).

## Supporting information

Information on location of datasets and analyses

## Acknowledgments

We thank G. Tsaprailis and C. Sharager-Tapia for assistance with mass spectrometry data collection. We thank S. K. R. Park, T. Jung, and J. R. Yates for assistance with mass spectrometry data analysis. We thank H. Rosen, R. L. Wiseman, and L. L. Lairson for access to instrumentation. This work was supported by NIH R21AI139744 (to D.W.W.) and Boehringer Ingelheim (to D.W.W., A.I.S., and W.J.C.)

