## Supplemental Information for "Quantitative metaproteomics of patient fecal microbiota identifies host and microbial proteins associated with ulcerative colitis"

Table of Contents…… 1
Supplementary Figures…… 2-23
Figure S1. Kingdom taxonomy bar plot…… 2
Figure S2. Phylum taxonomy bar plot…… 3
Figure S3. Class taxonomy bar plot…… 4
Figure S4. Order taxonomy bar plot…… 5
Figure S5. Family taxonomy bar plot…… 6
Figure S6. Genus taxonomy bar plot…… 7
Figure S7. Species taxonomy bar plot…… 8
Figure S8. Molecular function GO relative abundance bar plot…… 9
Figure S9. Biological process GO relative abundance bar plot…… 10
Figure S10. Cellular component GO relative abundance bar plot…… 11
Figure S11. Ulcerative colitis molecular function GO enrichment bubble plot…… 12
Figure S12. Ulcerative colitis biological process GO enrichment bubble plot…… 13
Figure S13. Ulcerative colitis cellular component GO enrichment bubble plot…… 14
Figure S14. Healthy molecular function GO enrichment bubble plot …… 15
Figure S15. Healthy biological process GO enrichment bubble plot…… 16
Figure S16. Healthy cellular component GO enrichment bubble plot…… 17
Figure S17. Ulcerative colitis combined enriched GO terms table…… 18
Figure S18. Healthy combined enriched healthy GO terms table…… 19
Figure S19. Novor score histograms for healthy patient fecal microbiomes…… 20
Figure S20. Novor score histograms for UC patient fecal microbiomes…… 21
Figure S21. Novor score/Novor-ComPIL2 similarity score joint plots for healthy patient fecal microbiomes…. 22
Figure S22. Novor score/Novor-ComPIL2 similarity score joint plots for UC patient fecal microbiomes…… 23
Methods…… 24-34
General information, reagent sourcing, and instrumentation information…… 24
Chemical synthesis…… 25-28
Synthetic scheme for FP-Probe…… 25
Compound preparation procedures and characterization…… 26-28
Sample preparation and data collection…… 29-30
Microbiome extraction…… 29
Microbial DNA extraction…… 29
16S library preparation and sequencing…… 29
Microbial protein extraction and proteomics sample preparation…… 29
FP probe labeling, enrichment, and proteomics sample preparation…… 29-30
Proteomics data collection (LC-MS/MS)…… 30
Data analysis…… 31-34
16S amplicon sequencing data analysis…… 31
Proteomics data processing and analysis…… 31-33
References…… 34-35
NMR Spectra…… 36-51

### Supplementary Figures

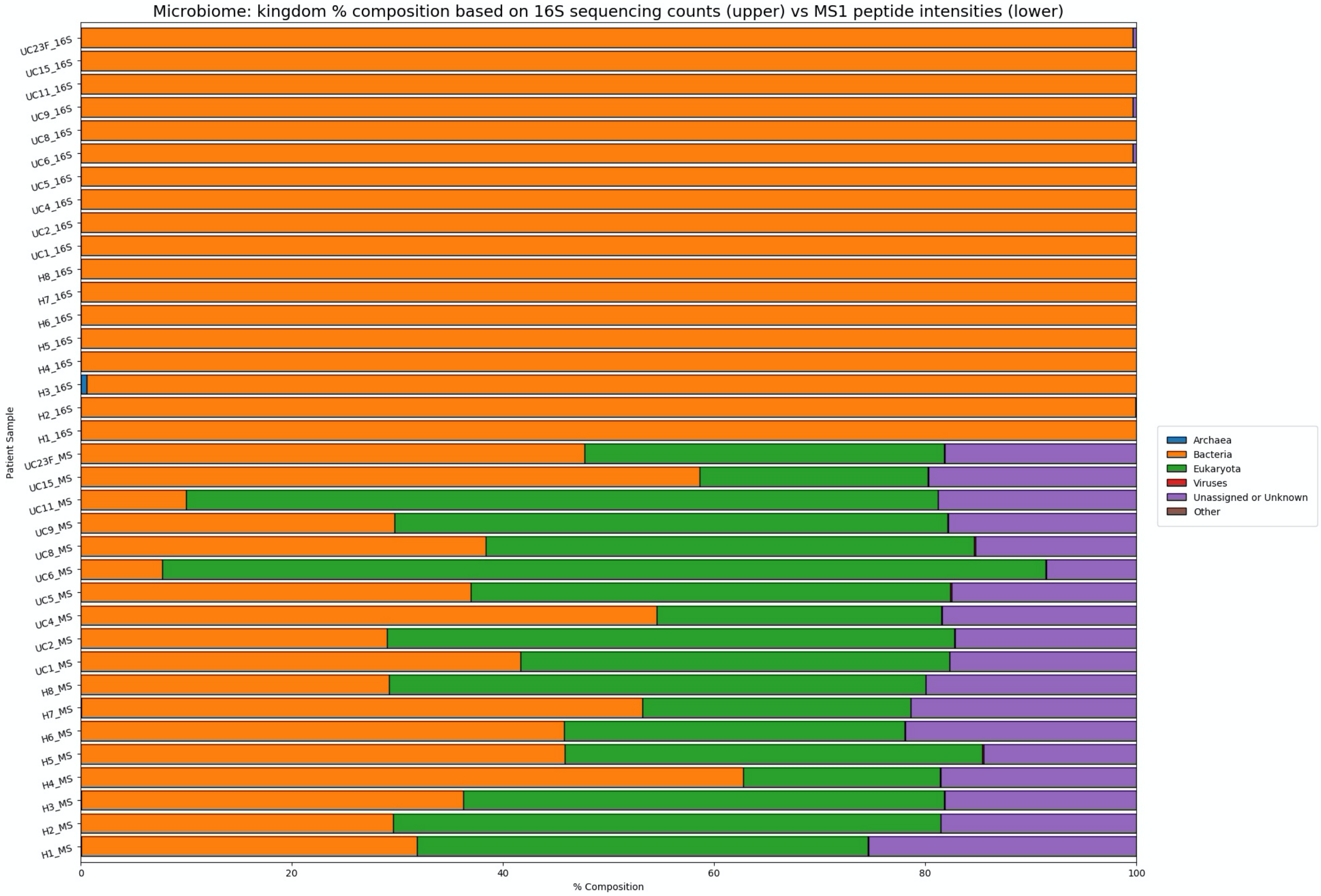


Figure S1. Microbiome taxonomy: kingdom level relative abundance by 16S amplicon sequencing counts (top 18 bars) versus LC-MS/MS peptide intensity means (lower 18 bars). Lowest abundance entries aggregated into “Other” category; for full deaggregated list, see **SI_A**.

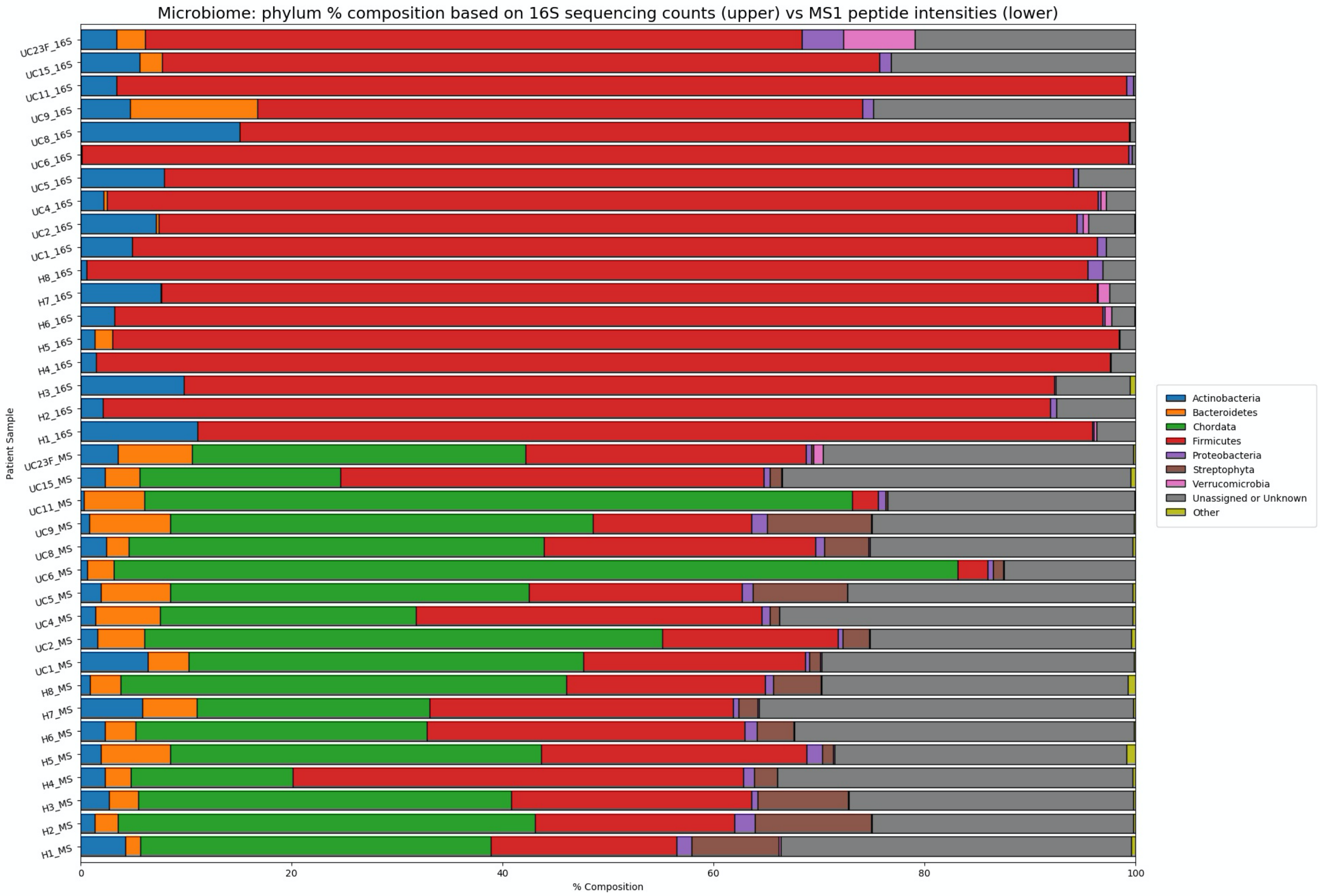


Figure S2. Microbiome taxonomy: phylum level relative abundance by 16S amplicon sequencing counts (top 18 bars) versus LC-MS/MS peptide intensity means (lower 18 bars). Lowest abundance entries aggregated into “Other” category; for full deaggregated list, see **SI_A**.

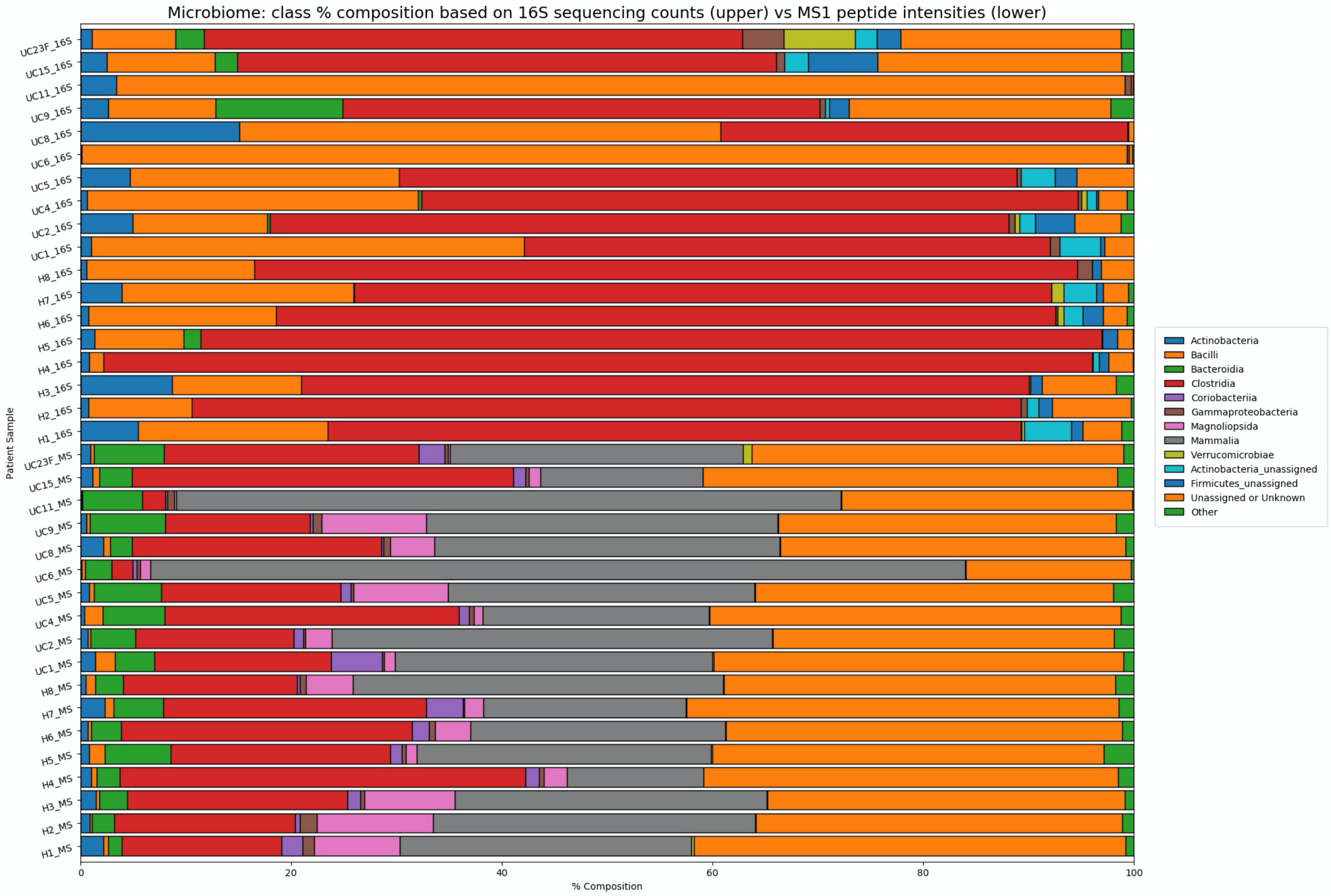


Figure S3. Microbiome taxonomy: class level relative abundance by 16S amplicon sequencing counts (top 18 bars) versus LC-MS/MS peptide intensity means (lower 18 bars). Lowest abundance entries aggregated into “Other” category; for full deaggregated list, see **SI_A**.

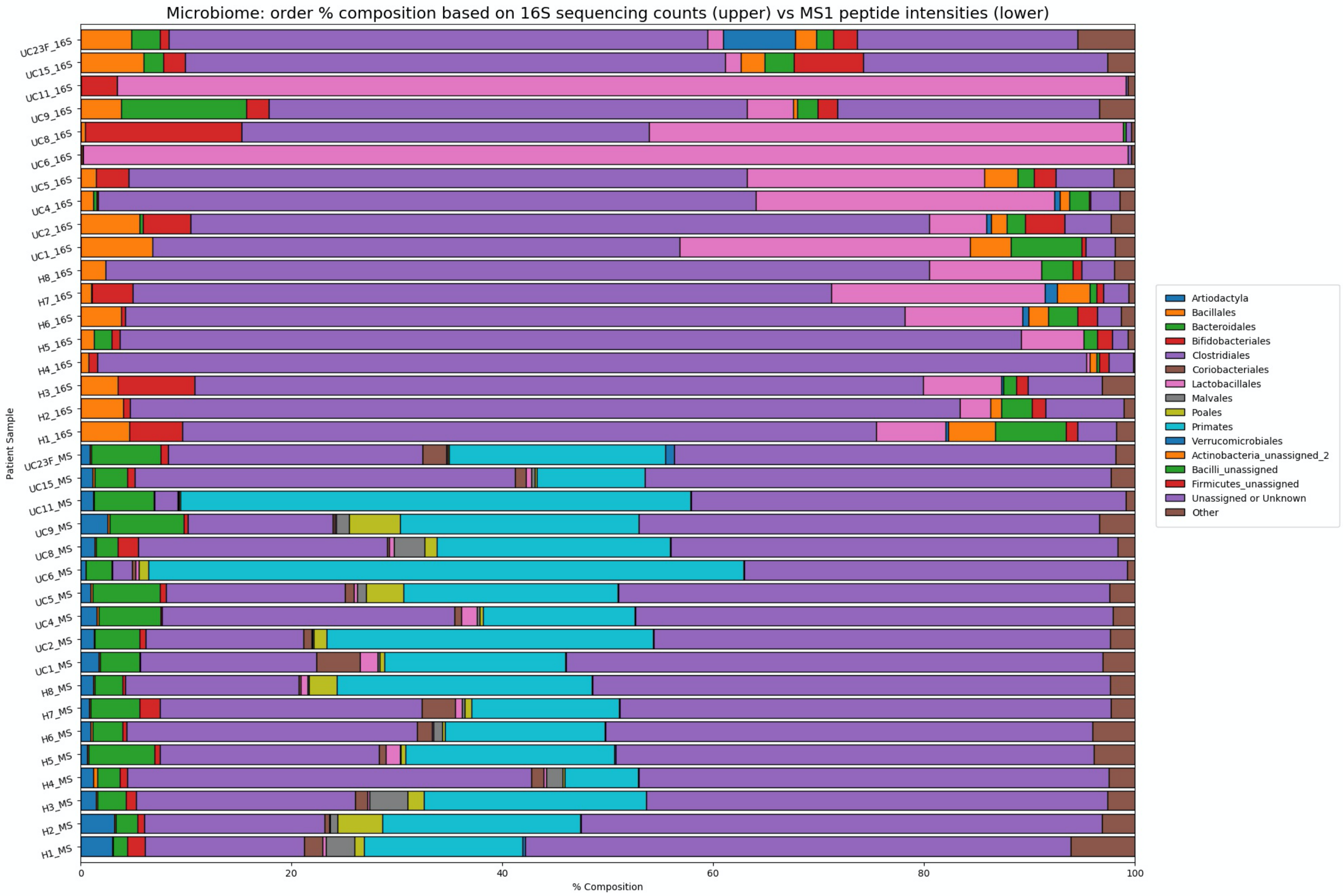


Figure S4. Microbiome taxonomy: order level relative abundance by 16S amplicon sequencing counts (top 18 bars) versus LC-MS/MS peptide intensity means (lower 18 bars). Lowest abundance entries aggregated into "Other" category; for full deaggregated list, see **SI_A**.

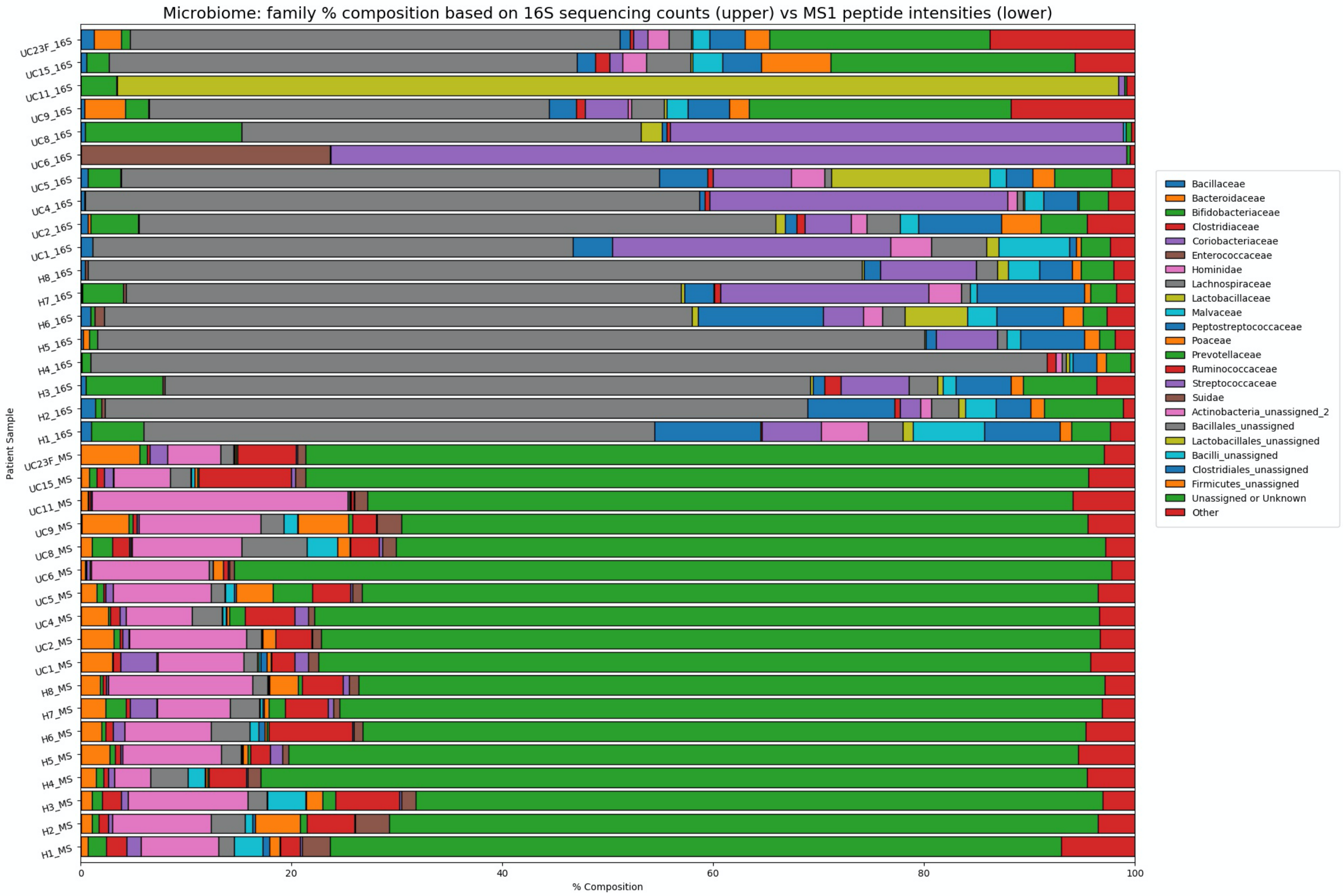


Figure S5. Microbiome taxonomy: family level relative abundance by 16S amplicon sequencing counts (top 18 bars) versus LC-MS/MS peptide intensity means (lower 18 bars). Lowest abundance entries aggregated into “Other” category; for full deaggregated list, see **SI_A**.

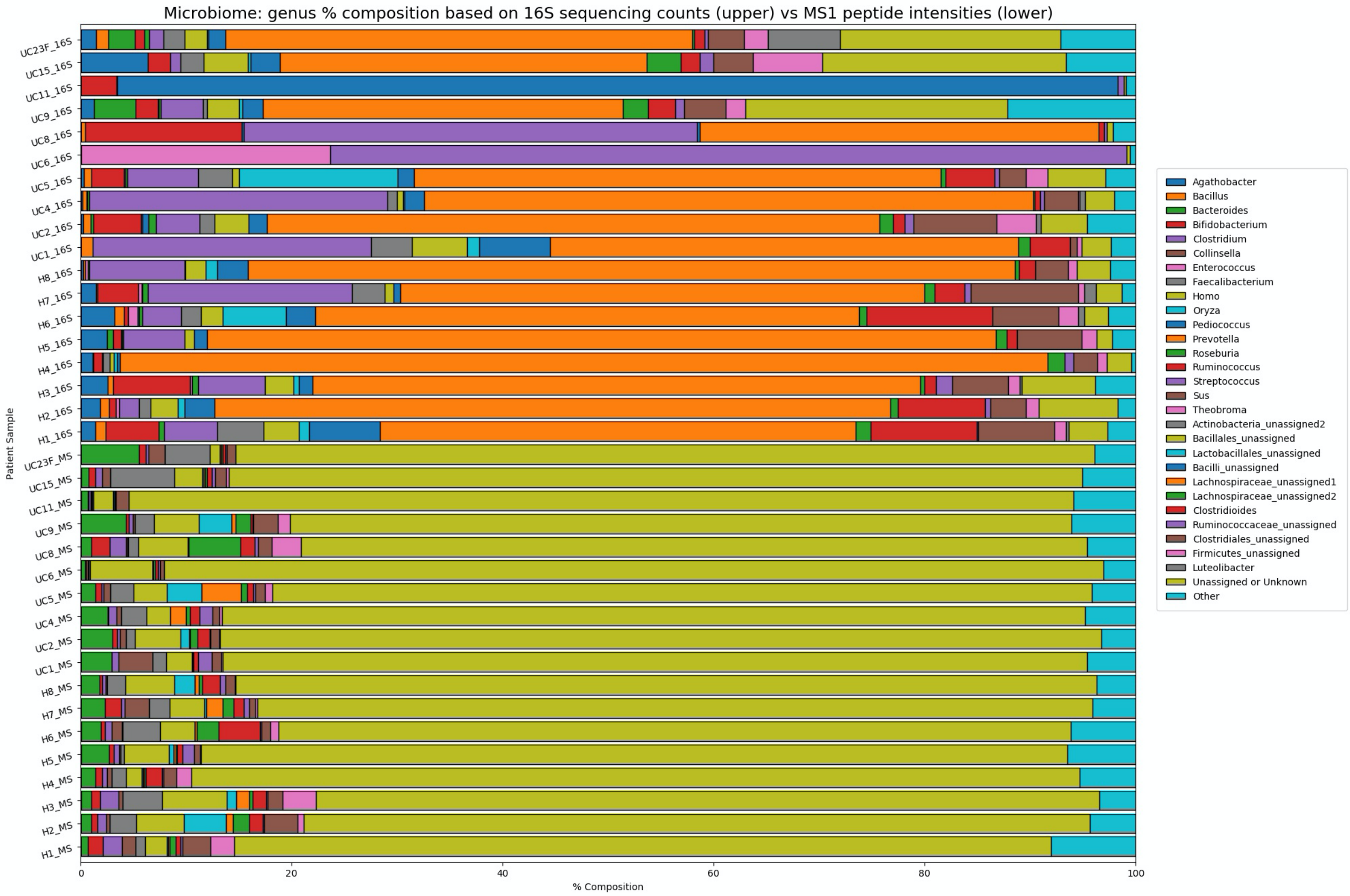


Figure S6. Microbiome taxonomy: genus level relative abundance by 16S amplicon sequencing counts (top 18 bars) versus LC-MS/MS peptide intensity means (lower 18 bars). Lowest abundance entries aggregated into “Other” category; for full deaggregated list, see **SI_A**.

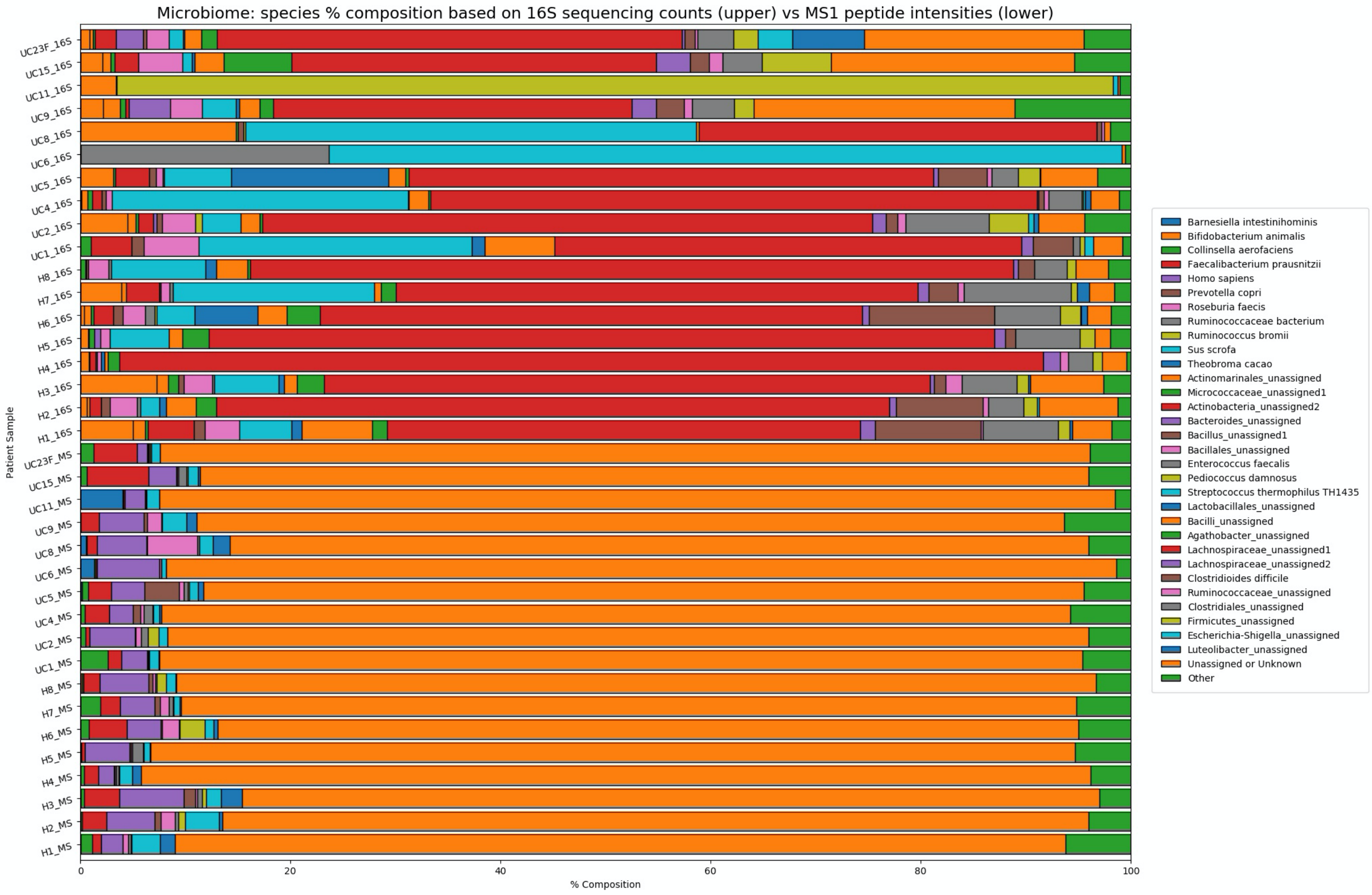


Figure S7. Microbiome taxonomy: species level relative abundance by 16S amplicon sequencing counts (top 18 bars) versus LC-MS/MS peptide intensity means (lower 18 bars). Lowest abundance entries aggregated into “Other” category; for full deaggregated list, see **SI_A**.

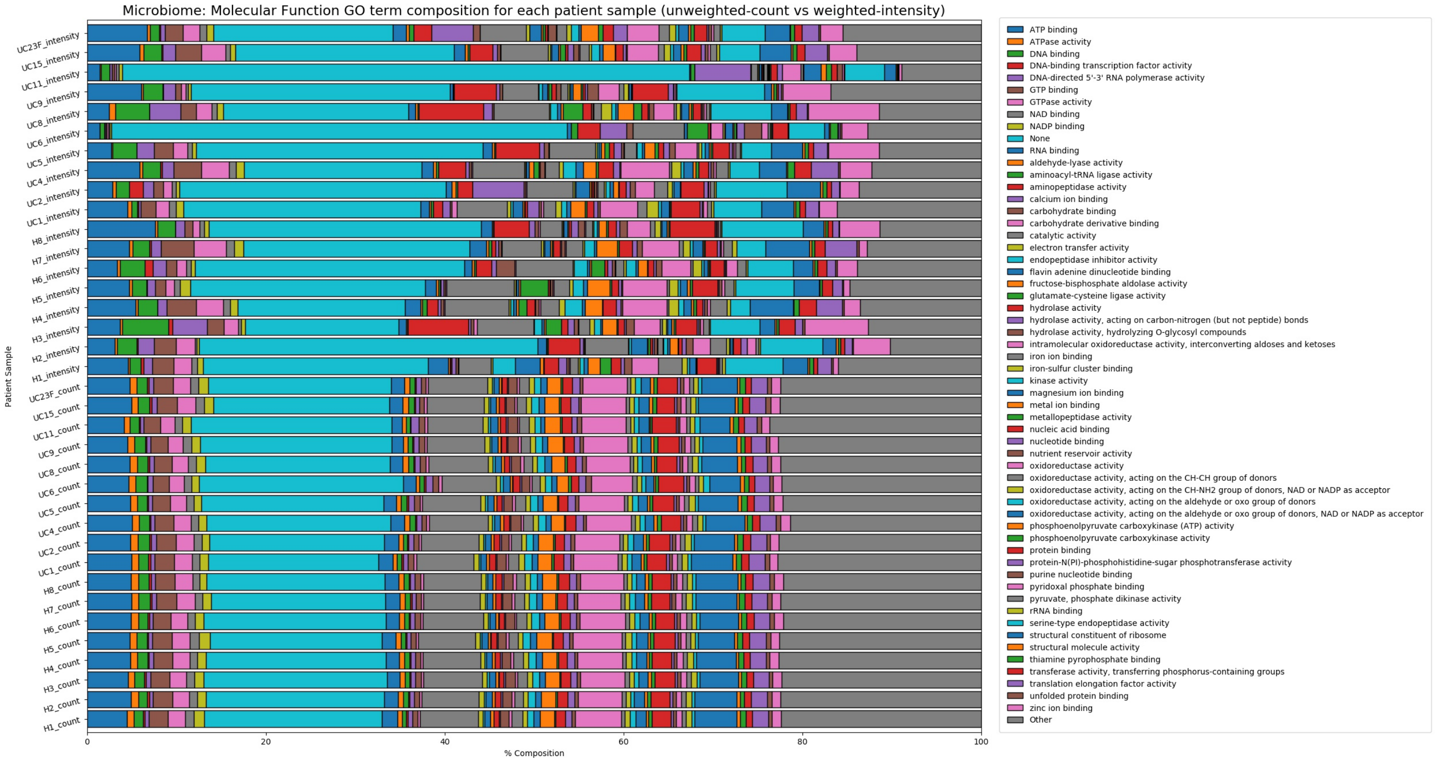


Figure S8. Microbiome molecular function GO term relative abundance: unweighted (lower 18 bars) versus weighted (upper 18 bars). For unweighted assembly, each protein group constituent contributes 1 count to an associated GO term; for weighted assembly, each protein group constituent contributes the average protein group peptide intensity to an associated GO term.

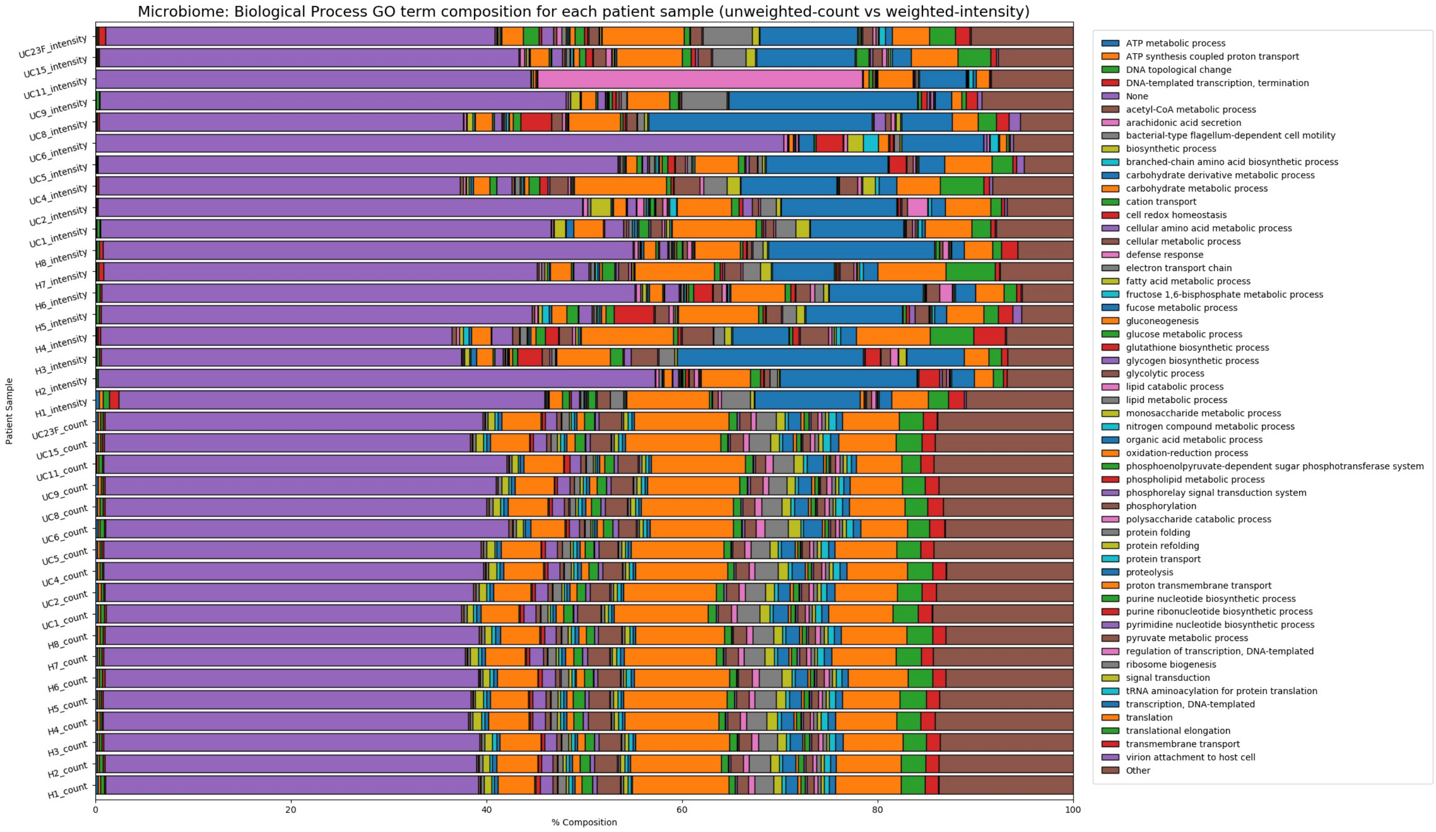


Figure S9. Microbiome biological process GO term relative abundance: unweighted (lower 18 bars) versus weighted (upper 18 bars). For unweighted assembly, each protein group constituent contributes 1 count to an associated GO term; for weighted assembly, each protein group constituent contributes the average protein group peptide intensity to an associated GO term.

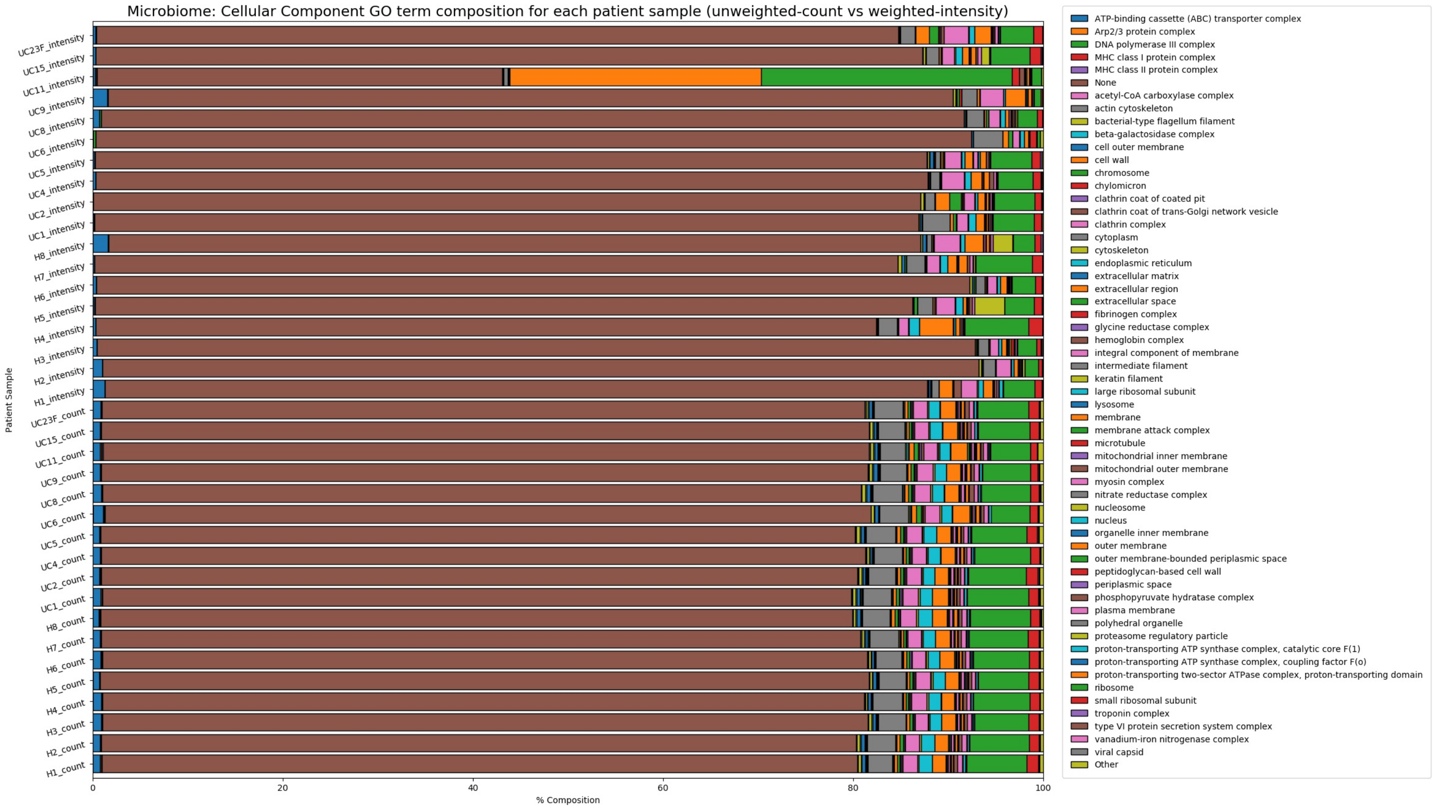


Figure S10. Microbiome cellular component GO term relative abundance: unweighted (lower 18 bars) versus weighted (upper 18 bars). For unweighted assembly, each protein group constituent contributes 1 count to an associated GO term; for weighted assembly, each protein group constituent contributes the average protein group peptide intensity to an associated GO term.

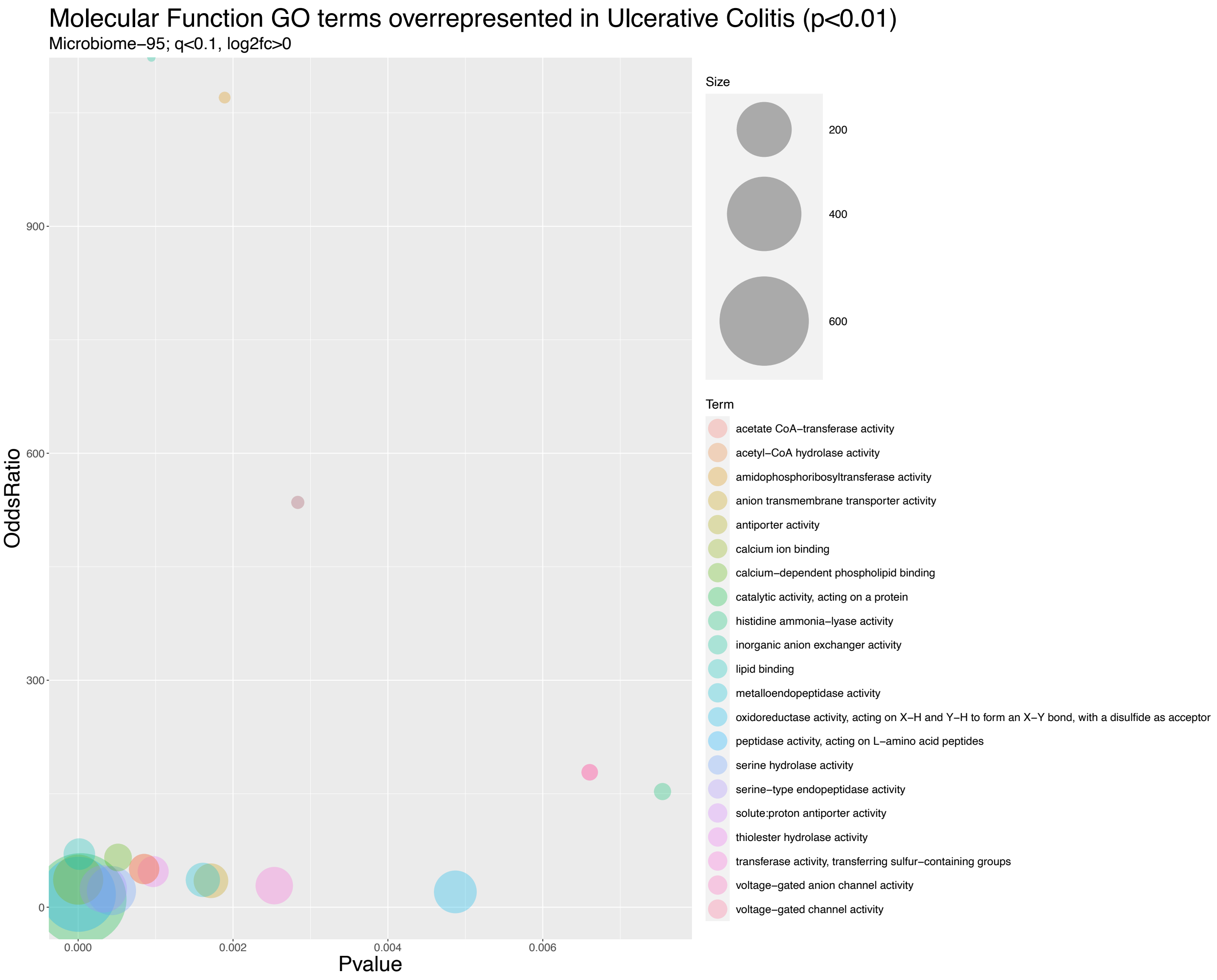


Figure S11. Bubble plot of molecular function GO terms enriched in ulcerative colitis patient fecal extracts ($p < 0.01$)

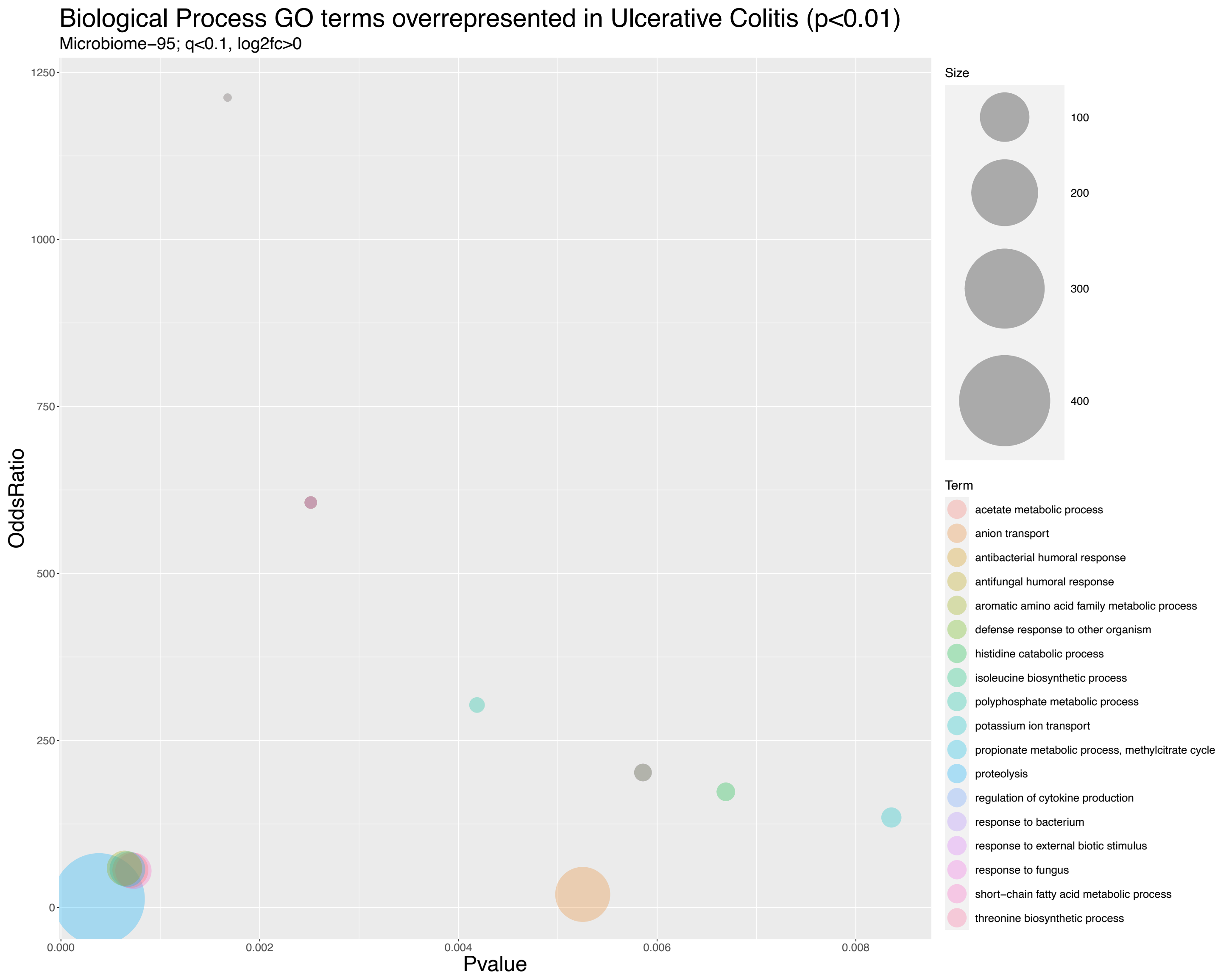


Figure S12. Bubble plot of biological process GO terms enriched in ulcerative colitis patient fecal extracts ($p < 0.01$)

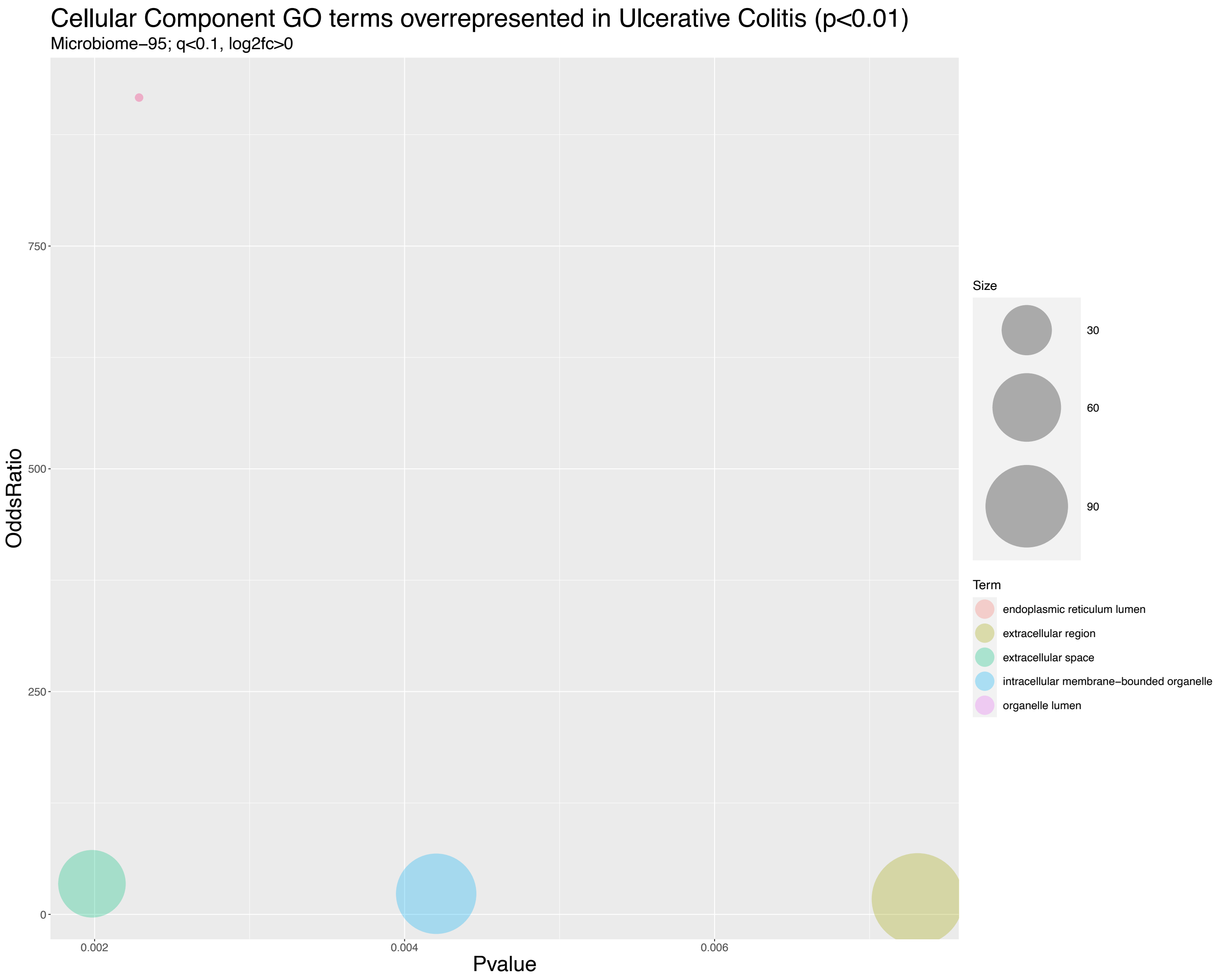


Figure S13. Bubble plot of cellular component GO terms enriched in ulcerative colitis patient fecal extracts ($p < 0.01$)

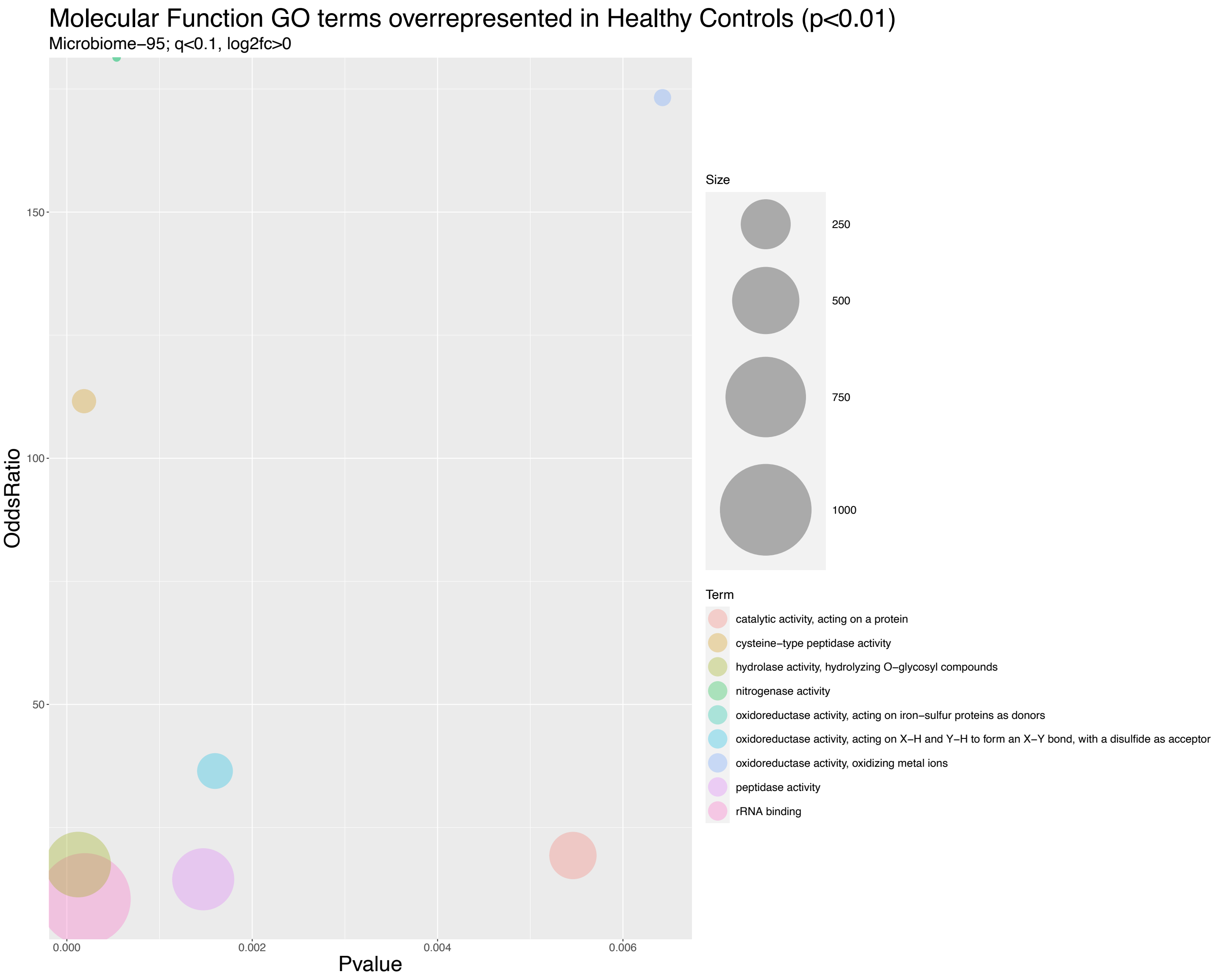


Figure S14. Bubble plot of molecular function GO terms enriched in healthy patient fecal extracts (*p* < 0.01)

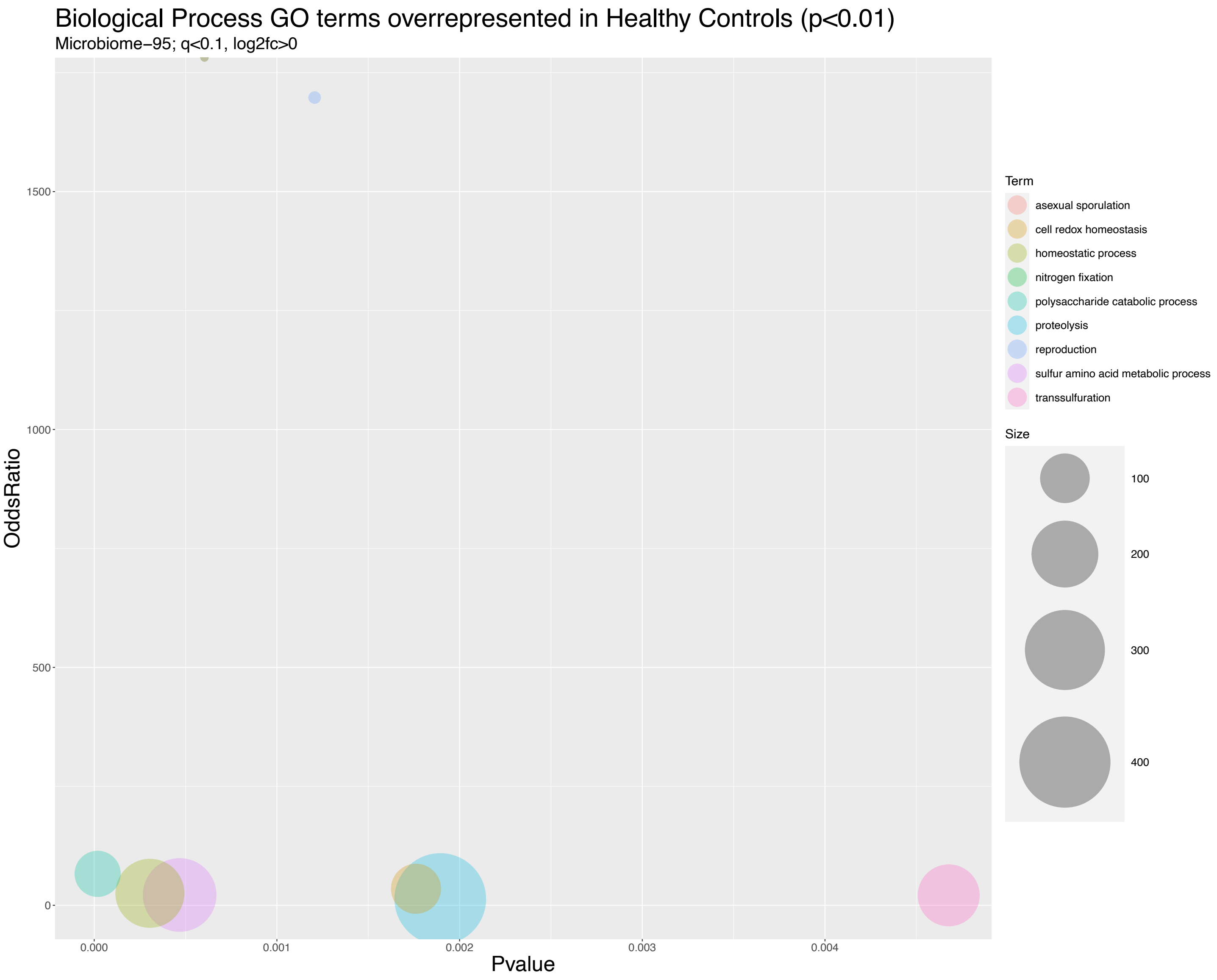


Figure S15. Bubble plot of biological process GO terms enriched in healthy patient fecal extracts ($p < 0.01$)

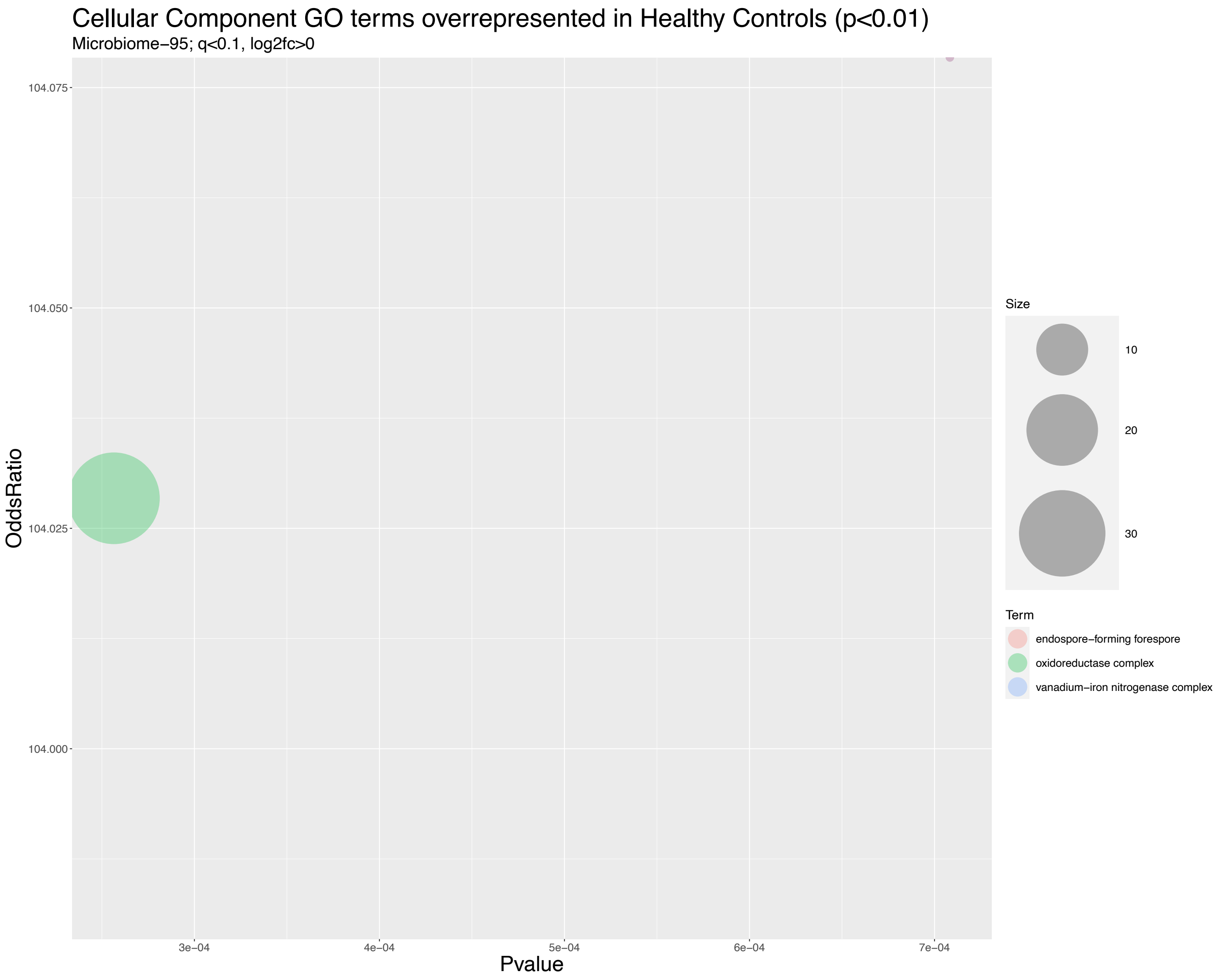


Figure S16. Bubble plot of cellular component GO terms enriched in healthy patient fecal extracts ($p < 0.01$)

| | # | GO ID | p-value | OddsRatio | Count | Size | Term |
|---|---|---|---|---|---|---|---|
| **Enriched in Ulcerative Colitis Patients** | | | | | | | |
| BP | 1 | GO:0006508 | 0.000383 | 12.70 | 4 | 405 | proteolysis |
| | 2 | GO:0009072 | 0.000639 | 58.74 | 2 | 44 | aromatic amino acid family metabolic process |
| | 3 | GO:0019679 | 0.000669 | 57.38 | 2 | 45 | propionate metabolic process, methylcitrate cycle |
| | 4 | GO:0006083 | 0.000699 | 56.07 | 2 | 46 | acetate metabolic process |
| | 5 | GO:0046459 | 0.000729 | 54.82 | 2 | 47 | short-chain fatty acid metabolic process |
| | 6 | GO:0009097 | 0.001677 | 1212.30 | 1 | 2 | isoleucine biosynthetic process |
| | 7 | GO:0009088 | 0.001677 | 1212.30 | 1 | 2 | threonine biosynthetic process |
| | 8 | GO:0019731 | 0.002515 | 606.14 | 1 | 3 | antibacterial humoral response |
| | 9 | GO:0019732 | 0.002515 | 606.14 | 1 | 3 | antifungal humoral response |
| | 10 | GO:0001817 | 0.002515 | 606.14 | 1 | 3 | regulation of cytokine production |
| | 11 | GO:0009620 | 0.002515 | 606.14 | 1 | 3 | response to fungus |
| | 12 | GO:0006797 | 0.004188 | 303.06 | 1 | 5 | polyphosphate metabolic process |
| | 13 | GO:0006820 | 0.005251 | 19.56 | 2 | 128 | anion transport |
| | 14 | GO:0043207 | 0.005858 | 202.04 | 1 | 7 | response to external biotic stimulus |
| | 15 | GO:0009617 | 0.005858 | 202.04 | 1 | 7 | response to bacterium |
| | 16 | GO:0098542 | 0.005858 | 202.04 | 1 | 7 | defense response to other organism |
| | 17 | GO:0006548 | 0.006692 | 173.17 | 1 | 8 | histidine catabolic process |
| | 18 | GO:0006813 | 0.008358 | 134.68 | 1 | 10 | potassium ion transport |
| MF | 1 | GO:0005509 | 4.842E-07 | 36.45 | 5 | 160 | calcium ion binding |
| | 2 | GO:0070011 | 2.634E-06 | 17.10 | 6 | 407 | peptidase activity, acting on L-amino acid peptides |
| | 3 | GO:0008289 | 1.544E-05 | 70.19 | 3 | 50 | lipid binding |
| | 4 | GO:0140096 | 3.249E-05 | 10.87 | 6 | 635 | catalytic activity, acting on a protein |
| | 5 | GO:0004252 | 0.000325 | 24.23 | 3 | 139 | serine-type endopeptidase activity |
| | 6 | GO:0017171 | 0.000430 | 21.96 | 3 | 153 | serine hydrolase activity |
| | 7 | GO:0005544 | 0.000515 | 65.73 | 2 | 35 | calcium-dependent phospholipid binding |
| | 8 | GO:0003986 | 0.000852 | 50.44 | 2 | 45 | acetyl-CoA hydrolase activity |
| | 9 | GO:0008775 | 0.000852 | 50.44 | 2 | 45 | acetate CoA-transferase activity |
| | 10 | GO:0005452 | 0.000946 | Inf | 1 | 1 | inorganic anion exchanger activity |
| | 11 | GO:0016790 | 0.000969 | 47.15 | 2 | 48 | thiolester hydrolase activity |
| | 12 | GO:0004222 | 0.001610 | 36.14 | 2 | 62 | metalloendopeptidase activity |
| | 13 | GO:0008509 | 0.001715 | 34.97 | 2 | 64 | anion transmembrane transporter activity |
| | 14 | GO:0004044 | 0.001892 | 1070.11 | 1 | 2 | amidophosphoribosyltransferase activity |
| | 15 | GO:0016782 | 0.002533 | 28.52 | 2 | 78 | transferase activity, transferring sulfur-containing groups |
| | 16 | GO:0015297 | 0.002836 | 535.05 | 1 | 3 | antiporter activity |
| | 17 | GO:0015299 | 0.002836 | 535.05 | 1 | 3 | solute:proton antiporter activity |
| | 18 | GO:0050485 | 0.004871 | 20.25 | 2 | 109 | oxidoreductase activity, acting on X-H and Y-H to form an X-Y bond, with a disulfide as acceptor |
| | 19 | GO:0022832 | 0.006606 | 178.34 | 1 | 7 | voltage-gated channel activity |
| | 20 | GO:0008308 | 0.006606 | 178.34 | 1 | 7 | voltage-gated anion channel activity |
| | 21 | GO:0004397 | 0.007546 | 152.86 | 1 | 8 | histidine ammonia-lyase activity |
| CC | 1 | GO:0005615 | 0.001983 | 34.36 | 2 | 58 | extracellular space |
| | 2 | GO:0043233 | 0.002287 | 916.60 | 1 | 2 | organelle lumen |
| | 3 | GO:0005788 | 0.002287 | 916.60 | 1 | 2 | endoplasmic reticulum lumen |
| | 4 | GO:0043231 | 0.004204 | 23.15 | 2 | 85 | intracellular membrane-bounded organelle |
| | 5 | GO:0005576 | 0.007309 | 17.28 | 2 | 113 | extracellular region |

Figure S17. All enriched GO terms from ulcerative colitis patient fecal samples identified by GOstats ($p < 0.01$)

| Enriched in Healthy Patients | | | | | | | |
|---|---|---|---|---|---|---|---|
| | # | GO ID | p-value | OddsRatio | Count | Size | Term |
| BP | 1 | GO:0000272 | 1.878E-05 | 66.11 | 3 | 84 | polysaccharide catabolic process |
| | 2 | GO:0042592 | 0.000305 | 25.21 | 3 | 215 | homeostatic process |
| | 3 | GO:0000096 | 0.000468 | 21.71 | 3 | 249 | sulfur amino acid metabolic process |
| | 4 | GO:0009399 | 0.000603 | Inf | 1 | 1 | nitrogen fixation |
| | 5 | GO:0030436 | 0.000603 | Inf | 1 | 1 | asexual sporulation |
| | 6 | GO:0000003 | 0.001206 | 1697.63 | 1 | 2 | reproduction |
| | 7 | GO:0045454 | 0.001761 | 34.77 | 2 | 102 | cell redox homeostasis |
| | 8 | GO:0006508 | 0.001894 | 13.26 | 3 | 405 | proteolysis |
| | 9 | GO:0019346 | 0.004677 | 20.93 | 2 | 168 | transsulfuration |
| MF | 1 | GO:0004553 | 0.000122 | 17.47 | 4 | 472 | hydrolase activity, hydrolyzing O-glycosyl compounds |
| | 2 | GO:0008234 | 0.000185 | 111.59 | 2 | 37 | cysteine-type peptidase activity |
| | 3 | GO:0019843 | 0.000195 | 10.48 | 5 | 1000 | rRNA binding |
| | 4 | GO:0016730 | 0.000537 | Inf | 1 | 1 | oxidoreductase activity, acting on iron-sulfur proteins as donors |
| | 5 | GO:0016163 | 0.000537 | Inf | 1 | 1 | nitrogenase activity |
| | 6 | GO:0008233 | 0.001471 | 14.48 | 3 | 416 | peptidase activity |
| | 7 | GO:0050485 | 0.001598 | 36.47 | 2 | 109 | oxidoreductase activity, acting on X-H and Y-H to form an X-Y bond, with a disulfide as acceptor |
| | 8 | GO:0140096 | 0.005460 | 19.31 | 2 | 219 | catalytic activity, acting on a protein |
| | 9 | GO:0016722 | 0.006427 | 173.26 | 1 | 12 | oxidoreductase activity, oxidizing metal ions |
| CC | 1 | GO:1990204 | 0.000257 | 104.03 | 2 | 34 | oxidoreductase complex |
| | 2 | GO:0016613 | 0.000708 | Inf | 1 | 1 | vanadium-iron nitrogenase complex |
| | 3 | GO:0042601 | 0.000708 | Inf | 1 | 1 | endospore-forming forespore |

Figure S18. All enriched GO terms from healthy patient fecal samples identified by GOstats (*p* < 0.01)

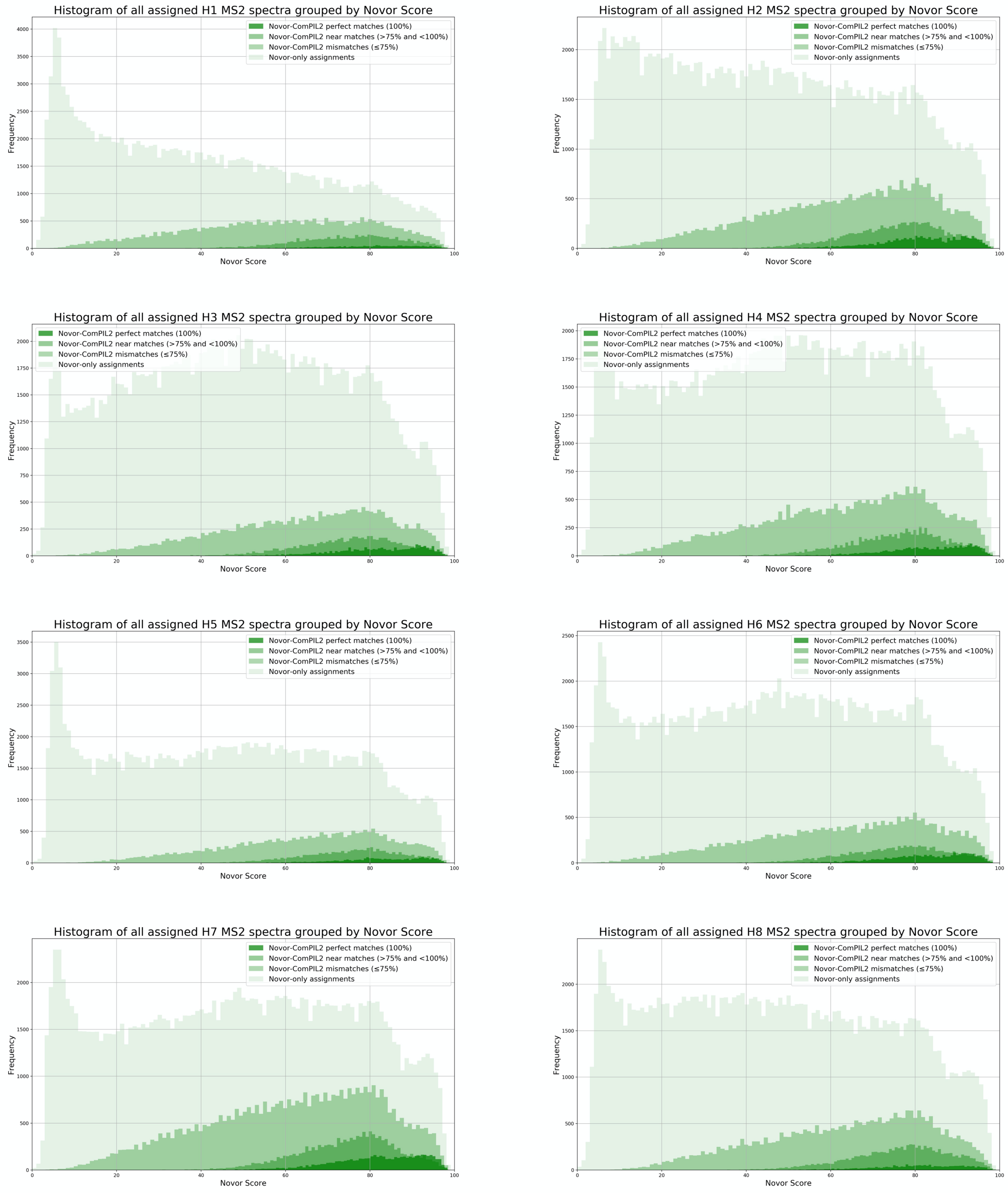


Figure S19. Novor score histograms for healthy patient fecal microbiomes.

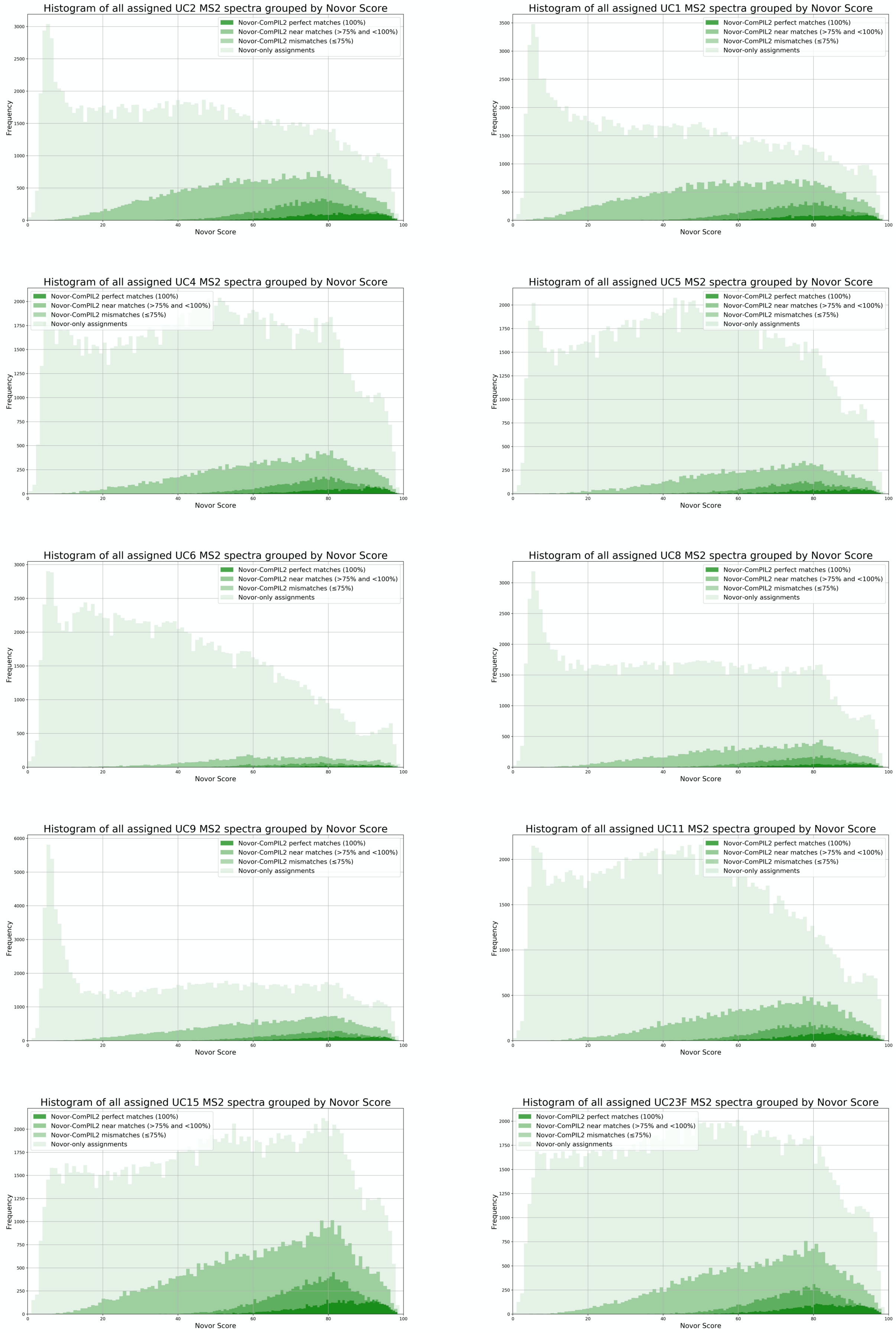


Figure S20. Novor score histograms for UC patient fecal microbiomes.

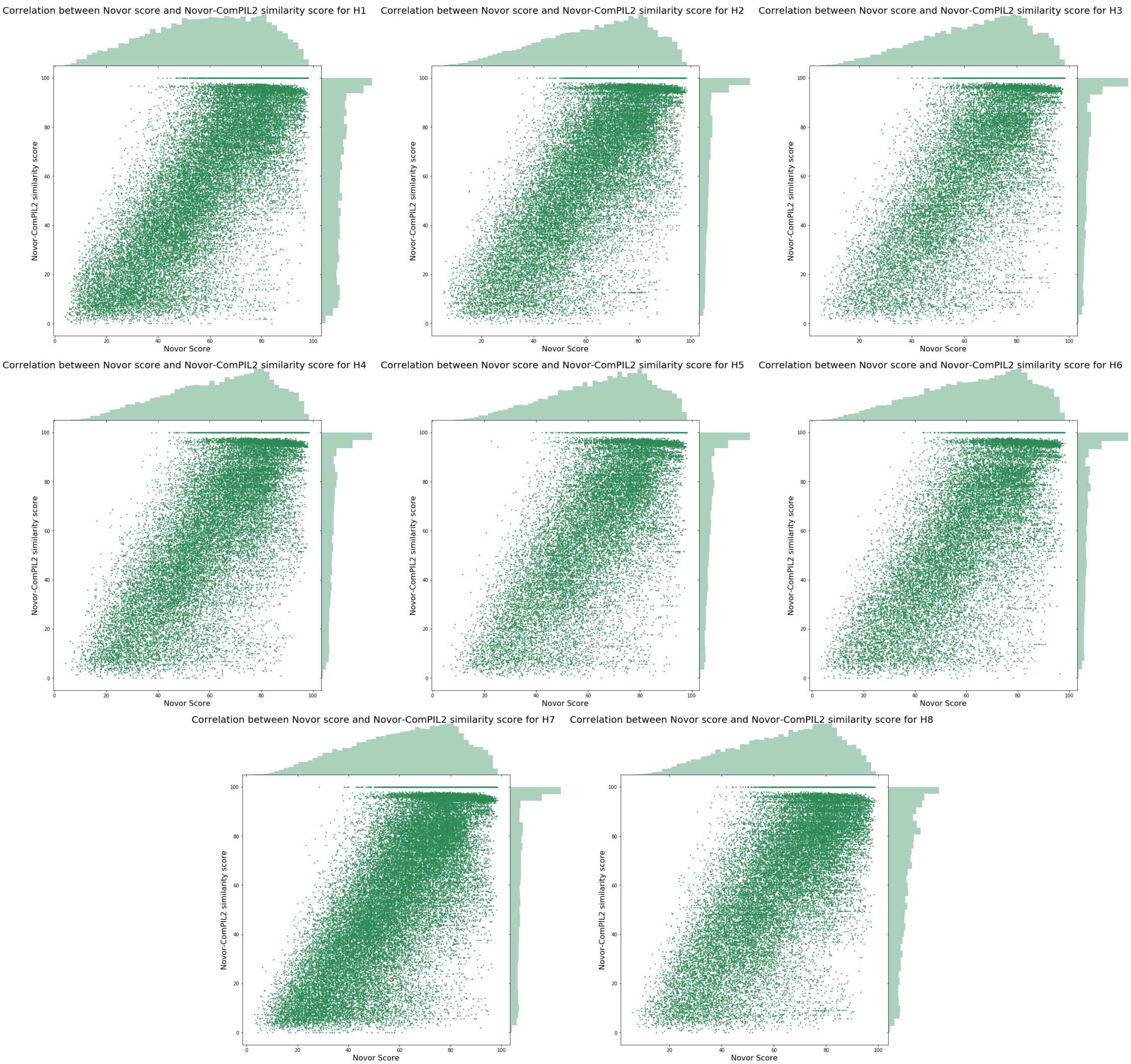


Figure S21. Novor score vs. Novor-ComPIL2 similarity score joint plots for healthy patient fecal microbiomes.

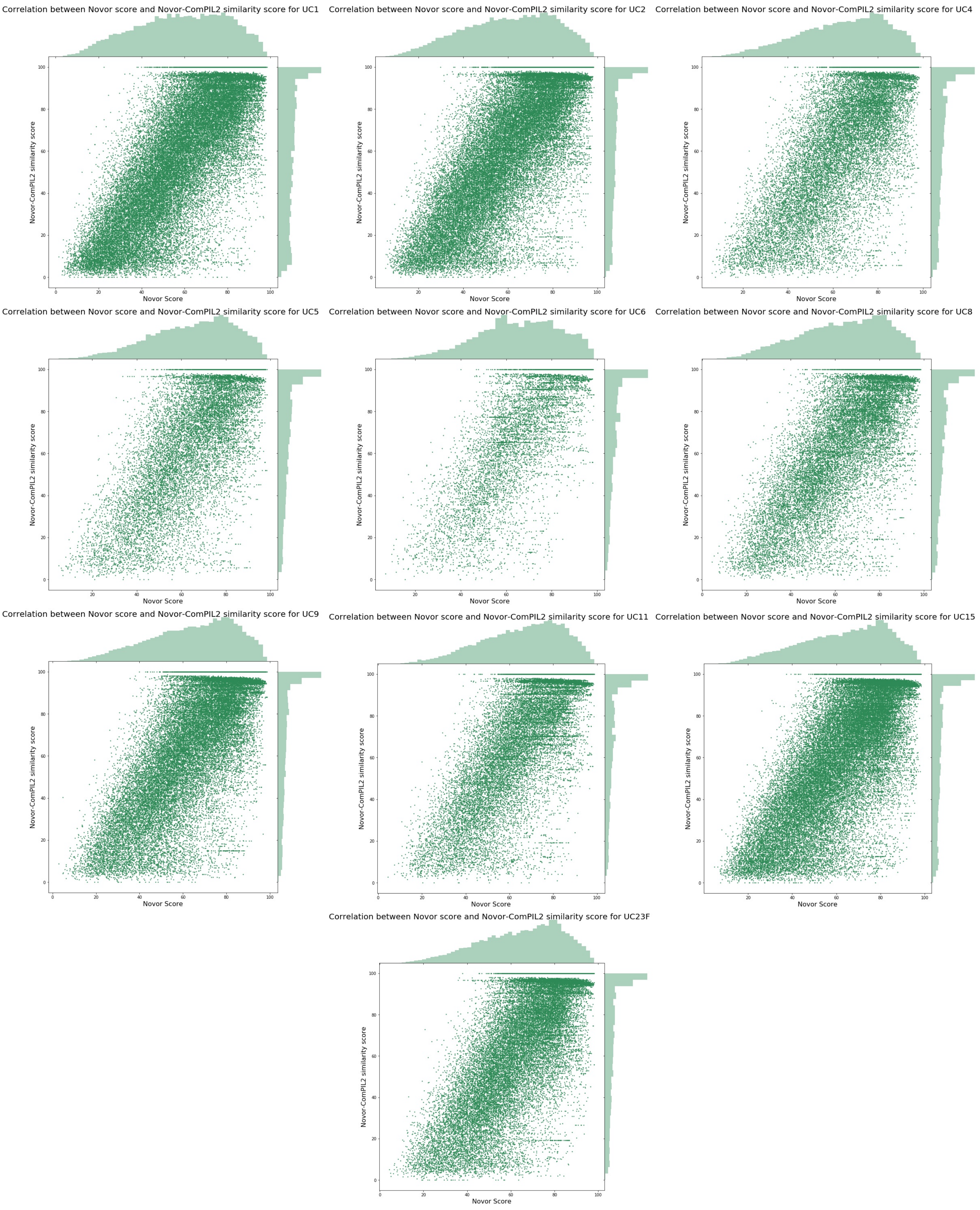


Figure S22. Novor score vs. Novor-ComPIL2 similarity score joint plots for ulcerative colitis patient fecal microbiomes.

### Methods

#### ***General information, reagent sourcing, and instrumentation information***

*Reagents and Consumables:* Unless otherwise noted, all materials were used as received from commercial sources without further purification. Urea was purchased from VWR (Radnor, PA, USA). Biotin was purchased from Combi-blocks (San Diego, CA, USA). Diethyl vinyl phosphonate, mesyl chloride and sodium hydride were purchased from Alfa Aesar (Haverhill, MA, USA). 2-chloracetamide and trifluoromethanesulfonic acid (triflic acid) were purchased from TCI (Tokyo, Japan). Palladium on carbon, iodine, tetraethylene glycol, sodium dodecyl sulfate, and calcium chloride were purchased from Sigma-Aldrich (St. Louis, MO, USA). EDC•HCl was purchased from Chem Implex (Wood Dale, IL, USA). Diethylaminosulfur trifluoride (DAST) was purchased from Oakwood (Estill, SC, USA). Hydrogen gas was purchased from Praxair (Danbury, CT, USA). Sequencing grade trypsin (V5111) was purchased from Promega, (Madison, WI, USA). Bicine, triethylamine, tris•HCl, pyridine, lithium azide, and 2-mercaptoethanol were purchased from Acros (Fair Lawn, NJ, USA). Formic acid (>99% purity, ampules), dimethyl sulfoxide, sodium azide, PBS (10X), TBS (10X), high capcity streptavidin agarose, TCEP•HCl, BCA Protein Assay Kit, and Micro BCA Protein Assay Kit were purchased from ThermoFisher (Waltham, MA, USA). Fecal/soil DNA extraction kit was purchased from Zymo Research (Irvine, CA, USA). ZipTip C18 pipette tips, 3K and 10K molecular weight cut-off (MWCO) centrifugal concentrators, and silica gel 60 $F_{254}$ plates (for thin layer chromatography) were purchased from MilliporeSigma (Burlington, MA, USA). Deuterated solvents for NMR were purchased from Cambridge Isotope Laboratories (Tewksbury, MA, USA). Laemmli Sample Buffer, running buffer, transfer buffer, and Mini-PROTEAN TGX Gels were purchased from Biorad (Hercules, CA, USA). IRDye streptavidin was purchased from LiCOR (Lincoln, NE, USA). Nylon mesh cell strainers were purchased from Corning (Corning, NY, USA).

*Equipment:* All synthetic reactions were run on stirring hot plates (IKA, Staufen, German) equipped with oil baths calibrated to an external thermometer. Prior to beginning an experiment, the hot plate was turned on, and the oil bath was allowed to equilibrate to the desired temperature for at least 30 minutes. $^{1}H$ and $^{13}C$ NMR spectra were recorded on a DRX (Bruker, Billerica, MA, USA) equipped with a 5mm DCH cryoprobe ($^{1}H$: 600 MHz and $^{13}C$: 151 MHz) instrument internally referenced to tetramethylsilane (TMS) or chloroform ($^{1}H$ NMR signals referenced to 0.0 ppm-TMS; $^{13}C$ NMR signal referenced to 77.16 ppm-$CDCl_3$, 49.00 ppm-MeOD, 39.52 ppm-DMSO-$d_6$) signals unless otherwise noted. $^{31}P$ NMR spectra were recorded on a DPX ($^{1}H$: 400 MHz and $^{31}P$: 162 MHz) (Bruker, Billerica, MA, USA). The following abbreviations (or combinations thereof) were used to explain multiplicities: s = singlet, d = doublet, t = triplet, q = quartet, p = pentet, m = multiplet, comp. = complex, br = broad, and app. = apparent. Several compounds were purified via a preparative HPLC system using a 19x100mm or 19x150mm C18 5 μm OBD column (Waters, Milford, MA, USA). High resolution mass spectra for compound characterization were recorded at the Center for Mass Spectrometry, Scripps Research. Low resolution mass spectra for compound characterization were recorded using a MSQ single-quadrapole mass spectrometer (ThermoFisher, Waltham, MA, USA). Microbiome cells were lysed using a Q700 sonicator with cup horn attachment (QSonica, Newtown, CT, USA). Protein and nucleic acid concentrations were measured with a NanoDrop spectrophotometer (ThermoFisher, Waltham, MA, USA). Electrophoresis and transfer were performed using Mini Trans-Blot equipment (Biorad, Hercules, CA, USA). Fluorescence assays were performed using an Envision 2104 Multilabel Reader (Perkin Elmer, Waltham, MA, USA). Fluorescence-based western blots were recorded using Odyssey instruments (LI-COR, Lincoln, NE, USA). Proteomics mass spectrometry data were collected using an Orbitrap Fusion Tribrid instrument equipped with a nEASY-LC1000 and a nanospray Flex ion source (ThermoFisher, Waltham, MA, USA).

### ***Chemical Synthesis***

#### *Synthetic scheme for FP-Probe*

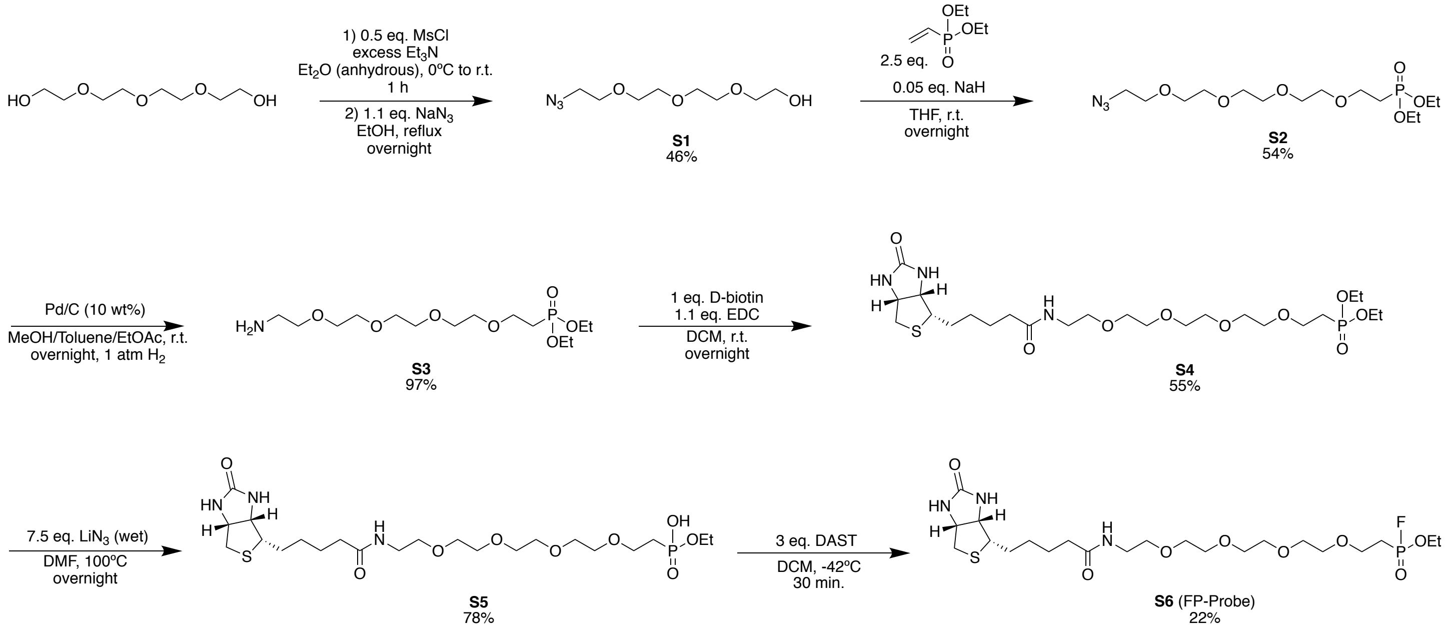

*Compound preparation procedures and characterization*

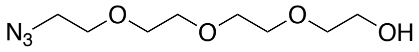

S1

**2-(2-(2-(2-azidoethoxy)ethoxy)ethoxy)ethan-1-ol (S1)** (1)
In a flame-dried round bottom flask equipped with stir bar, tetraethyleneglycol (7.5 g, 38.6 mmol) was dissolved in anhydrous diethyl ether (75 mL). Anhydrous triethylamine (7.5 mL) was added and the solution was chilled to 0ºC. Mesyl chloride (2.21 g, 19.3 mmol) was diluted in anhydrous diethyl ether (10 mL) then added dropwise to the vigorously stirring solution of tetrathetyleneglycol at 0ºC over 30 minutes. The mixture stirred at 0ºC for 30 minutes before warming to room temperature. Solvent was removed *in vacuo* to yield a viscous pale yellow oil which was used directly for the next step. The pale yellow oil was dissolved in ethanol (150 mL) then sodium azide (2.76 g, 42.5 mmol) was added to the stirring solution. The resulting suspension was refluxed overnight before solvent was removed *in vacuo* to yield an oily solid. The oily solid was resuspended in ethyl acetate and filtered over Celite. The filtrate was condensed and purified by silica gel flash chromatography using an ethyl acetate/hexanes gradient (50:50 → 100:0) to yield **S1** (1.94 g, 8.8 mmol, 46%).
**$R_f$**: 0.12 (EtOAc, visualized with CAM stain)
**$^1$H NMR (600 MHz, $CDCl_3$):** δ 3.73 (br m, 2H), 3.68 (m, 10 H), 3.62 (m, 2H), 3.40 (t, $J$ = 5.1 Hz, 2H).
**$^{13}$CNMR (151 MHz, $CDCl_3$):** δ 72.59, 70.81, 70.77, 70.70, 70.45, 70.15, 61.82, 50.79.
**LC-MS (ESI)**: 220.15 $[M+H]^+$
**HRMS (ESI-TOF):** m/z calculated for $C_8H_{17}N_3O_4$ $[M+H]^+$: 220.1292, found: 220.1290.

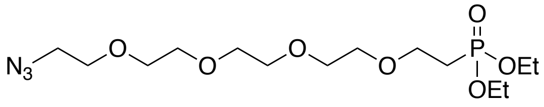

S2

**diethyl (14-azido-3,6,9,12-tetraoxatetradecyl)phosphonate (S2)**
In a round bottom flask equipped with stir bar, **S1** (500 mg, 2.28 mmol) and diethyl vinylphosphonate (936 mg, 5.70 mmol) were combined in anhydrous tetrahydrofuran (10mL). Sodium hydride (4.8 mg, 0.114 mmol) was suspended in anhydrous THF (1mL), then added to the stirring solution of **S1** and diethyl vinylphosphonate at room temperature while stirring vigorously. The reaction was stirred overnight at room temperature then dried in vacuo to yield a viscous oil. The oil was purified by silica gel flash chromatography using an Ethyl acetate/hexanes/methanol gradient (50:50:0 → 100:0:0 → 95:0:5 → 90:0:10) to yield **S2** (476 mg, 1.24 mmol, 54%) as a colorless oil.
**$R_f$**: 0.91 (EtOAc, visualized with CAM)
**$^1$H NMR (600 MHz, $CDCl_3$):** δ 4.06-4.13 (m, 4H), 3.60-3.75 (m, 16H), 3.40 (t, $J$ = 5.1 Hz, 2H), 2.13 (m, 2H), 1.32 (t, $J$ = 7.1 Hz, 6H).
**$^{13}$CNMR (151 MHz, $CDCl_3$):** δ 70.72, 70.67, 70.64, 70.59, 70.47, 70.25, 70.16, 65.19 (d, $J_{C-P}$ = 1.5 Hz), 61.93 (d, $J_{C-P}$ = 6.1 Hz), 50.77, 26.94 (d, $J_{C-P}$ = 139.9 Hz), 16.53 (d, $J_{C-P}$ = 6.1 Hz).
**$^{31}$PNMR (162 MHz, $CDCl_3$):** δ 29.16 (Note that 151.07 signal is an artifact).
**LC-MS (ESI-Q)**: 384.08 $[M+H]^+$
**HRMS (ESI-TOF):** m/z calculated for $C_{14}H_{30}N_3O_7P$ $[M+Na]^+$: 406.1713, found: 406.1723.

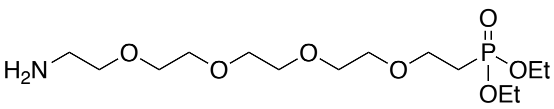

S3

**diethyl (14-amino-3,6,9,12-tetraoxatetradecyl)phosphonate (S3)**
Azide **S2** (589 mg, 1.54 mmol) was dissolved in methanol/ethyl acetate (50:50; 5 mL) and charged into a round bottom flask equipped with stir bar. A toluene suspension of palladium on carbon was slowly added to the stirring azide solution at room temperature. After addition, the flask was sealed with a septum, purged of air, then backfilled with hydrogen via balloon. This purge and backfilling process was repeated for a total of 3 cycles. After the third cycle, the slurry was allowed to stir under a hydrogen atmosphere for 3 hours at room temperature before purging and backfilling one last time with hydrogen. The resulting slurry was stirred overnight at room temperature and reaction progress was monitored by TLC (staining with ninhydrin). After starting material was completely consumed, the flask was open to air and solvent removed *in vacuo*. The crude residue was dissolved in ethyl acetate, and filtered through a 22 $\mu$m syringe filter and dried again to yield **S3** (532 mg, 1.49 mmol, 97%) as a relatively pure colorless oil. For synthetic applications, no further purification was applied. Analytically pure **S3** was obtained by preparative HPLC (19x100 mm C18) using a $H_2O$/MeCN (0.1% TFA) gradient (80:20 → 20:80).
**$R_f$**: 0.46 (TFA salt; EtOAc, visualized with ninhydrin)
**$^1$H NMR (600 MHz, $CDCl_3$):** δ 4.09 (m, 4H), 3.81 (m, 2H), 3.58-3.75 (m, 14H), 3.20 (br m, 2H), 2.12 (dt, $J$ = 17.9, 6.2 Hz, 2H), 1.32 (t, $J$ = 7.1 Hz, 6H).
**$^{13}$CNMR (151 MHz, $CDCl_3$):** δ 70.42, 70.11, 70.04, 70.03, 69.86, 69.28, 66.81, 65.12 (d, $J_{C-P}$ = 7.2 Hz), 62.49 (d, $J_{C-P}$ = 6.6 Hz), 39.46, 26.26 (d, $J_{C-P}$ = 142.1 Hz), 16.38 (d, $J_{C-P}$ = 6.1 Hz).

**$^{31}$PNMR (162 MHz, $CDCl_3$):** δ 30.51 (Note that 151.07 signal is an artifact).
**LC-MS (ESI-Q)**: 358.16 $[M+H]^+$
**HRMS (ESI-TOF):** m/z calculated for $C_{14}H_{32}NO_7P$ $[M+H]^+$: 358.1989, found: 358.1996.

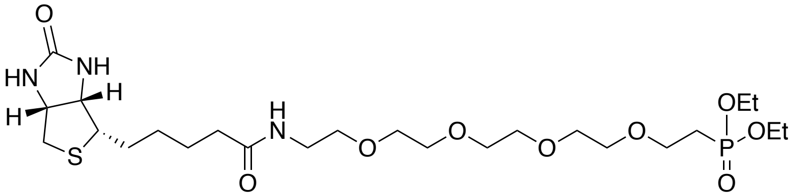

**S4**

**diethyl (16-oxo-20-((3a*S*,4*S*,6a*R*)-2-oxohexahydro-1*H*-thieno[3,4-*d*]imidazol-4-yl)-3,6,9,12-tetraoxa-15-azaicosyl)phosphonate (S4)**
**S3** (532 mg, 1.49 mmol) was dissolved in dichloromethane (5 mL) and charged into a round bottom flask equipped with stir bar. D-Biotin (364 mg, 1.49 mmol) was added to the solution to yield a white suspension. A dichloromethane solution of EDC•HCl (314 mg, 1.64 mmol) was added dropwise to the suspension yielding a more soluble slurry over 10 min. After stirring overnight at room temperature, solvent was removed *in vacuo*. Crude product was purified by silica gel flash chromatography using a dichloromethane/methanol (90:10 → 80:10) gradient to yield **S4** (476 mg, 0.815 mmol, 55%) as a white waxy solid after drying.
**$R_f$**: 0.64 (EtOAc, visualized with CAM)
**$^1$H NMR (600 MHz, $CDCl_3$):** δ 7.07 (br s, 1H), 6.87 (br s, 1H), 4.58 (dd, *J* = 8.0, 4.2 Hz, 1H), 4.39 (dd, *J* = 8.0, 4.5 Hz, 1H), 4.11 (m, 4H), 3.73 (m, 2H), 3.64 (m, 13H), 3.56 (t, *J* = 4.6 Hz, 2H), 3.45 (m, 2H), 3.19 (td, *J* = 7.4, 4.5 Hz, 1H), 2.94 (dd, *J* = 13.0, 4.9 Hz, 1H), 2.78 (d, *J* = 12.8 Hz, 1H), 2.27 (td, *J* = 7.3, 4.1, 2H), 2.14 (m, 2H), 1.69 (m, 4H), 1.46 (m, 2H), 1.32 (t, *J* = 7.1 Hz, 6H).
**$^{13}$CNMR (151 MHz, $CDCl_3$):** δ 174.15, 164.81, 70.53, 70.31, 70.48, 70.39, 70.20, 70.13, 69.92, 65.03, 62.29 (d, $J_{C\text{-}P}$ = 1.7 Hz), 62.25, 60.87, 55.37, 40.48, 39.44, 35.53, 27.90 (d, $J_{C\text{-}P}$ = 8.8 Hz), 26.80 (d, $J_{C\text{-}P}$ = 140.4 Hz), 25.50, 16.47 (d, $J_{C\text{-}P}$ = 6.1 Hz) (Note that 1 peak is occluded but appears in **S5** and **S6**).
**$^{31}$PNMR (162 MHz, $CDCl_3$):** δ 29.28 (Note that 151.07 signal is an artifact).
**LC-MS (ESI-Q)**: 584.17 $[M+H]^+$
**HRMS (ESI-TOF):** m/z calculated for $C_{24}H_{46}N_3O_9PS$ $[M+H]^+$: 584.2764, found: 584.2773.

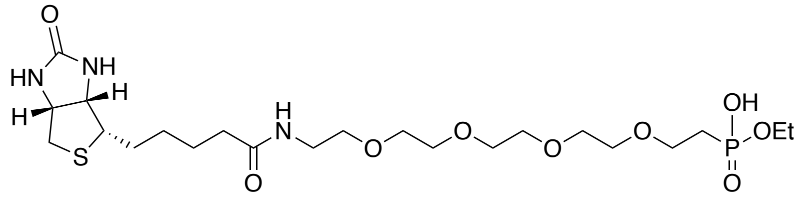

**S5**

**ethyl hydrogen (16-oxo-20-((3a*S*,4*S*,6a*R*)-2-oxohexahydro-1*H*-thieno[3,4-*d*]imidazol-4-yl)-3,6,9,12-tetraoxa-15-azaicosyl)phosphonate (S5)** (2)
**S4** (50 mg, 0.085 mmol) was dissolved in DMF (3 mL) and charged into a flask equipped with stir bar. Lithium azide (158 mg of solution, 0.645 mmol; 20 wt% solution in water) was added to the dissolved **S4** and the mixture stirred at 100°C overnight. Solvent was reduced *in vacuo*, then purified by silica gel flash chromatography using a dichloromethane/methanol (80:20 → 50:50) gradient to give **S5** (37 mg, 0.067 mmol, 78%) as a colorless oil. Analytically pure sample was obtained by preparative HPLC (19x100 mm C18) using a $H_2O$/MeCN (0.1% TFA) gradient (100:0 → 20:80).
**$R_f$**: 0.38 (DCM/MeOH 50:50, visualized with $I_2$/silica or CAM, streaks)
**$^1$H NMR (600 MHz, $CDCl_3$):** δ 7.72 (br s, 1H), 7.49 (br s, 1H), 4.51 (dd, *J* = 8.0, 4.6 Hz, 1H), 4.36 (dd, *J* = 7.9, 4.6 Hz, 1H), 4.03 (m, 2H), 3.78 (m, 2H), 3.61 (m, 16H), 3.52 (dq, *J* = 15.0, 5.0 Hz, 1H), 3.39 (dq, *J* = 14.7, 4.9 Hz, 1H), 3.16 (td, *J* = 7.4, 4.5 Hz, 1H), 2.91 (dd, *J* = 12.9, 4.9 Hz, 1H), 2.72 (d, *J* = 12.8 Hz, 1H), 2.29 (td, *J* = 7.3, 3.7 Hz, 2H), 2.07 (m, 2H), 1.73 (m, 4H), 1.47 (m, 2H), 1.29 (t, *J* = 7.1 Hz, 3H).
**$^{13}$CNMR (151 MHz, $CDCl_3$):** δ 173.91, 164.82, 70.48, 70.44, 70.37, 70.35, 70.29, 69.97, 69.91, 66.13 (d, $J_{C\text{-}P}$ = 1.6 Hz), 62.41, 60.75 (d, $J_{C\text{-}P}$ = 6.1 Hz), 60.30, 55.95, 40.75, 39.26, 35.63, 28.21, 27.98, 27.60 (d, $J_{C\text{-}P}$ = 137.6 Hz), 25.65, 16.70 (d, $J_{C\text{-}P}$ = 6.6 Hz).
**$^{31}$PNMR (162 MHz, $CDCl_3$):** δ 27.40 (Note that 151.07 signal is an artifact).
**LC-MS (ESI-Q)**: 556.12 $[M+H]^+$, 578.10 $[M+Na]^+$
**HRMS (ESI-TOF):** m/z calculated for $C_{22}H_{46}N_3O_9PS$ $[M+H]^+$: 556.2452, found: 556.2451.

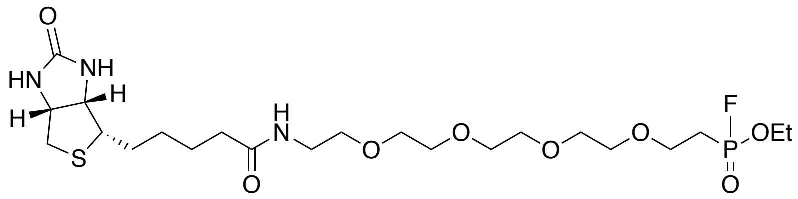

**S6** (FP-Probe)

**ethyl (16-oxo-20-((3a*S*,4*S*,6a*R*)-2-oxohexahydro-1*H*-thieno[3,4-*d*]imidazol-4-yl)-3,6,9,12-tetraoxa-15-azaicosyl)phosphonofluoridate (S6; FP-Probe)** (3, 4)

**S5** (15 mg, 0.027 mmol) was dissolved in anhydrous dichloromethane (3 mL) and charged into a flame-dried scintillation vial equipped with stir bar. The solution was cooled to -42ºC using an acetonitrile/dry ice bath before diethylaminosulfur trifluoride (13 mg/10.7 $\mu$L, 0.081 mmol) was added via auto pipette. The reaction stirred at -42ºC for 30 minutes before warming to room temperature. The reaction was quenched with 500 $\mu$L of water, then quickly treated with excess anhydrous magnesium sulfate to dry. The resultant slurry was filtered over Celite and dried *in vacuo* to give a crude waxy solid. The solid was dissolved in dichloromethane, then purified by flash chromatography using a small silica gel plug and a dichloromethane/methanol (80:20) solvent system to yield **S6** (3.4 mg, 0.006 mmol, 22%) as a waxy white residue. **S6** was dried *in vacuo* thoroughly and immediately after purification to limit P–F bond hydrolysis from ambient moisture. Probe activity was verified by selective labeling and enrichment of cell-culture grade trypsin (Gibco Trypsin-EDTA with phenol red).

**$R_f$**: 0.47 (DCM/MeOH 80:20, visualized with CAM)

**$^{1}$H NMR (600 MHz, $CDCl_3$):** δ 6.87 (br s, 1H), 6.19 (br s, 1H), 5.27 (br s, 1H), 4.50 (m, 1H), 4.32 (m, 1H), 4.27 (m, 2H), 3.79 (dt, *J* = 15.6, 7.1 Hz, 2H), 3.64 (m, 12H), 3.57 (t, *J* = 5.0 Hz, 2H), 3.44 (p, *J* = 5.3 Hz, 2H), 3.15 (td, *J* = 7.4, 4.4 Hz, 1H), 2.91 (dd, *J* = 12.8, 5.0 Hz, 1H), 2.74 (d, *J* = 12.7 Hz, 1H), 2.29 (m, 2H), 2.24 (m, 2H), 1.69 (m, 4H), 1.45 (p, *J* = 7.4 Hz, 2H), 1.38 (t, *J* = 7.1, 3H).

**$^{13}$CNMR (151 MHz, $CDCl_3$):** δ 173.50, 163.78, 77.16, 70.61, 70.50, 70.49, 70.48, 70.46, 70.20, 70.13, 64.31 (d, $J_{C\text{-}P}$ = 2.8 Hz), 63.70 (d, $J_{C\text{-}P}$ = 7.2 Hz), 61.89, 60.27, 55.60, 40.68, 39.22, 35.94, 28.23, 25.88 (dd, $J_{C\text{-}P}$ = 144.0 Hz, $J_{C\text{-}F}$ = 22.9 Hz), 25.68, 16.47 (d, $J_{C\text{-}P}$ = 5.5 Hz).

**LC-MS (ESI-Q)**: 558.63 $[M+H]^+$

**HRMS (ESI-TOF):** m/z calculated for $C_{22}H_{41}FN_3O_8PS$ $[M+H]^+$: 580.2228, found: 580.2238.

### ***Sample Preparation and Data Collection***

<u>Microbiome extraction</u>

Stool samples stored at -20ºC were thawed to room temperature, diluted with PBS (pH 7.4), and vortexed thoroughly to yield heterogeneous slurries. Slurries were centrifuged at 100 xg for 1 min before flocculent supernatants were extracted and filtered over disposable 70 µm nylon mesh cell strainers to remove large recalcitrant masses. Microbial cells were pelleted by centrifugation at 8000 xg for 5 min and separated from supernatants. Pellets were rinsed twice with, then resuspended in, PBS (pH 7.4). Suspensions were transferred into 1.5 mL Eppendorf tubes to a final density of 100 mg wet microbial pellet per 500 µL of suspension. Microbiome aliquots were stored at -20 ºC.

<u>Microbial DNA extraction</u>

Microbial DNA was extracted from thawed fecal microbe aliquots (see "Microbial extraction" section) using the Zymo fecal/soil extraction kit according to the provided protocol. Extracted DNA was quantified using a NanoDrop spectrophotometer then stored at -20 ºC.

<u>16S library preparation and sequencing</u>

Extracted microbiome DNA was sent to the Scripps Research genomics core for library preparation and sequencing. Libraries were prepared using the NEXTflex$^{TM}$ 16S V4 Amplicon-Seq Kit according to the manufacturer's instructions. Paired-end sequencing (2 x 300 bp) covering the ends of whole PCR product was carried out on the MiSeq platform (Illumina Inc, San Diego, CA, USA) aiming for 100,000 reads per sample. For access to raw 16S amplicon sequencing data sets, see Zenodo data repository doi:10.5281/zenodo.4265371.

<u>Microbial protein extraction and proteomics sample preparation (general)</u>

Protein extraction was performed according to previously described procedure (5). Microbiome extracts (100 mg pellets in Eppendorf tubes, *see above*) stored at -20ºC were lyophilized. Lyophilized samples were then treated with 300 µL of freshly prepared triflic acid solution (1:9 anhydrous toluene/triflic acid) at -78ºC (dry ice/acetone bath). Samples were warmed to 4ºC and agitated by nutation for 60 min with venting every 20 min to release pressure. After incubation, samples were cooled to -78ºC, neutralized with 900 µL of cold water/methanol/pyridine (1:1:3) solution, then slowly warmed to room temperature. Upon complete neutralization, resultant suspensions were diluted into 10mL of methanol/water (1:1) solution and transferred into 3K MWCO Amicon Ultra-15 centrifugal concentrators. Suspensions were concentrated down to 1-2 mL, resuspended in $H_2O$ up to 15 mL, then concentrated down to 1-2 mL again. This was repeated for a minimum of 3 cycles before transferring suspensions into clean 15 mL conical tubes and lyophilizing. Lyophilized material was resuspended in 100-200 µL of $H_2O$ before protein concentrations were measured by BCA assay.

100 µg of protein per sample was suspended in 120 µL of denaturation buffer (5 M Urea, 100 mM Bicine, pH 8.0). Protein suspensions were reduced with TCEP (5 mM final concentration), vortexed, and incubated for 15 min at room temperature. Protein suspensions were then alkylated with chloroacetamide (25 mM final concentration) and incubated for 45 min at room temperature shielded from light. Resultant mixtures were diluted to a final volume of 500 µL with trypsin buffer (100 mM Tris•HCl, 1 mM calcium chloride, pH 8.0) then treated with 2 µg of sequencing grade trypsin. Mixtures were incubated with agitation overnight at 37ºC. After incubation, resultant solutions were treated with 25 µL of formic acid, then centrifuged at 12,000 xg for 5 min. 475 µL of peptide supernatant were extracted and divided into a 375 µL portion for long-term storage at -20ºC, and a 100 µL portion for desalting by Ziptip C18. Desalted peptides were dried by centrifugal vacuum concentration and sent for LC-MS/MS analysis.

<u>FP probe labeling, enrichment, and proteomics sample preparation</u>

Microbiome aliquots (100 mg pellets suspended in 500 µL volume, *see above*) were thawed and treated with SDS to a final concentration of 0.1% (w/v). Samples were then sonicated at 25% amplitude for 10 min at 4ºC using a cup horn sonicator. Sonicated suspensions were expanded to 1 mL with 500 µL of PBS pH 7.4, vortexed, then divided into two 500 µL portions. One portion was treated with DMSO (vehicle) and the other portion was treated with 10 mM FP-Probe (in anhydrous DMSO) to a final concentration of 10 µM. Samples were vortexed and incubated at room temperature for 30 minutes with continuous agitation by inversion. After incubation, samples were loaded into 10K MWCO Amicon Ultra-15 centrifugal concentrators, treated with 5 mL denaturation buffer (8 M Urea, 100 mM Tris, pH 8.5) and concentrated. Samples were then diluted with 15 mL of PBS pH 7.4, then concentrated to remove unreacted/hydrolyzed FP probe. After concentration to ~400 µL, samples were transferred to clean Eppendorf tubes, treated with PBS to a final volume of 500 µL and SDS to a final concentration of 2%. Samples were boiled for 1 minute, then centrifuged at 12,000 xg to pellet insolubles. Supernatants were extracted, expanded to 10 mL with PBS pH 7.4, then treated with 200 µL of washed high capacity streptavidin-agarose beads. Bead treated supernatants were vortexed, agitated by inversion at room temperature for 3 h, then centrifuged at 4000 xg for 10 min to pellet streptavidin-agarose beads. Supernatants were discarded and remaining beads were treated with 10 mL PBS pH 7.4 + 0.1% SDS, vortexed, and agitated by inversion for 15 min. This was repeated two more times to wash beads of non-specifically bound proteins. Beads were then washed three more cycles with PBS pH 7.4 to remove residual SDS.

Approximately 10% of beads were extracted, treated with Laemmli buffer and boiled. Supernatants were analyzed by Western blot and stained with IRDye streptavidin to visualize probe enriched proteins. The remaining beads were prepared for LC-MS/MS. Beads were resuspended in denaturation buffer (5M Urea, 100 mM Bicine, pH 8.0) to a volume of 120 µL. Suspended beads were reduced with TCEP (5 mM final concentration), vortexed, then incubated for 15 min at room temperature. Beads were then alkylated with chloroacetamide (25 mM final concentration), vortexed, then incubated for 45 min at room temperature. Bead suspensions were diluted to 500 µL with trypsin buffer (100 mM Tris•HCl, 1 mM calcium chloride, pH 8.0), then treated with 2 µg of sequencing grade trypsin, vortexed, and incubated with agitation overnight at 37ºC. After an overnight digestion, bead slurries were treated with 25 µL of formic acid, vortexed, then centrifuged at 12,000 xg for 5 min. Supernatants were extracted, dried by centrifugal vacuum concentration, and desalted by Ziptip C18 to yield peptides for LC-MS/MS.

*Proteomics data collection (LC-MS/MS)*

Dried Ziptip C18 desalted peptide mixtures (1 µg) were resuspended in 10 µL of $H_2O$ (0.1% formic acid) and eluted from an Acclaim PepMap™ RSLC nano Viper analytical column (75 µm ID x 15 cm, Thermo Scientific, San Jose, CA) using a binary solvent gradient (A: $H_2O$ + 0.1% formic acid; B: 80:20 MeCN/$H_2O$ + 0.1% formic acid) at a flowrate of 300 nL/min delivered by a nEasy-LC1000 nano liquid chromatography system (Thermo Fischer Scientific, San Jose, CA). The following gradient was used: 5-25% solvent B over 180 min, followed by 25-44% solvent B over 60 min, 44-80% solvent B over 0.1 min, a 5 min hold at 80% solvent B, a return to 5% solvent B over 0.1 min, and finally a 20 min hold at 100% solvent B. Ions were created at 1.8 kV using the Nanospray Flex™ ion source (Thermo Fischer Scientific, San Jose, CA). Data dependent scanning was performed by the Xcalibur (v. 4.0.27.10) software package using a survey scan at 120,000 resolution in the Orbitrap mass analyzer scanning between m/z 380-1400 at an AGC target of 1.0e5 and a maximum injection time of 50 ms, followed by HCD fragmentation at a normalized collision energy of 30% for the most intense ions at maximum speed (topN) and AGC setting of 1.0e4. Precursor ions were selected by monoisotopic precursor selection (MIPS) setting to peptide and MS/MS was performed in the Orbitrap on ions with charges +2 to +8 at a resolution of 30,000. Dynamic exclusion was set to exclude ions after two times within a 30 sec window for 20 sec at a 10 ppm low and high mass tolerance. All LC-MS/MS data have been deposited to the ProteomeXchange Consortium via the PRIDE partner repository with the project accession identifier PXD022433.

### ***Data Analysis***

*16S amplicon sequencing data analysis*

Compressed paired end fastq files were imported into QIIME2, demultiplexed, and visualized (6). Sequences were trimmed (f-6, r-7), truncated (f-289,r-220) and further processed using the Dada2 pipeline (7). Multiple sequence alignment was performed with MAFFT followed by masking (8, 9). An unrooted phylogenetic tree was constructed with FastTree followed by a rooted tree using the midpoint-root method (10). Diversity analysis was performed using QIIME2 core metrics selecting a sampling depth of 15,000. Alpha rarefaction was performed at a max depth of 15,000 with 25 steps. Next a classifier was trained using the SILVA 132 database (arb-silva.de; downloaded March 2020) (11) extracting the V3/V4 regions using #341 f-primer: 'CCTAYGGGRBGCASCAG' and the #806 r-primer: 'GGACTACNNGGGTATCTAAT' with a min-length of 300 and a max length of 600. The classifier was trained using a naïve Bayes approach with the QIIME2 feature-classifier plugin (12). After classifying features, a taxa barplot and corresponding table were generated in QIIME2 and used for further taxonomy plotting shown in supplementary figures above.

*Proteomics data processing and analysis*

**Peptide and protein identification***:* Thermo .raw files were converted to .ms2 files using RawConverter 1.1.0.18 (13) operating in data dependent mode and selecting for monoisotopic m/z. Tandem mass spectra (.ms2 files) were identified by database search method using the Integrated Proteomics Pipeline 6.5.4 (IP2, Integrated Proteomics Applications, Inc., http://www.integratedproteomics.com). Briefly, databases containing forward and reverse (decoy) (14, 15) peptide sequences were generated from *in silico* trypsin digestion of protein sequences derived from large comprehensive public repositories (ComPIL 2.0) (16, 17). Tandem mass spectra were matched to peptide sequences using the ProLuCID/SEQUEST (1.4) (18-20) software package. The validity of spectrum-peptide matches were assessed using the SEQUEST-defined parameters XCorr (cross-correlation score) and DeltaCN (normalized difference in cross-correlation scores) in the DTASelect2 (2.1.4) (21, 22) software package. Search settings were configured as follows: (1) 5 ppm precursor ion mass tolerance, (2) 10 ppm fragment ion mass tolerance, (3) 1% peptide false discovery rate, (4) 2 peptide per protein minimum, (5) 600-6000 Da precursor mass window, (6) 2 differential modifications per peptide maximum (methionine oxidation: M+15.994915 Da), (7) unlimited static modications per peptide (cysteine carbamidomethylation: C+57.02146 Da), and (8) the search space included half- and fully tryptic (cleavage C-terminal to K and R residues) peptide candidates with up to 2 missed cleavage events. For the full collection of proteomics database-searched outputs, see PRIDE repository project PXD022433.

**Peptide and protein quantification:** Peptides identified from all 18 unenriched (samples not treated with FP probe) fecal extract LC-MS/MS runs (searched against the ComPIL 2.0 database) were combined into an index file and used as input for quantification by FlashLFQ (23). This input file contained information for each database-identified MS2 spectrum spanning the following columns, "File Name", "Base Sequence", "Full Sequence", "Peptide Monoisotopic Mass", "Scan Retention Time", "Precursor Charge", and "Protein Accession". The "Protein Accession" column was left blank due to the abundance of peptides that map to multiple disparate proteins; FlashLFQ was only used to quantify peptide intensities. FlashLFQ used the index file and all 18 Thermo .raw files directly to determine MS1 peak intensities associated input peptides. The following settings were used, 10 ppm precursor tolerance, match between runs enabled, integrate peak areas enabled, 15 min maximum match between runs window, 2 isotopes required, and 5 ppm isotope mass tolerance.

For protein quantification, all detected protein sequences identified by initial database searching were clustered into groups based on sequence similarity. This was done by first combining all sequences into a single protein fasta file, then clustering was accomplished using CD-HIT (4.8.1) grouping at a 95% similarity cut-off (24-26). The following command line input was used, "cd-hit -i fastafile.fasta -o outputfile -c 0.95 -g 1 -d 0". Protein groups contained anywhere from one sequence to several dozen sequences. A total of 95,000 protein groups were generated from 576,625 protein sequences. Peptide intensities generated by FlashLFQ were then mapped to their parent protein sequences and subsequently mapped to CD-HIT generated protein groups. Peptides mapping to more than one protein group were discarded. Peptide intensities for each protein group were summed to generate total intensities for each protein group. These intensities were used for differential expression analysis. Note also that CD-HIT generates protein groups and designates one sequence within a given group with an asterisk ("*"); this protein sequence was designated the protein group/cluster representative. Finally, prior to differential expression analysis, a filtering step was applied to the data to exclude protein groups/clusters with too many missing/null entries. Healthy patient proteomics data were grouped into the healthy/control condition while ulcerative colitis patient proteomics data were grouped in the UC condition. For protein groups/clusters where one condition contained all null values, the other condition needed to contain >3 non-null values to be retained. For protein groups/clusters where both conditions contained non-null values, each condition needed >3 non-null values to be retained. Protein groups/clusters containing only null values for both conditions were discarded.

**Protein differential expression analysis**: Differential expression analysis of peptide ion intensity data were performed using the DEP package operating in the R statistical computing environment (27). Protein group/cluster summed intensities were automatically Log2 transformed in DEP then normalized by the “normalize_vsn” function (28). Missing data were imputed using a mixed approach. Protein groups/clusters where either healthy/control or UC condition contained only null entries were classified as 'missing not at random' (MNAR) and imputed with 0 values. All other groups were treated as 'missing at random' (MAR) and imputed using the Bayesian principal component analysis ('BPCA') imputation method (29). Note that for a given cluster, missing values for each condition were imputed separately by condition. Differential expression analysis was performed on filled-in protein group/cluster intensity tables using the “test_diff” function (30) and multiple testing correction was performed using the “add_rejections” function. Nominal *p* values and Benjamini-Hochberg multiple testing correction values were calculated in DEP (31). Storey *q*-values were calculated using the “qvalue” package in the R environment (32). Volcano plots, PCA plots, and heat maps were also generated in R (ggplot2: ggplot2.tidyverse.org; factoextra: rpkgs.datanovia.com/factoextra/index.html; enhancedvolcano: github.com/kevinblighe/EnhancedVolcano) (33). A significance level of $q < 0.1$ was chosen as a relevant cut-off revealing a combined 176 differentially enriched protein groups/clusters from healthy and ulcerative colitis patient fecal extracts. For a complete list of tested protein groups/clusters and their associated differential expression analysis outputs, see **SI_C**.

**STRING network analysis:** For STRING analysis, host protein groups/clusters identified as differentially enriched were extracted and submitted to the STRING web server (v.11; string-db.org) (34). No host proteins were identified as differentially enriched in healthy patient fecal extracts so analysis was only conducted on host proteins enriched in ulcerative colitis patient fecal extracts. With “homo sapiens” selected under organism type, the following proteins (representing protein groups/clusters) were submitted for analysis: NEP_HUMAN, ATPA_HUMAN, B3AT_HUMAN, EVI2B_HUMAN, M2OM_HUMAN, PHB2_HUMAN, IDHC_HUMAN, HBB_HUMAN, ANXA3_HUMAN, S10AC_HUMAN, IGG1_HUMAN, PRTN3_HUMAN, TRFL_HUMAN, ECP_HUMAN, CAP7_HUMAN, BPI_HUMAN, LV151_HUMAN, S10A9_HUMAN, SAMP_HUMAN, KVD11_HUMAN, SORCN_HUMAN, ANX11_HUMAN, KV320_HUMAN, IC1_HUMAN, ERP29_HUMAN, AMYP_HUMAN, ABCG2_HUMAN, EF2_HUMAN, VDAC1_HUMAN. Minimum required interaction scores were varied to identify high and medium confidence edges. Network graphs were generated by the web server and used without further modification.

**Proteomics-based taxonomic analysis**: Peptides identified from all 18 unenriched (samples not treated with FP probe) fecal extract LC-MS/MS runs (searched against the ComPIL 2.0 database) were combined into an index file. Modified peptides (oxidized methionine and carbamidomethylated cysteine) were converted to their non-modified forms. All duplicate peptides were collapsed such that the final peptide list contained no redundant entries. The resultant peptide list was submitted for Unipept metaproteomics analysis using the online web application (unipept.ugent.be) (35-37). Isoleucine and leucine were not equated and “filter duplicate peptides” was enabled. Taxonomy assignments for each peptide were outputted in a .csv file and this file was used to generate taxonomy tables. FlashLFQ generated peptide intensities (*see above*) were mapped to Unipept peptides and subsequently to organisms at each of seven taxonomic levels (kingdom, phylum, class, order, family, genus, and species). For each organism at a given taxonomic level, corresponding constituent peptide intensities were summed. Peptide intensities that could not be mapped were binned into the “Unassigned/unknown” category. At each taxonomic level, organism intensities were divided by total intensity at that taxonomic level to generate relative abundance values. Relative abundance values were used to create stacked bar plots depicting taxonomy. To simplify taxonomy stacked bar plots, low abundance organisms were binned into the “Other” category. Full relative abundance values can be found in **SI_A**.

**GO relative abundance analysis:** The corresponding sequences of protein group/cluster representatives generated by CD-HIT were concatenated into a protein fasta file then submitted for InterProScan (5.40-77.0) analysis using the following command line input, “./interproscan.sh -i inputfile.fasta -f tsv -dp -goterms” (38-41). GO terms contained in output .tsv files were mapped to their corresponding protein sequences and corresponding protein groups/clusters. Protein group/clusters were functionally annotated using only the InterProScan output for their respective representative. GO terms were mapped to their name spaces using the comprehensive go.obo listing available at geneontology.org (42, 43).

For unweighted GO relative abundance assembly, GO terms were organized such that all representative protein sequences contributed at least 1 count to each of the three “Biological Process,” “Molecular Function,” and “Cellular Component” GO name spaces. If sequences could not be annotated, 1 count of “None” was contributed to each of the three name spaces. For sequences annotated multiple times, each annotation contributed 1 count to the GO term’s respective name space. GO term relative abundances were calculated by dividing term counts by respective name space total counts. Relative abundance values were used to generate stacked bar plots. To simplify GO relative abundance stacked bar plots, low abundance terms were binned into the “Other” category. Full relative abundance values can be found in the **SI_D**.

For weighted GO relative abundance assembly, protein group/cluster average peptide intensities were mapped to each GO term identified for that respective group/cluster. Average peptide intensities were used instead of summed peptide intensities to reduce sequence length bias (smaller proteins can be as functionally impactful as larger proteins). GO terms were organized such that all representative protein sequences contributed their average peptide intensities to each of three "Biological Process," "Molecular Function," and "Cellular Component" GO name spaces. If representative sequences could not be annotated, average peptide intensities were added to the "None" category for each GO name space. For sequences annotated multiple times, the average peptide intensity for a group/cluster was added to each GO annotation's total, undivided. GO term relative abundances were calculated by dividing GO term intensities by respective name space total intensities. Relative abundance values were used to generate stacked bar plots. To simplify GO relative abundance stacked bar plots, low abundance terms were binned into the "Other" category. Full relative abundance values can be found in the **SI_D**.

**GO enrichment analysis**: Enrichment analysis was performed using the GOstats (2.52.0) package in the R statistical computing environment (44). GO terms derived from significantly enriched ($q < 0.1$) protein groups as determined by differential expression analysis (see above) were selected for testing (subset proteins). GO terms mapped to all CD-HIT protein groups/clusters were selected as the 'universe' GO term set. Enrichment analysis was performed for each GO name space and independently for GO term sets enriched in the healthy/control condition versus the UC condition. The "pvalueCutoff" for significant enrichment was set at 0.01 and the test directions were set to "over". Output tables containing hypergeometric testing p values, odds ratios, expected counts, counts, size, and elaborated GO terms were used to construct GO enrichment bubble plots.

**FP Probe enriched peptide and protein identification:** A two-step database search strategy was used to identify FP probe enriched proteins (45, 46). .raw files from LC-MS/MS runs of FP probe enriched patient fecal extracts were searched against the ComPIL 2.0 database as previously described (see "Peptide and protein identification" section above) with the exception that a 100% peptide false discovery rate was enabled. Protein sequences identified from all 3 FP probe-enriched patient samples were concatenated into a single protein fasta file along with all sequences from the human proteome (Homo sapiens canonical and isoforms, Uniprot.org, downloaded 22 March 2020). Reverse sequences were then generated and concatenated to this fasta file before in silico trypsin digestion to generate a peptide database for a second proteomics database search. A second database search was conducted using the same settings as the first search except that peptide false discovery rate was set to 1%. Results for this second search were used to identify FP probe enriched proteins. Host proteins and probable host proteins were aggregated. Non-host proteins were listed separately.

***De novo* peptide sequencing and estimation of unidentified peptide space**: Unenriched (not treated with FP probe) microbiome LC-MS/MS .raw files were converted to .ms2 files using RawConverter 1.1.0.18 (13) operating in data dependent mode and selecting for monoisotopic m/z. Output .ms2 files were subject to *de novo* peptide sequencing using Novor (1.06.0634) (47). The following settings were used, (1) enzyme = Trypsin, (2) fragmentation = HCD, (3) massAnalyzer = FT, (4) fragmentIonErrorTol = 0.03Da, (5) precursorErrorTol = 10ppm, (6) variableModiciations = Oxidation of M (0; M+15.9949); Deamidation of N (1; N+0.9840); Deamidation of Q (2; Q+0.9840), (7) fixedModifications = Carbamidomethylation of C (3; C+57.0215), and (8) forbiddenResidues = I, U. Output .csv files contained *de novo* evaluations for all MS2 fragmentation spectra along with scores characterizing the quality of *de novo* evaluations. This output was mapped to ComPIL 2.0 database searching output (see "Peptide and protein identification" section above) such that each MS2 fragmentation spectrum possessed a *de novo*-assigned peptide and when applicable, a database-assigned peptide. For each MS2 fragmentation spectrum with a database-assignment, a Needleman-Wunsch comparison was performed using the "global_pairwise_align_protein" function provided by the Scikit-bio package for Python 3+ (scikit-bio.org) (48). Needleman-Wunsch scores were scaled to 100 by dividing *de novo*-database comparison scores by database-database (self-self; maximum attainable value for a sequence) comparison scores. These scaled scores were used to determine the quality of *de novo* peptide assignments. Note that prior to performing comparisons, peptide modifications were removed (i.e. oxidation, carbamidomethylation, deamidation) and leucine and isoleucine residues were rendered equivalent. Novor evaluation scores and *de novo*-database comparison scores were used to generate plots and data tables to estimate the number of database-elusive MS2 fragmentation spectra that likely represent true peptides. Because Novor evaluation scores correlate strongly with *de novo*-database comparison scores, a reasonably high Novor evaluation score of >75 was chosen as the lower limit for estimating the number of high confidence database-elusive MS2 fragmentation spectra that likely map to peptides. First, the number of MS2 fragmentation spectra with Novor evaluation scores >75 and *de novo*-database comparison scores >75% were divided by the number of MS2 fragmentation spectra with Novor evaluation scores >75 and database-assignments to generate a proportion value. The number of MS2 fragmentation spectra with Novor evaluation scores >75 and no database-assignments was multiplied by this proportion value to determine the number of database-elusive MS2 fragmentation spectra that are likely peptides. This estimation was applied to each patient sample LC-MS/MS run separately. The number of peptide-likely MS2 fragmentation spectra with Novor evaluation scores >75 was divided by the total number MS2 fragmentation spectra collected for a whole run to determine the percent of high confidence database-elusive MS2 fragmentation spectra that are likely peptides for a whole LC-MS/MS run. This percent value was determined for each LC-MS/MS run and the global mean was determined to be 9%.

### NMR Spectra

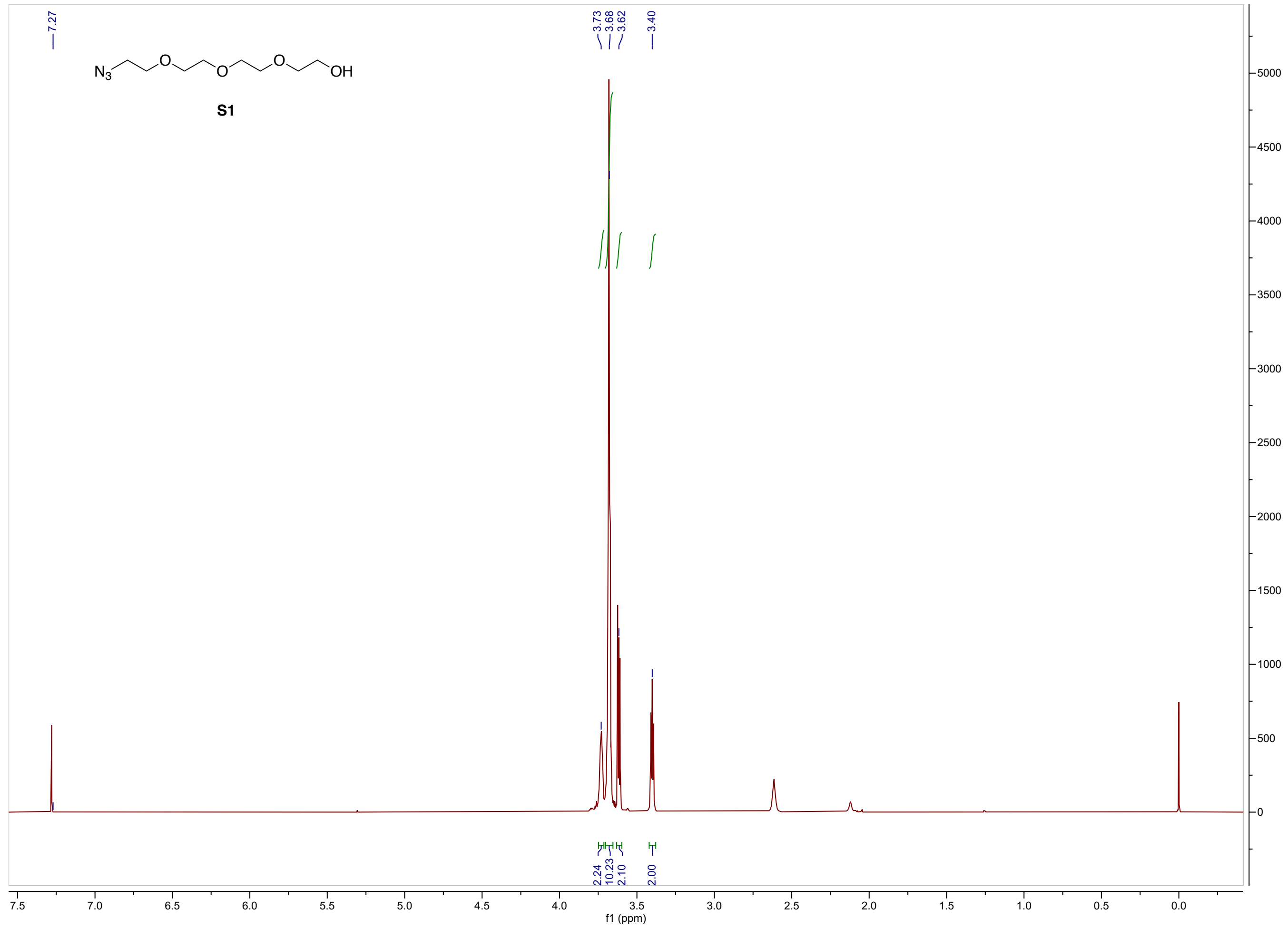


**S1** $^1$H

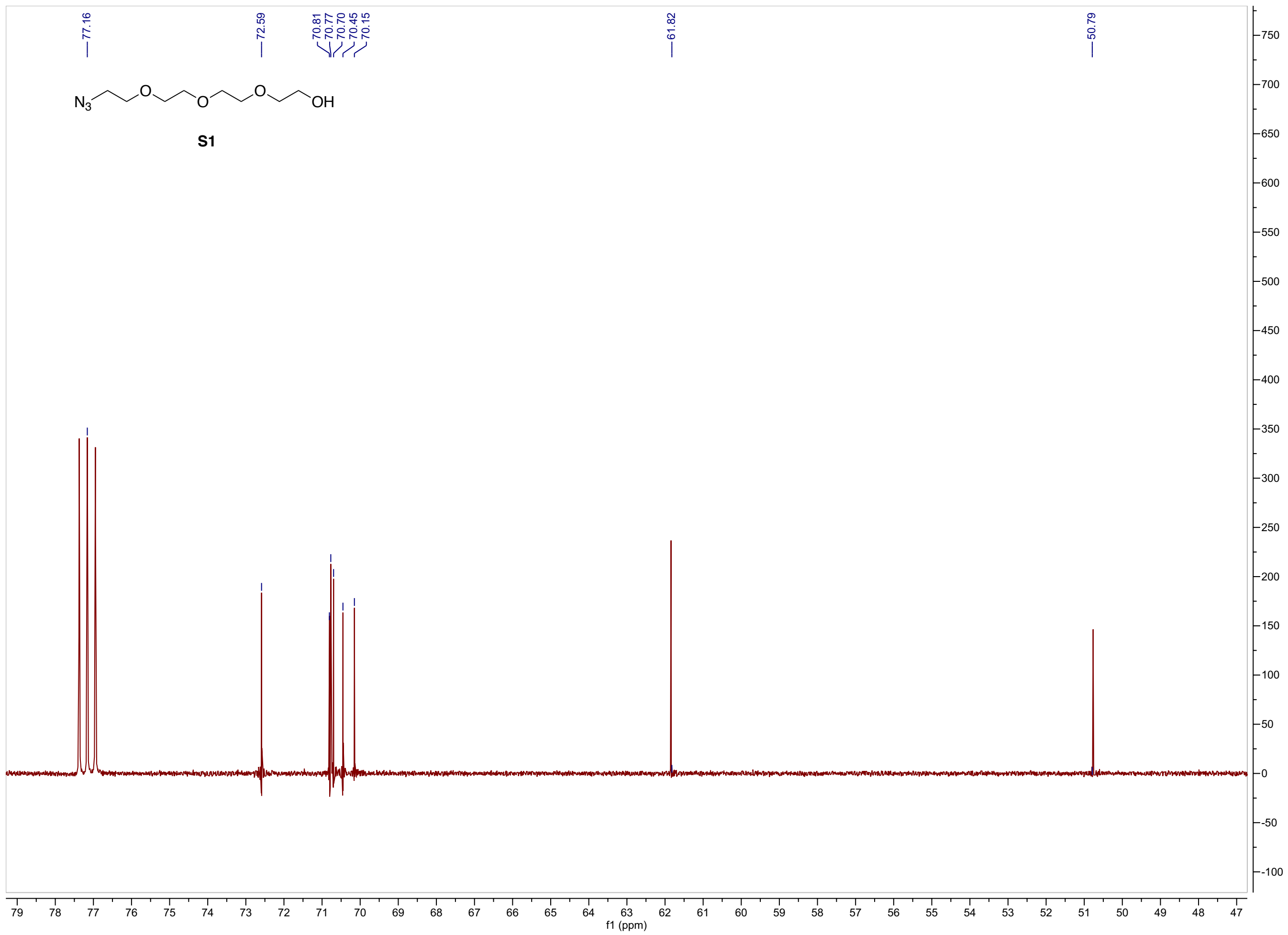
77.16
72.59
70.81
70.77
70.70
70.45
70.15
61.82
50.79
N3
O
O
O
OH
S1
f1 (ppm)

**S1** $^{13}C$

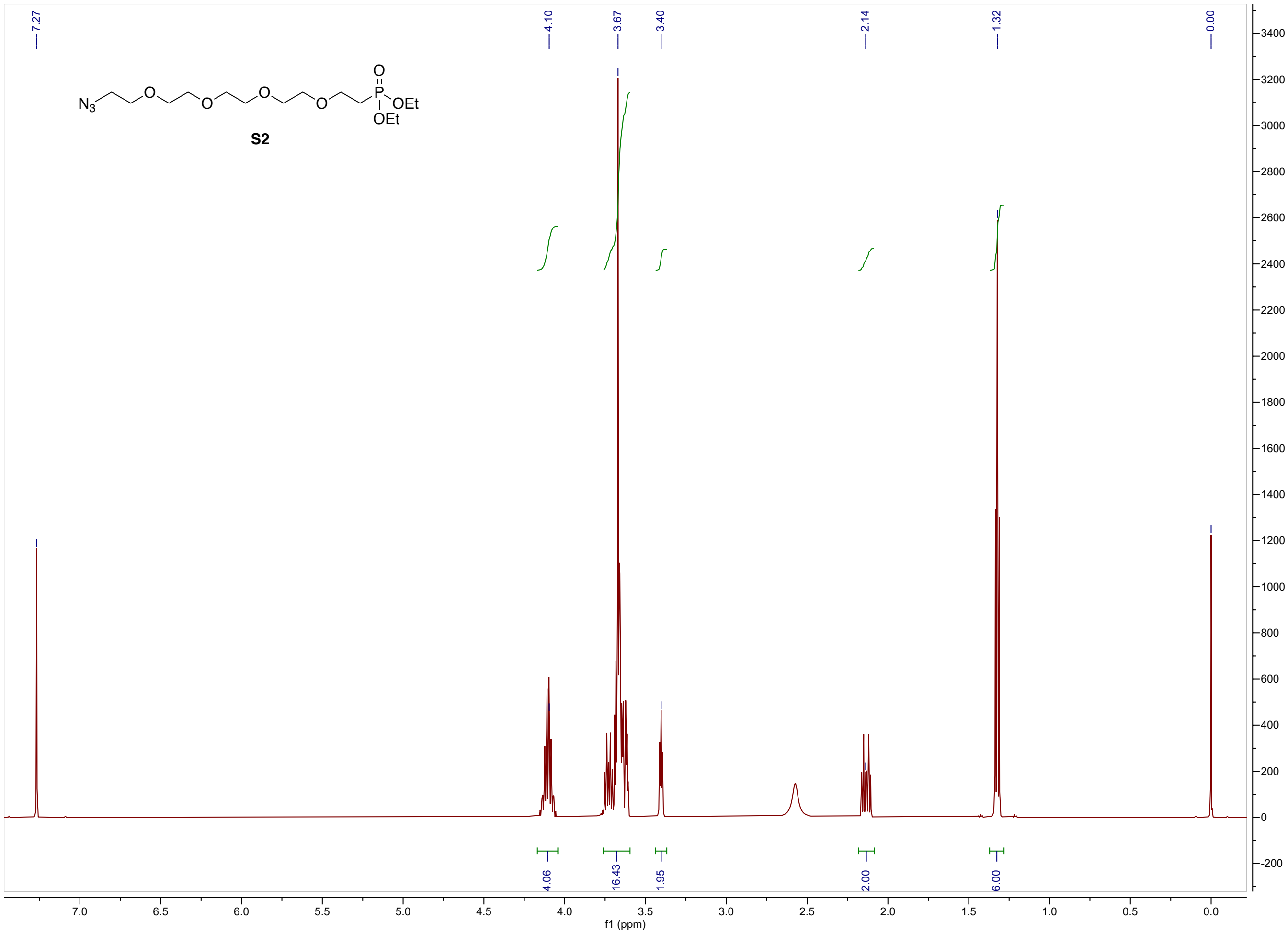


**S2** $^{1}$H

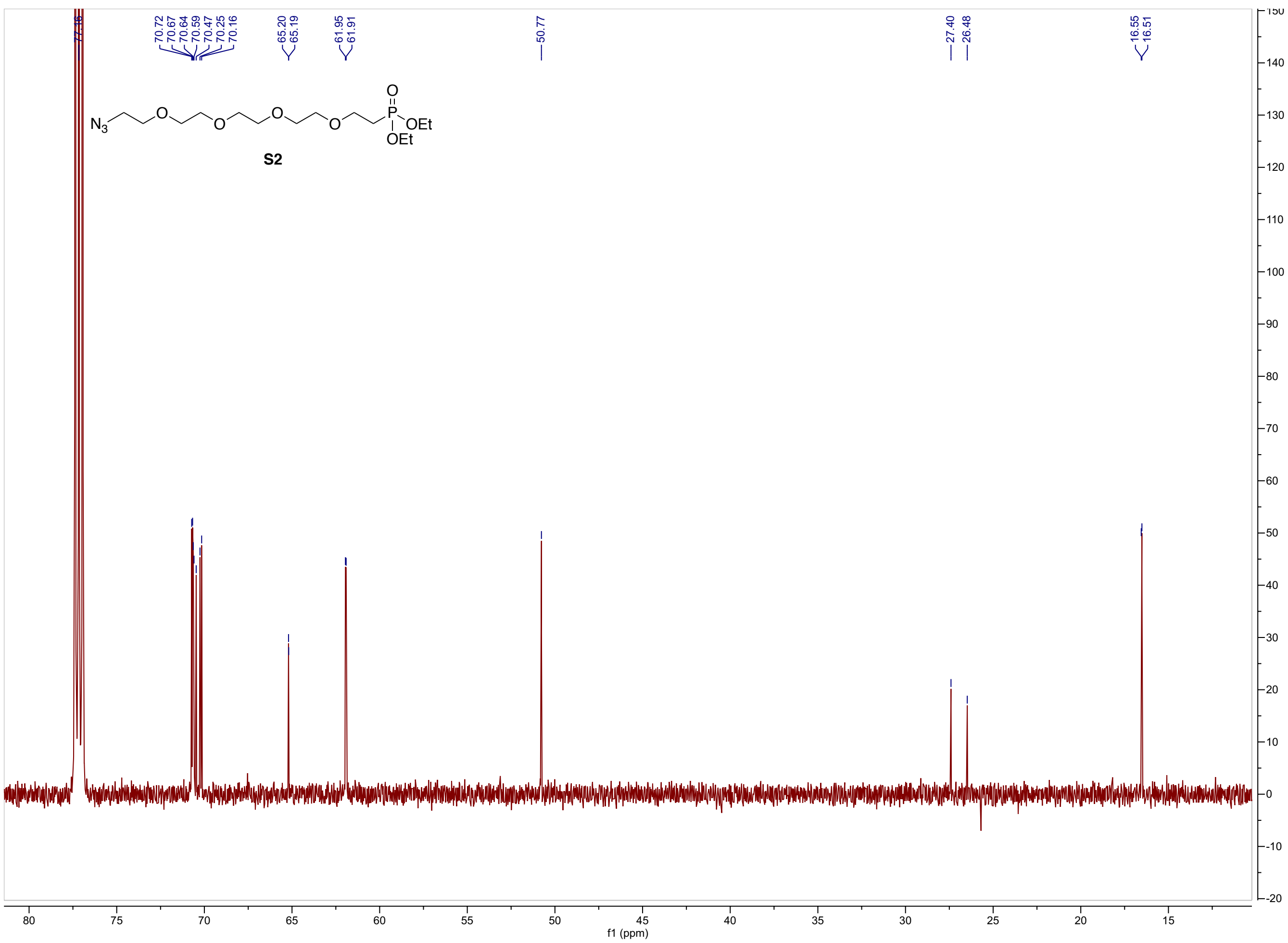
77.16
70.72
70.67
70.64
70.59
70.47
70.25
70.16
65.20
65.19
61.95
61.91
50.77
27.40
26.48
16.55
16.51
S2
f1 (ppm)

**S2** $^{13}$C

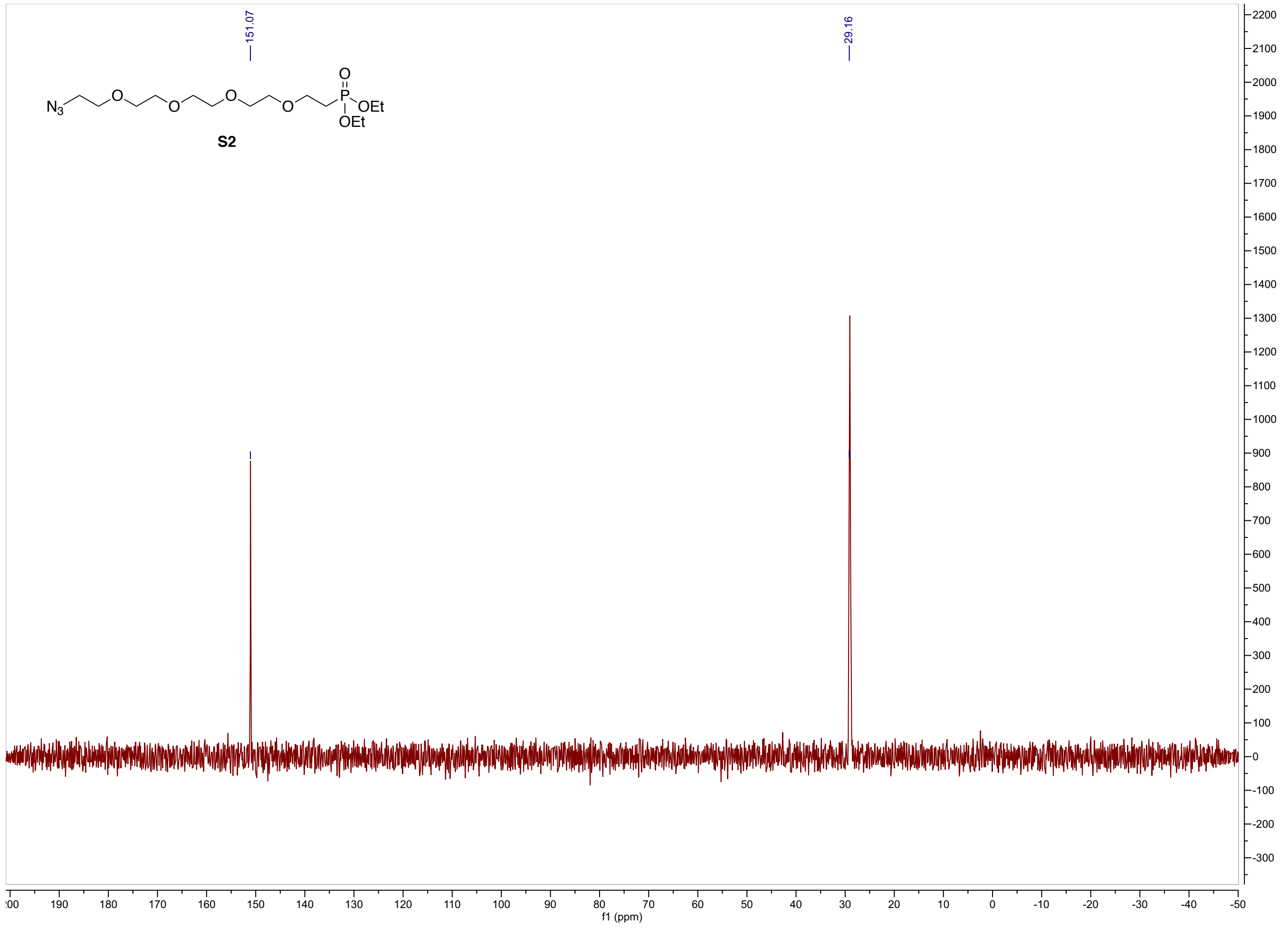


**S2** $^{31}$P

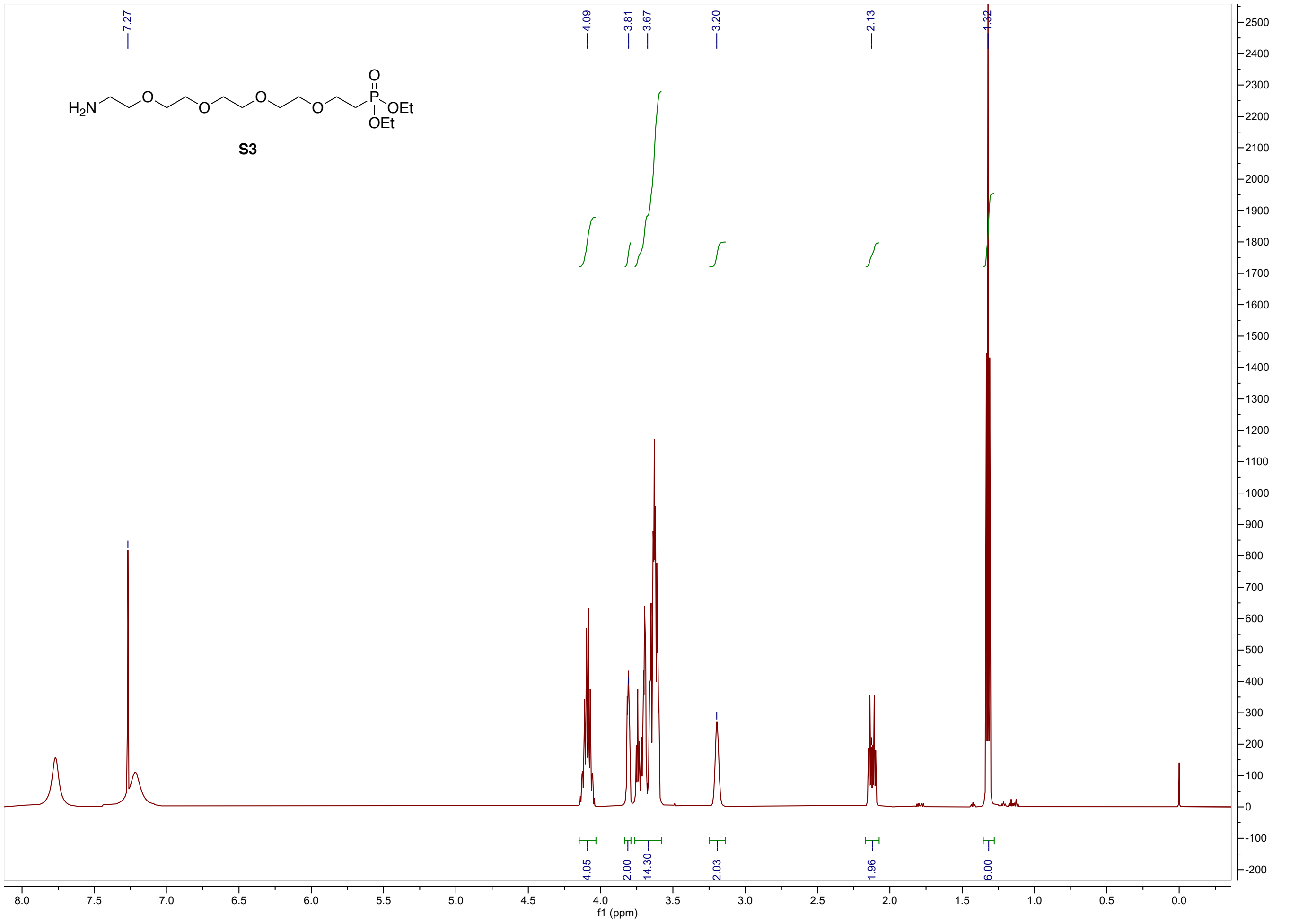

7.27
4.09
3.81
3.67
3.20
2.13
1.32
H2N
O
O
O
O
P
OEt
OEt
S3
4.05
2.00
14.30
2.03
1.96
6.00
f1 (ppm)


**S3** $^{1}$H

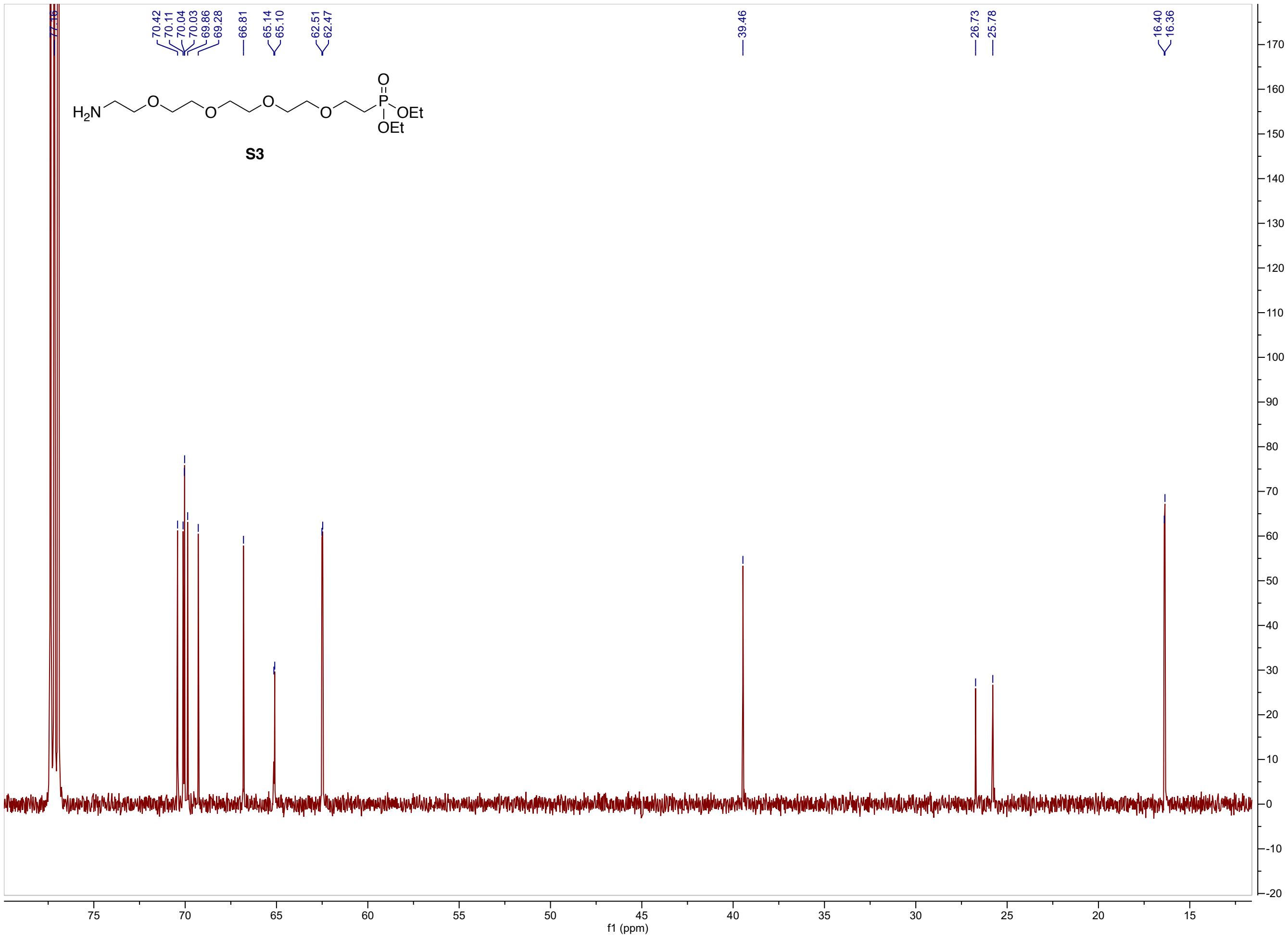

77.16
70.42
70.11
70.04
70.03
69.86
69.28
66.81
65.14
65.10
62.51
62.47
39.46
26.73
25.78
16.40
16.36
H2N
O
P
OEt
OEt
S3
f1 (ppm)


**S3** $^{13}C$

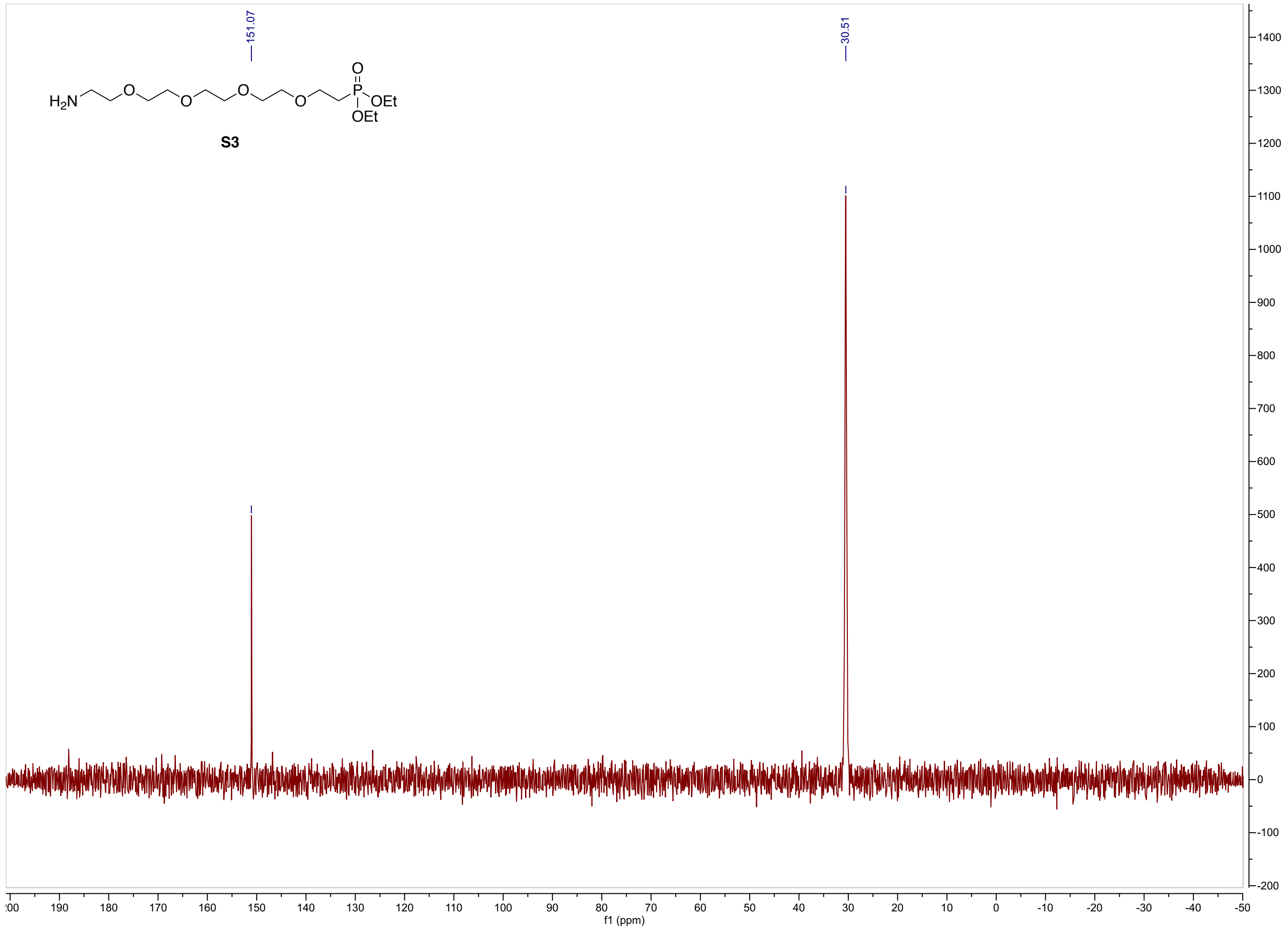


**S3** $^{31}$P

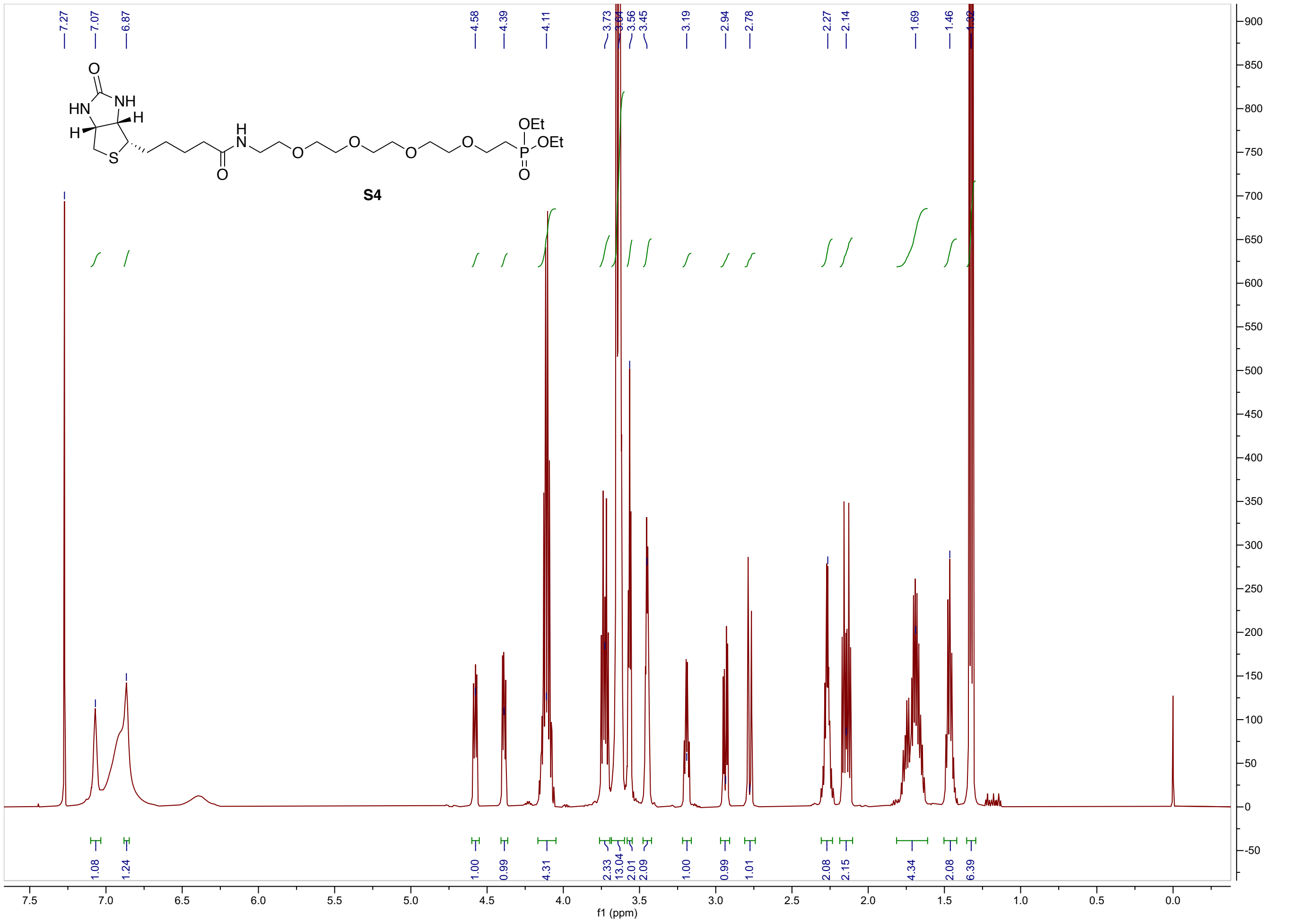
7.27
7.07
6.87
4.58
4.39
4.11
3.73
3.56
3.45
3.19
2.94
2.78
2.27
2.14
1.69
1.46
S4
1.08
1.24
1.00
0.99
4.31
2.33
13.04
2.01
2.09
1.00
0.99
1.01
2.08
2.15
4.34
2.08
6.39
f1 (ppm)

**S4** $^{1}$H

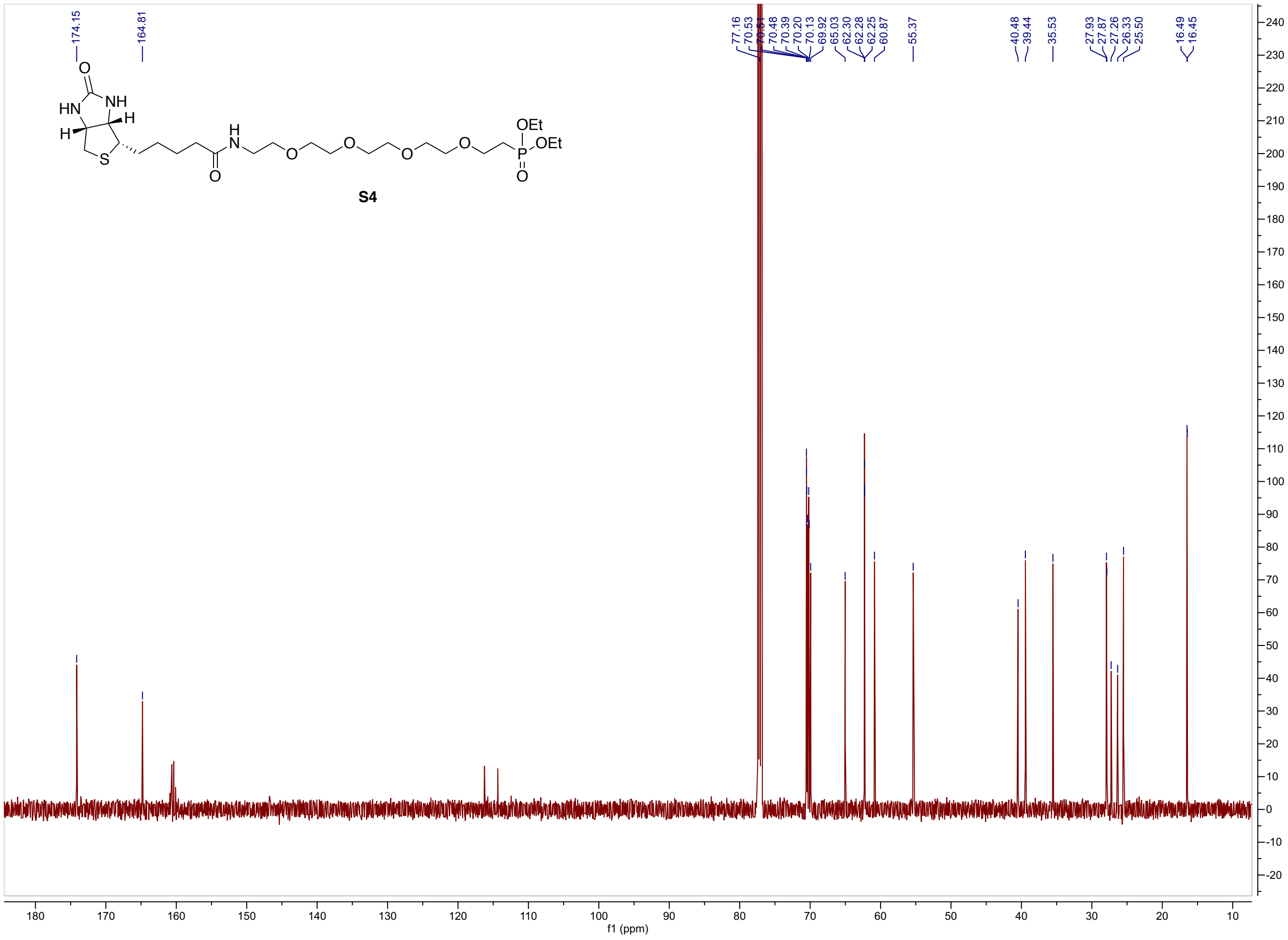
174.15
164.81
77.16
70.53
70.48
70.39
70.20
70.13
69.92
65.03
62.30
62.28
62.25
60.87
55.37
40.48
39.44
35.53
27.93
27.87
27.26
26.33
25.50
16.49
16.45
S4
f1 (ppm)

**S4** $^{13}$C

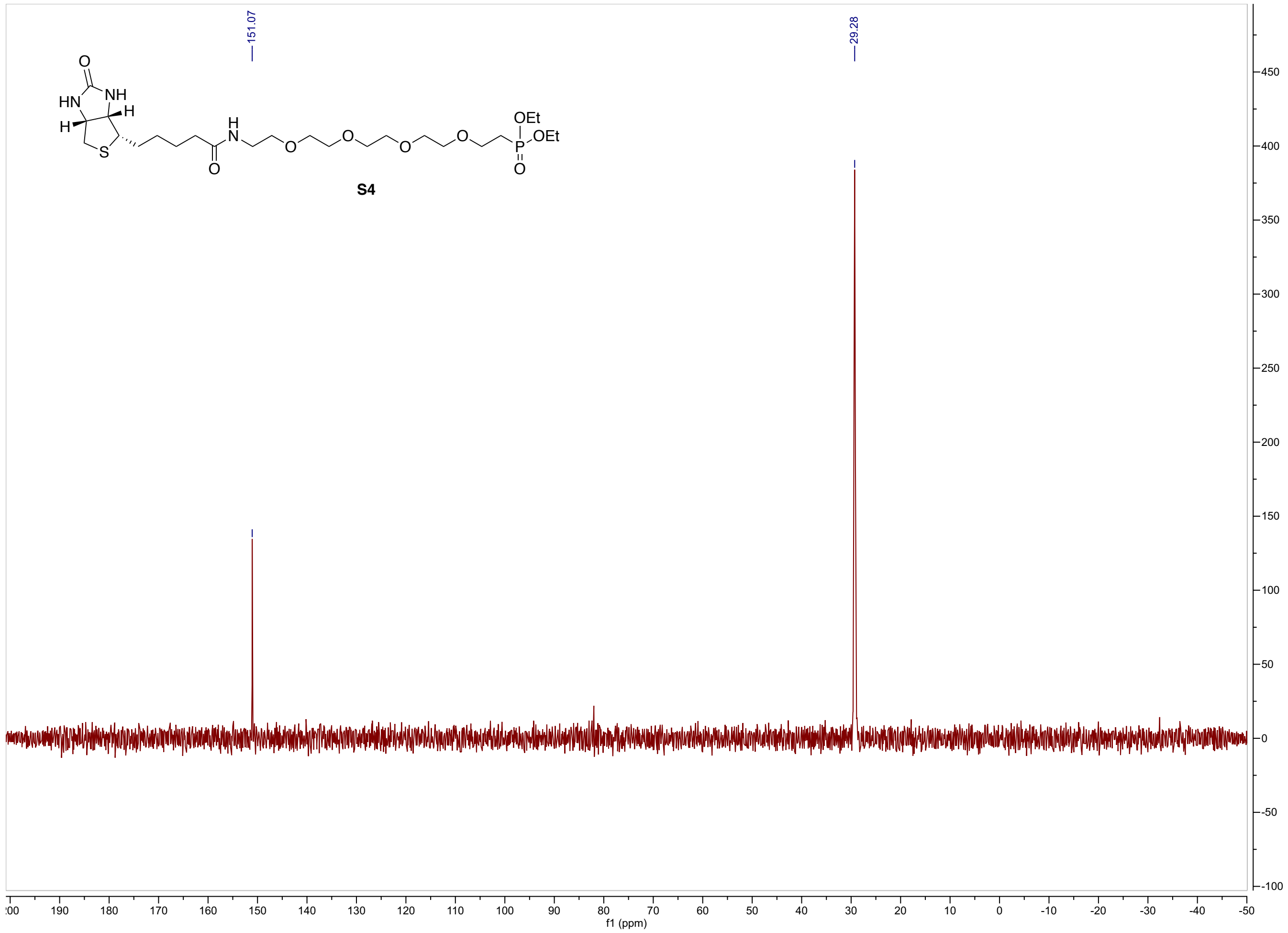
151.07
29.28
O
NH
HN
H
H
H
N
S
O
O
O
O
O
OEt
OEt
P
O
S4
450
400
350
300
250
200
150
100
50
0
-50
-100
f1 (ppm)

**S4** $^{31}P$

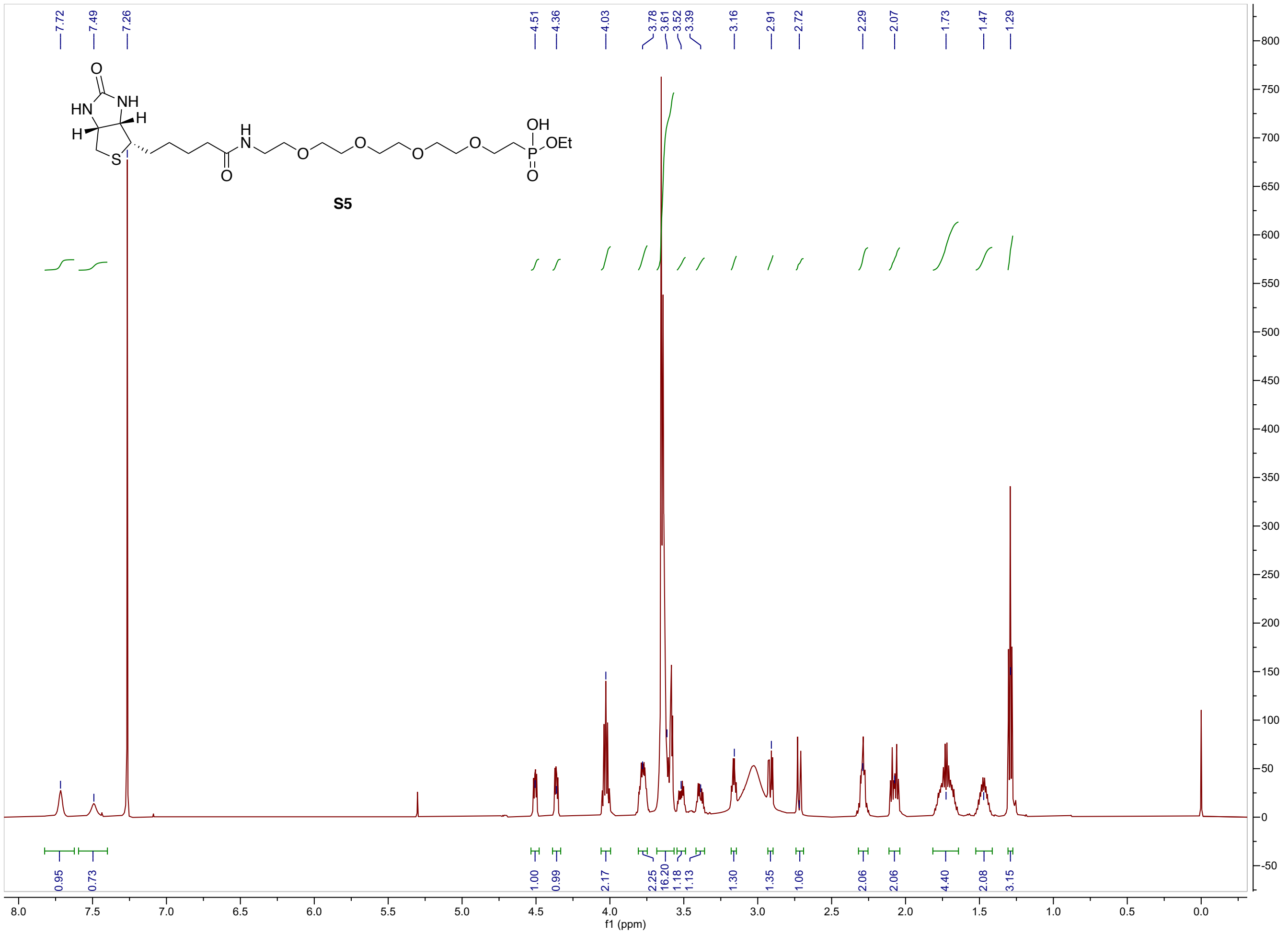

7.72
7.49
7.26
4.51
4.36
4.03
3.78
3.61
3.52
3.39
3.16
2.91
2.72
2.29
2.07
1.73
1.47
1.29
S5
0.95
0.73
1.00
0.99
2.17
2.25
16.20
1.18
1.13
1.30
1.35
1.06
2.06
2.06
4.40
2.08
3.15
f1 (ppm)


**S5** $^{1}H$

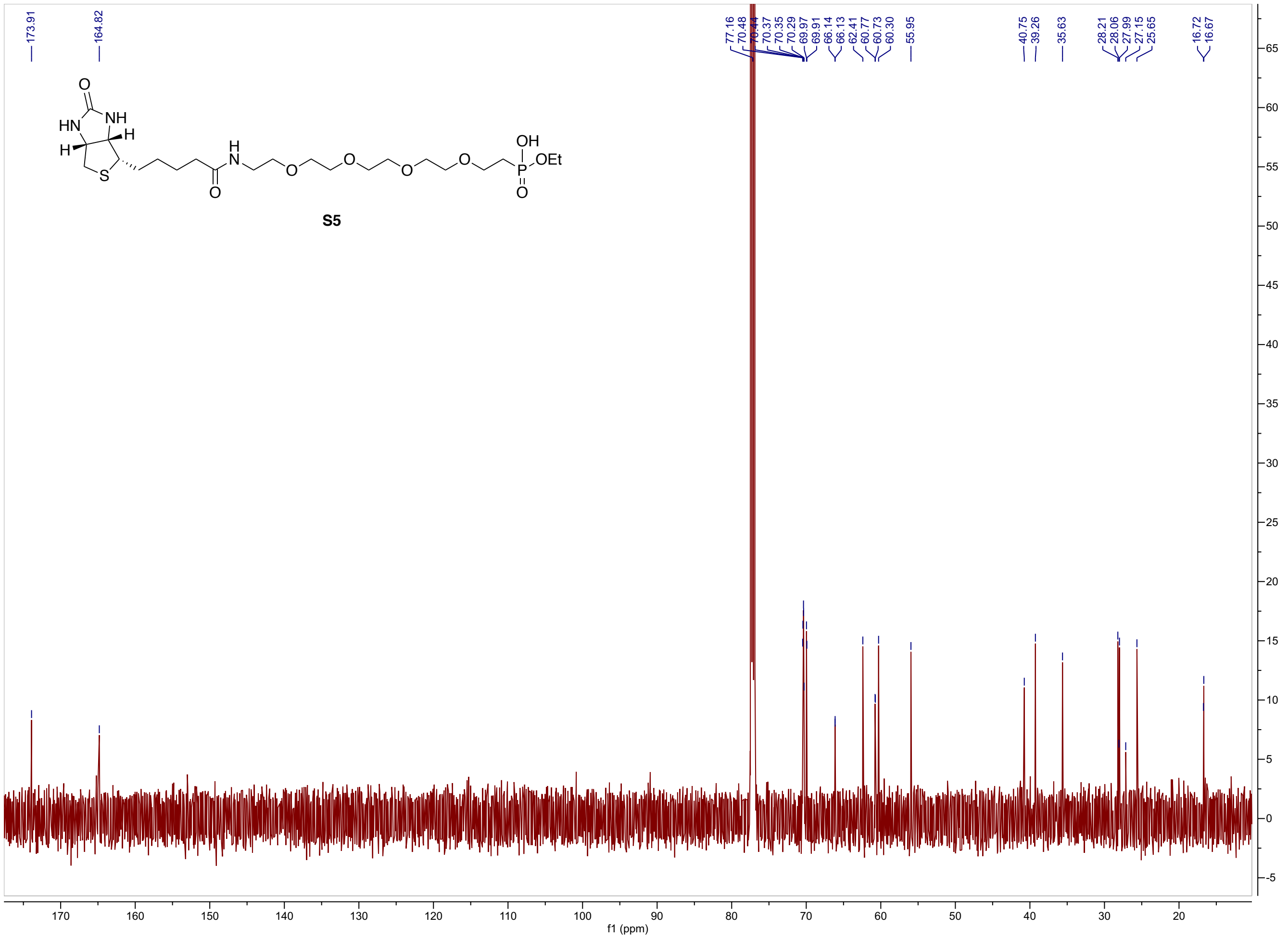
173.91
164.82
77.16
70.48
70.44
70.37
70.35
70.29
69.97
69.91
66.14
66.13
62.41
60.77
60.73
60.30
55.95
40.75
39.26
35.63
28.21
28.06
27.99
27.15
25.65
16.72
16.67
S5
f1 (ppm)

**S5** $^{13}C$

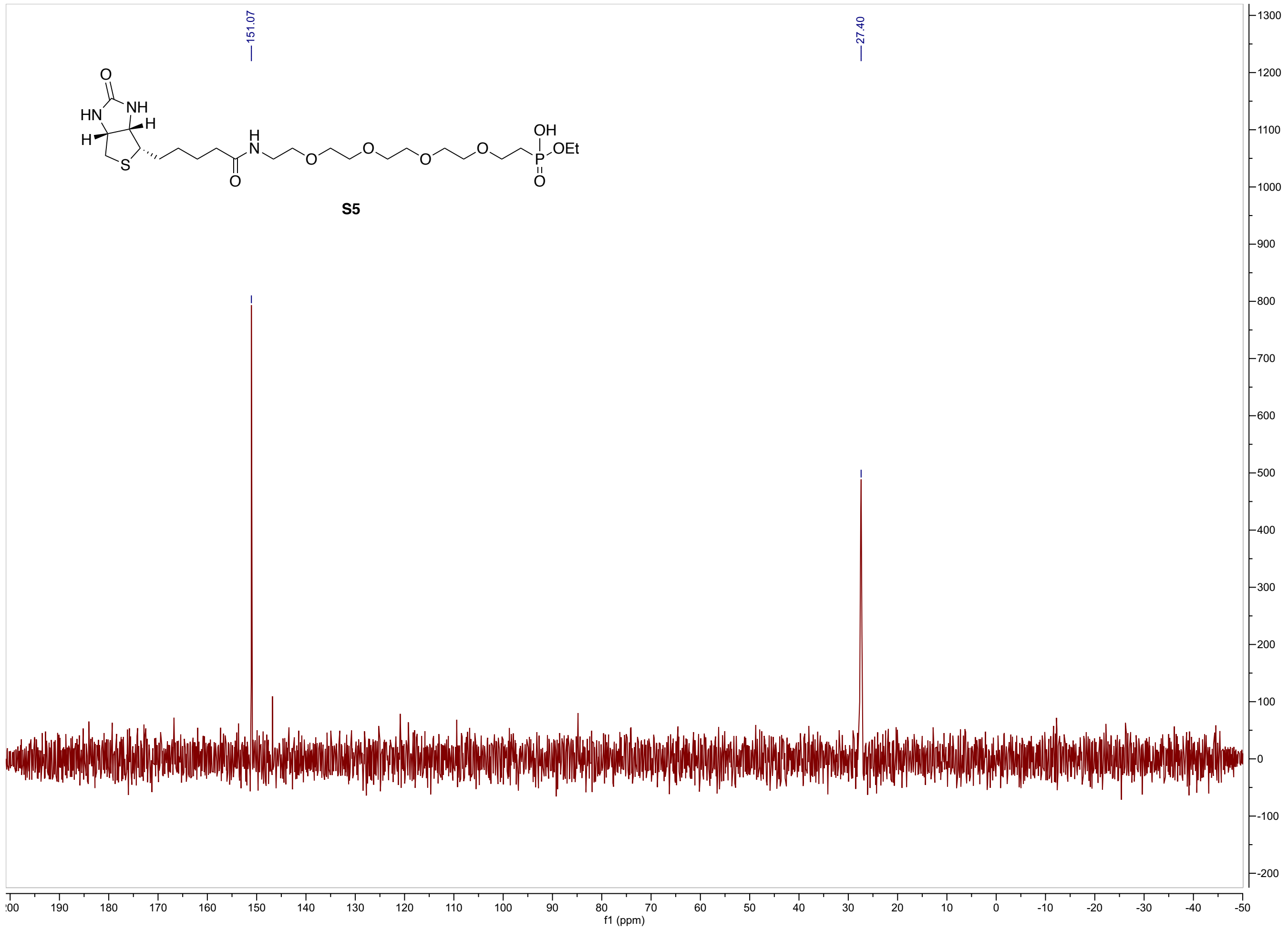


**S5** $^{31}P$

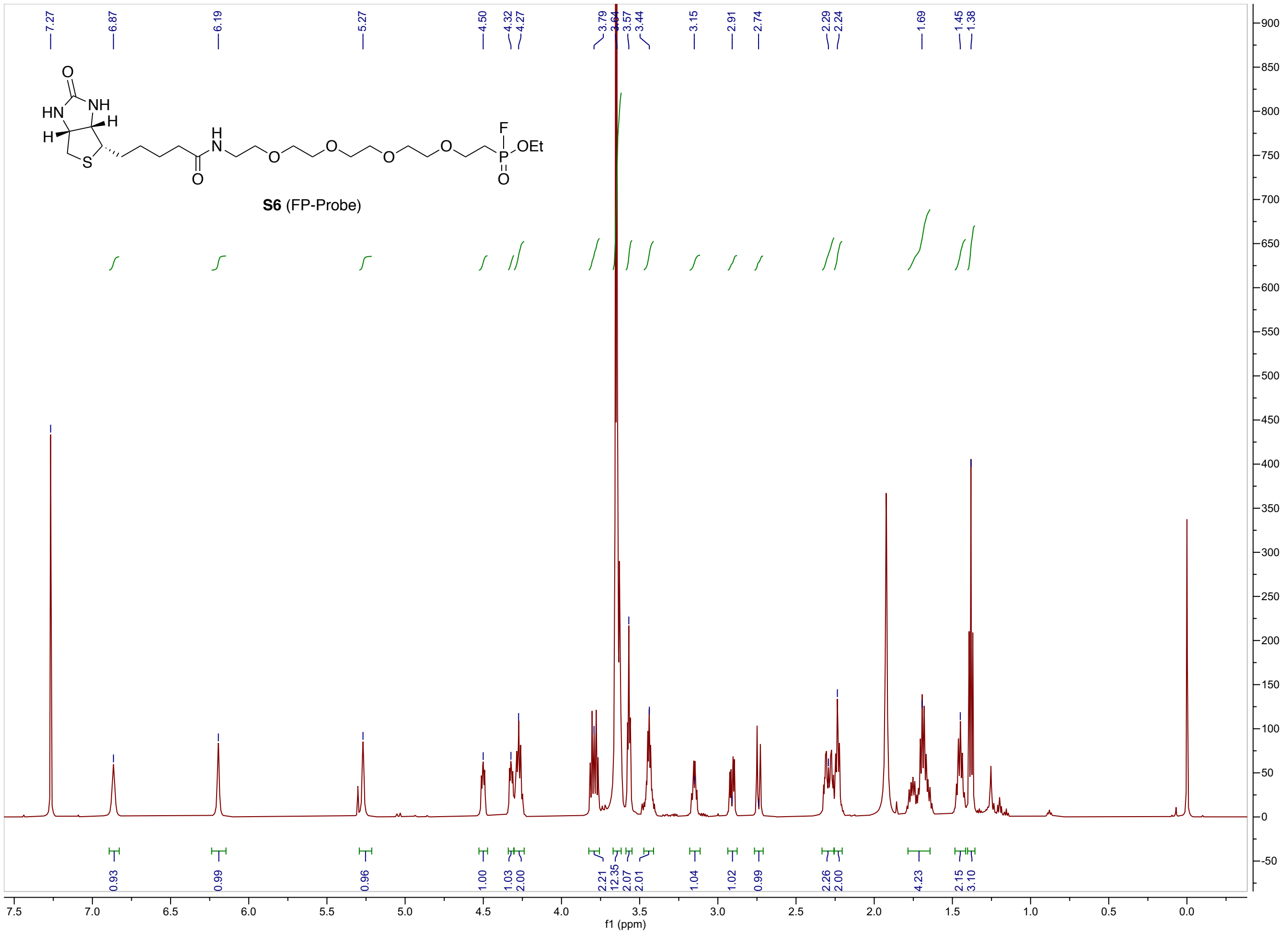
S6 (FP-Probe)
f1 (ppm)
7.27
6.87
6.19
5.27
4.50
4.32
4.27
3.79
3.64
3.57
3.44
3.15
2.91
2.74
2.29
2.24
1.69
1.45
1.38
0.93
0.99
0.96
1.00
1.03
2.00
2.21
12.35
2.07
2.01
1.04
1.02
0.99
2.26
2.00
4.23
2.15
3.10

**S6** ¹H

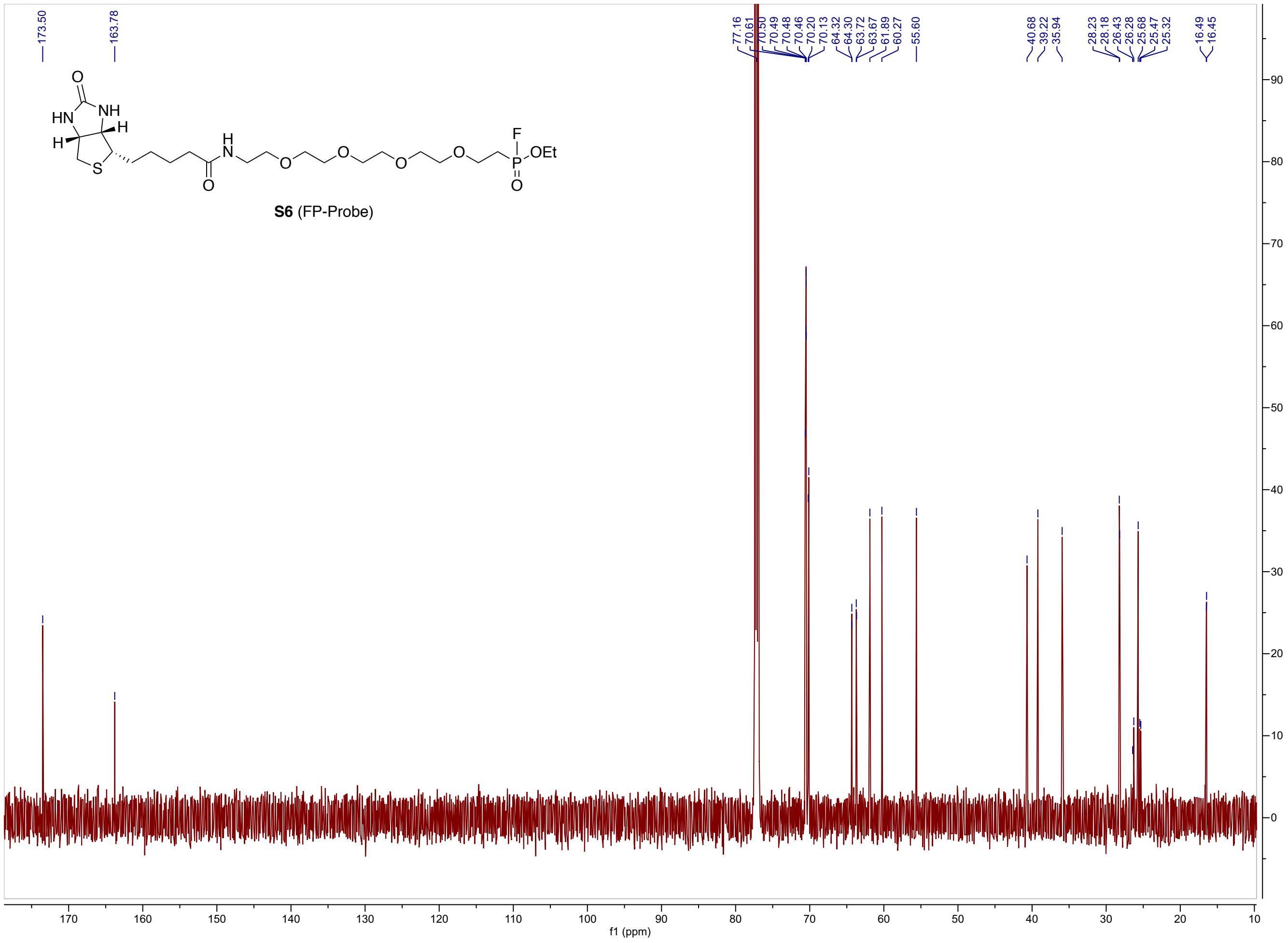
173.50
163.78
77.16
70.61
70.50
70.49
70.48
70.46
70.20
70.13
64.32
64.30
63.72
63.67
61.89
60.27
55.60
40.68
39.22
35.94
28.23
28.18
26.43
26.28
25.68
25.47
25.32
16.49
16.45
S6 (FP-Probe)
f1 (ppm)

**S6** $^{13}C$
